## Supplementary Figures & Appendix for "Ancient Trans-Species Polymorphism at the Major Histocompatibility Complex in Primates": MHC_Paper_2_eLife_Revisions_supplementary.pdf

### Appendix 1

#### MHC Nomenclature

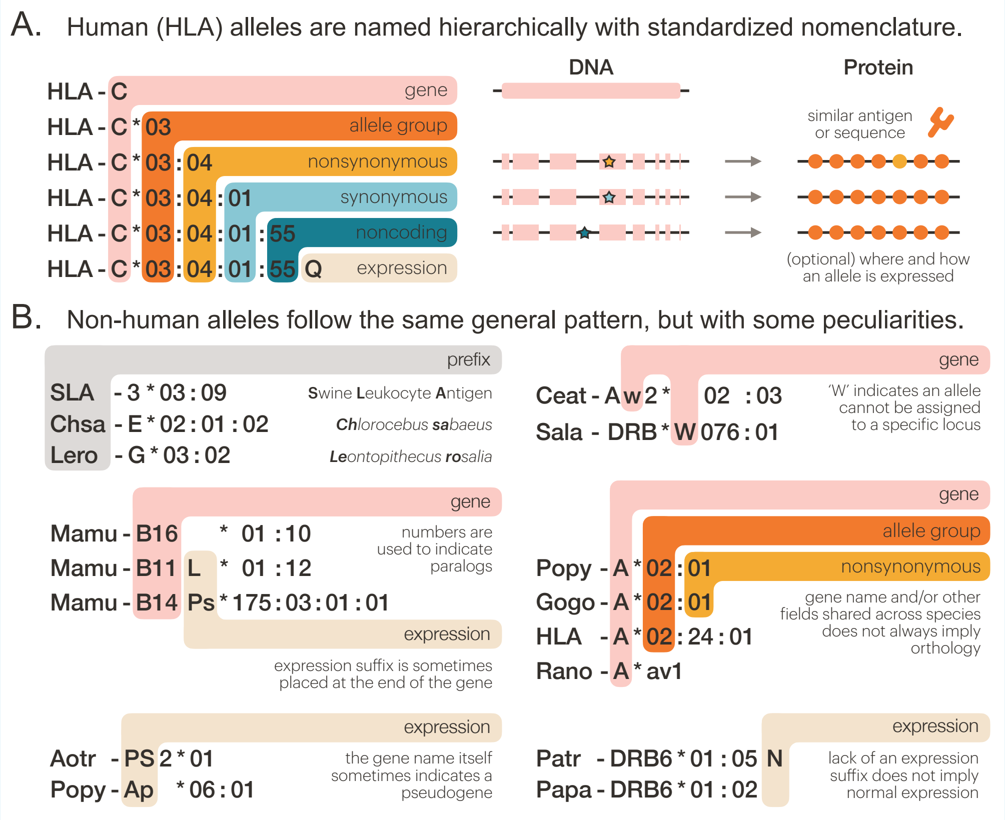


**Appendix 1—figure 1. MHC allele nomenclature. A)** Human HLA alleles are named in a standard fashion, with the gene name followed by four colon-separated fields. The first field indicates a broad-scale allele group which sometimes corresponds to a serological antigen. The second field denotes a specific HLA protein. The third field indicates synonymous changes to the nucleotide sequence in the coding region, while the fourth field is used to distinguish alleles with differences in the noncoding regions. If an allele's expression has been characterized, an informative suffix is sometimes added (***Robinson et al., 2024***; ***Marsh et al., 2010***). **B)** Researchers have applied the same format to non-human alleles, with some key differences. Instead of "HLA", a prefix which concatenates the first two letters of the genus name with the first two letters of the species name is used, except in certain cases where the species' MHC system was named long ago. Paralogs can be distinguished using numbers, but sequences unassigned to a particular locus or paralog might incorporate a 'W' in the gene name. Use of expression tags varies, with some being added to the end of the gene name instead of the end of the entire allele name. Pseudogenes can be denoted with gene name suffixes, gene names themselves, expression suffixes, or not at all. For both human and non-human alleles, the lack of an expression suffix does not imply normal expression (***de Groot et al., 2020***). SLA: Swine Leukocyte Antigen; Chsa: *Chlorocebus sabaeus*—green monkey; Lero: *Leontopithecus rosalia*—golden lion tamarin; Mamu: *Macaca mulatta*—rhesus macaque; Aotr: *Aotus trivirgatus*—three-striped night monkey; Popy: *Pongo pygmaeus*—Bornean orangutan; Ceat: *Cercocebus atys*—sooty mangabey; Sala: *Saguinus labiatus*—white-lipped tamarin; Gogo: *Gorilla gorilla*—Western gorilla; Rano: *Rattus norvegicus*—brown rat; Patr: *Pan troglodytes*—chimpanzee; Papa: *Pan paniscus*—bonobo.

The large number of genes, some with thousands of alleles, necessitates a consistent naming scheme. Known alleles are given names such as "Aole-DQB1*23:01", and names are maintained and updated by the WHO Nomenclature Committee for Factors of the HLA System (***Robinson et al., 2024***; ***Marsh et al., 2010***). First, the species of origin is indicated by a four-letter prefix consisting of the first two letters of the genus name and the first two letters of the species name, e.g. "Chsa-" for *Chlorocebus sabaeus*, the green monkey. There are some exceptions, usually because these MHC systems were first investigated before the naming scheme was put into place. These include "HLA-" for human, "H2-" for mouse, "RT1-" for rat, and "SLA-" for swine, among others (***de Groot et al., 2020***; ***De Groot et al., 2012***).

After the hyphen is the locus designation. Some species, such as human, have a relatively simple landscape of MHC genes, making it easy to identify sequences that belong to a particular gene. However, other species have recent gene expansions and considerable region conformation diversity, making it difficult to assign alleles to genes. In some cases, these are given generic locus designations; for example, rhesus macaques have at least 19 paralogous B loci but most are given the ambiguous name "Mamu-B", with the exception of a few well-characterized genes such as Mamu-B17. In other cases, unassigned sequences are given a working designation indicated by a "W", such as "Popy-DRB*W113:01". In this example, the allele definitely belongs to a DRB paralog, but it is unclear which one. Some locus names are given a "Ps" suffix to indicate they are pseudogenes, such as "Caja-G5Ps". However, not all pseudogenes are labeled this way, so one should not assume the lack of a "Ps" suffix means a gene is functional (***de Groot et al., 2020***; ***De Groot et al., 2012***).

After the species and locus name, each MHC allele is designated by up to four fields separated by colons. The first field designates the type or family. Types often, but not always, correspond to the broad serological reactivity of the allele, as many were named before full sequences were known. To facilitate comparison across closely-related species, researchers generally try to give related MHC alleles the same first-field designation, e.g. Gogo-A*02 and HLA-A*02. However, certain genes do not follow this general rule. For example, MHC-DPB1 has undergone considerable gene conversion, resulting in no distinct types; thus, a shared first-field designation between species is meaningless for this gene (***De Groot et al., 2012***; ***de Groot et al., 2020***). The second field designates the allele subtype, or unique amino acid sequence. For example, "Patr-A*08:01" and "Patr-A*08:02" are part of the same allelic family, but have some nonsynonymous differences. Synonymous changes are specified by the third field. For example, "Paan-DPB1*03:01:01" and"Paan-DPB1*03:01:02" have silent substitutions which ultimately result in the same protein. Lastly, the fourth field is used to describe changes to the noncoding regions—that is, the 5’ and 3’ UTRs and the introns. Of course, this requires that these regions have been sequenced, so not all alleles will have a fourth field. Finally, alleles can also be followed by an optional suffix to describe expression changes, most commonly "N" for a null/nonexpressed allele or "L" for a lowly-expressed allele (***Hurley, 2021***; ***Douillard et al., 2021***).

In general, caution must be taken in interpreting allele names. First, because not all alleles are resolved at three- and four-field resolution, the names are not all strictly hierarchical; alleles which have all four fields cannot always simply be truncated to obtain the two-field version. Second, because human alleles were named in order of discovery, alleles with very different one- or two-field designations could ultimately have the same nucleotide or amino acid sequence in the peptide-binding groove. When discussing functional consequences, it is relevant to group alleles by their nucleotide or amino acid sequence in the PBR (designated G- and P-groups, respectively) and not necessarily by their one- or two-field name (***Hurley, 2021***; ***Douillard et al., 2021***). Additionally, the suffixes can be misleading because not every allele has had its expression level characterized—the absence of an "L" does not mean that an allele has normal expression (***Hurley, 2021***). Despite these small issues, the naming system is generally intuitive and very useful for understanding alleles at a glance. In this work, alleles obtained from the IPD-MHC and IPD-IMGT/HLA databases are named this way, but sequences obtained from RefSeq are labeled by accession number or location in a genome.

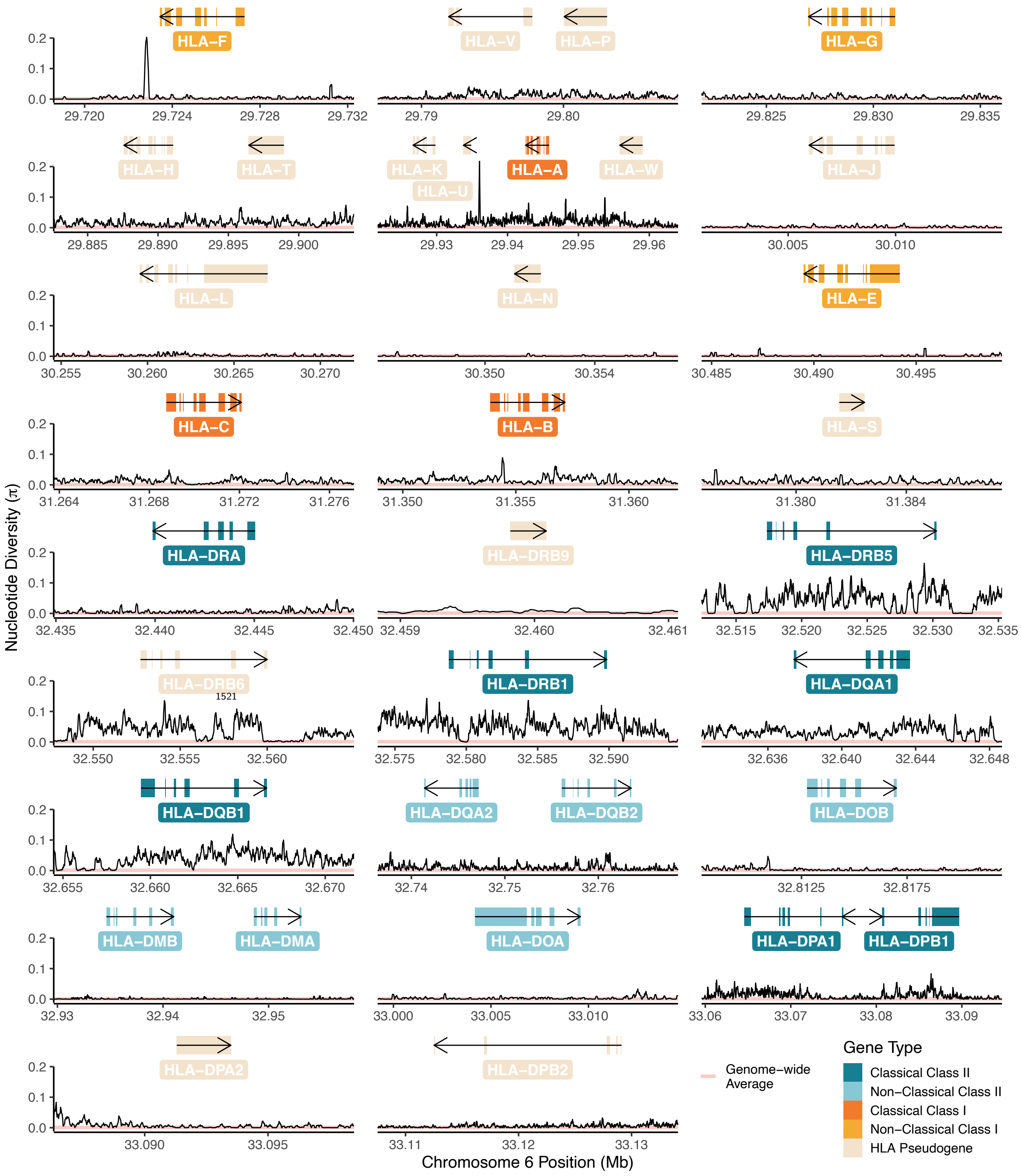


**Figure 1—figure supplement 1. Nucleotide Diversity in the Human HLA Region.** Nucleotide diversity (Nei and Li's $\pi$) around each HLA gene is shown in black, while the pink horizontal line shows the genome-wide average nucleotide diversity ($\pi \approx 0.001$) (***Sachidanandam et al., 2001***). The genes are shown in order along the genome (from top left to bottom right), but the x-axis is repeatedly broken in order to zoom in on detail around each gene. The genes are colored according to their type, with key shown at bottom right. For the functional genes (and some pseudogenes), the exon structure is shown by boxes; other pseudogenes are not annotated with this level of detail.

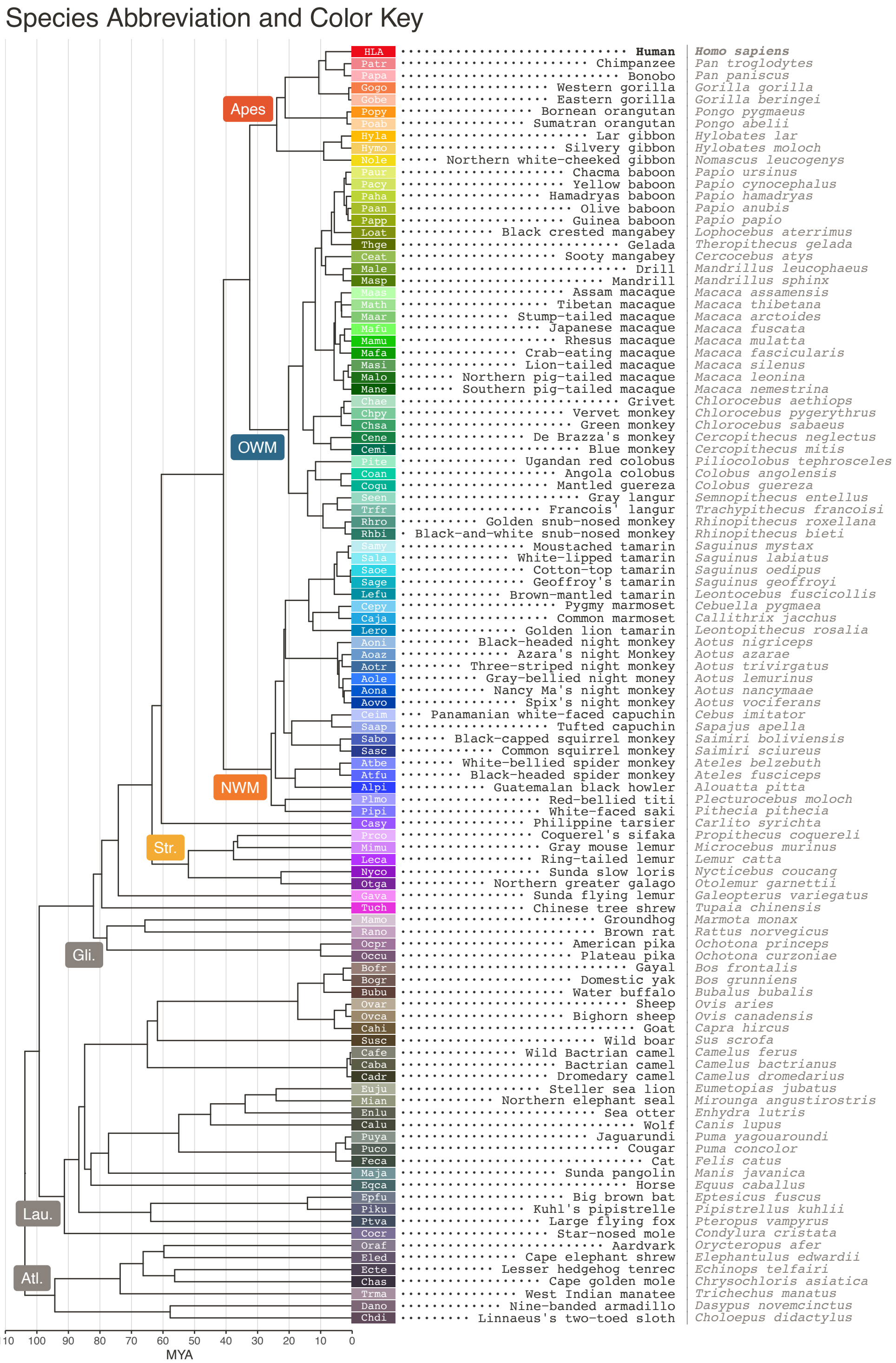


**Figure 1—figure supplement 2. Species Key.** On the left hand side is the species tree relating the species used in this study (***Kuderna et al., 2023***; ***Foley et al., 2023***). Each species/tip is labeled with a unique color and 4-letter abbreviation, which is composed of the first two letters of the genus name and first two letters of the species name. The common name and Latin name for each species is shown on the right hand side. *Plecturocebus moloch* is listed in the IPD-MHC database under its old name, *Callicebus moloch*, and uses a different abbreviation (Camo) in that resource. Similarly, *Leontocebus fuscicolis* was formerly known as *Saguinus fuscicollis* (Safu) in the IPD-MHC database. There appears to be some debate as to whether the pygmy marmoset should be placed in the *Callithrix* or *Cebuella* genus, but we have used the name *Cebuella pygmaea* (Cepy) in accordance with a recent primate study (***Kuderna et al., 2023***). This species is known as *Callithrix pygmaea* (Capy) in IPD-MHC. OWM, Old-World Monkeys; NWM, New World Monkeys; Str., *Strepsirrhini*; Gli., *Glires*; Lau., *Laurasiatheria*; Atl., *Atlantogenata*.

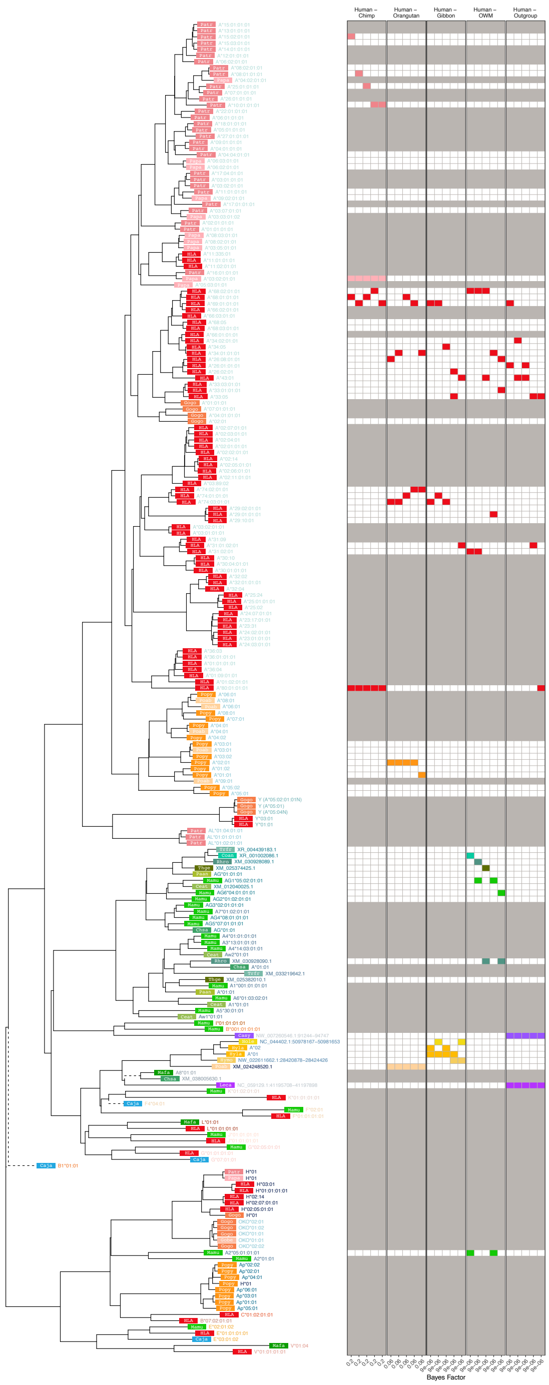


**Figure 2—figure supplement 1. MHC-A-Related Group *BEAST2* tree for exon 2 (PBR-encoding).** In the tree, each tip represents a sequence (see ***Appendix 1*** for more details on nomenclature), with the colored rectangle and four-letter abbreviation indicating the species (see ***Figure 1—figure Supplement 2*** for full species key). Following the rectangle, tips are labeled with the sequence name; sequences which have been assigned to loci are colored according to the gene group, while unassigned sequences are written in gray. Dashed branches are shortened to 10% of their length to expand detail in the rest of the tree. The left-hand side shows example Bayes factors we calculated from the set of posterior trees; the tree tips correspond to the rows of the grid. Each panel represents a type of comparison (labeled at top) and each column represents one of the top 5 highest-Bayes-factor comparisons (exact value at bottom; $> 100$ indicates strong evidence for TSP). The 4 colored blocks in each column include two red blocks (human sequences) that were tested against two sequences from other species (see Methods). Grayed-out rows correspond to sequences that were not considered for Bayes factors, either because they belong to a non-orthologous gene or backbone sequence (and thus not relevant to compare with the human gene in question), or because they may have been involved in a gene conversion event in this exon (according to *GENECONV* or from the literature).

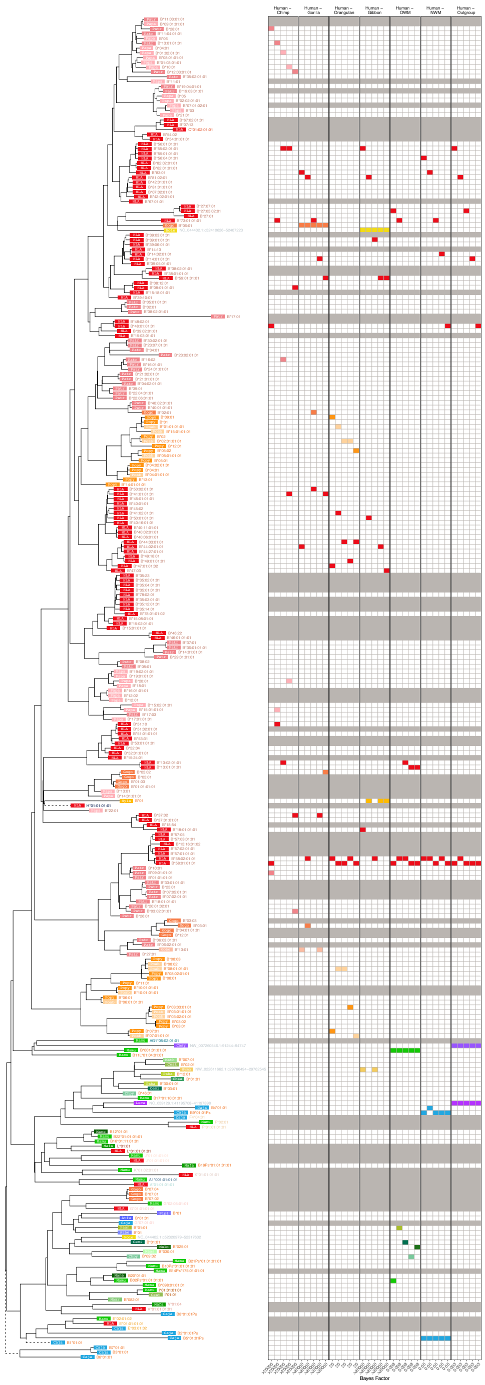


**Figure 2—figure supplement 2. MHC-B-Related Group *BEAST2* tree for exon 2 (PBR-encoding).** In the tree, each tip represents a sequence (see ***Appendix 1*** for more details on nomenclature), with the colored rectangle and four-letter abbreviation indicating the species (see ***Figure 1—figure Supplement 2*** for full species key). Following the rectangle, tips are labeled with the sequence name; sequences which have been assigned to loci are colored according to the gene group, while unassigned sequences are written in gray. Dashed branches are shortened to 10% of their length to expand detail in the rest of the tree. The left-hand side shows example Bayes factors we calculated from the set of posterior trees; the tree tips correspond to the rows of the grid. Each panel represents a type of comparison (labeled at top) and each column represents one of the top 5 highest-Bayes-factor comparisons (exact value at bottom; $> 100$ indicates strong evidence for TSP). The 4 colored blocks in each column include two red blocks (human sequences) that were tested against two sequences from other species (see Methods). Grayed-out rows correspond to sequences that were not considered for Bayes factors, either because they belong to a non-orthologous gene or backbone sequence (and thus not relevant to compare with the human gene in question), or because they may have been involved in a gene conversion event in this exon (according to *GENECONV* or from the literature).

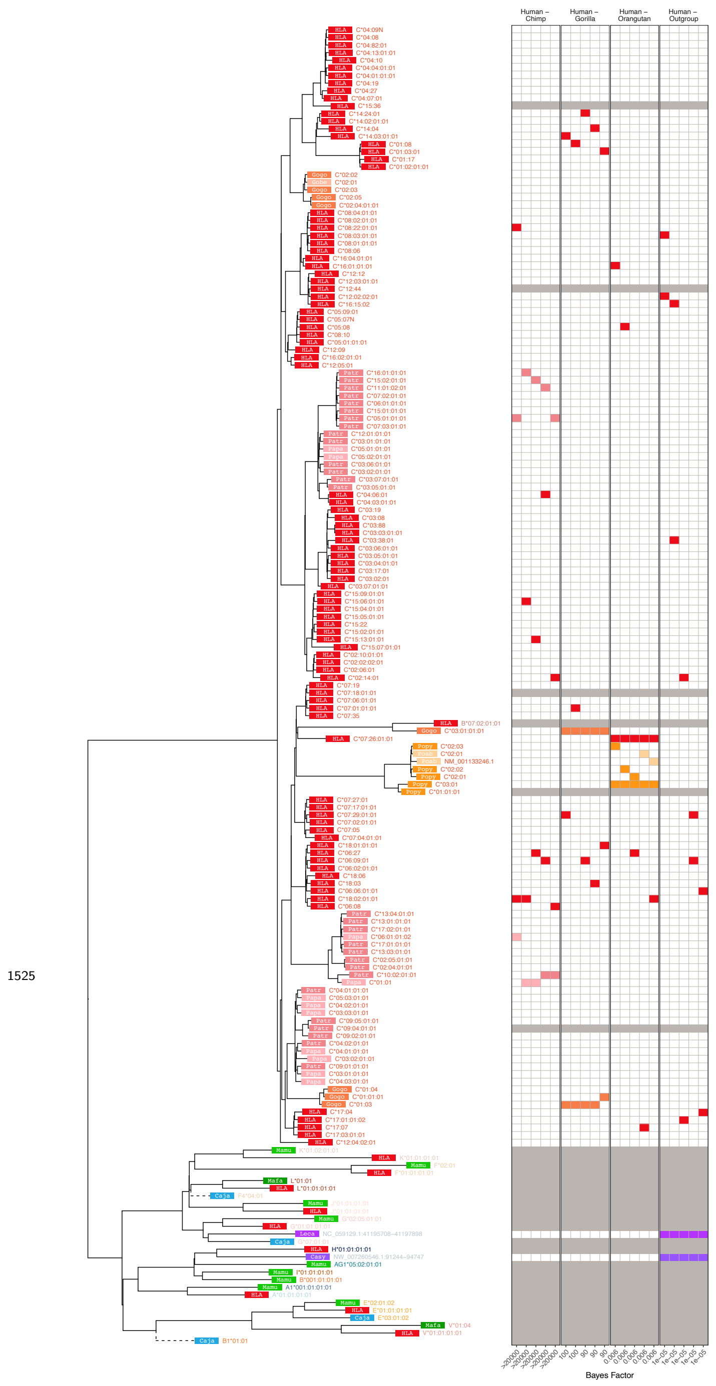


**Figure 2—figure supplement 3. MHC-C-Related Group *BEAST2* tree for exon 2 (PBR-encoding).** In the tree, each tip represents a sequence (see ***Appendix 1*** for more details on nomenclature), with the colored rectangle and four-letter abbreviation indicating the species (see ***Figure 1—figure Supplement 2*** for full species key). Following the rectangle, tips are labeled with the sequence name; sequences which have been assigned to loci are colored according to the gene group, while unassigned sequences are written in gray. Dashed branches are shortened to 10% of their length to expand detail in the rest of the tree. The left-hand side shows example Bayes factors we calculated from the set of posterior trees; the tree tips correspond to the rows of the grid. Each panel represents a type of comparison (labeled at top) and each column represents one of the top 5 highest-Bayes-factor comparisons (exact value at bottom; $> 100$ indicates strong evidence for TSP). The 4 colored blocks in each column include two red blocks (human sequences) that were tested against two sequences from other species (see Methods). Grayed-out rows correspond to sequences that were not considered for Bayes factors, either because they belong to a non-orthologous gene or backbone sequence (and thus not relevant to compare with the human gene in question), or because they may have been involved in a gene conversion event in this exon (according to *GENECONV* or from the literature).

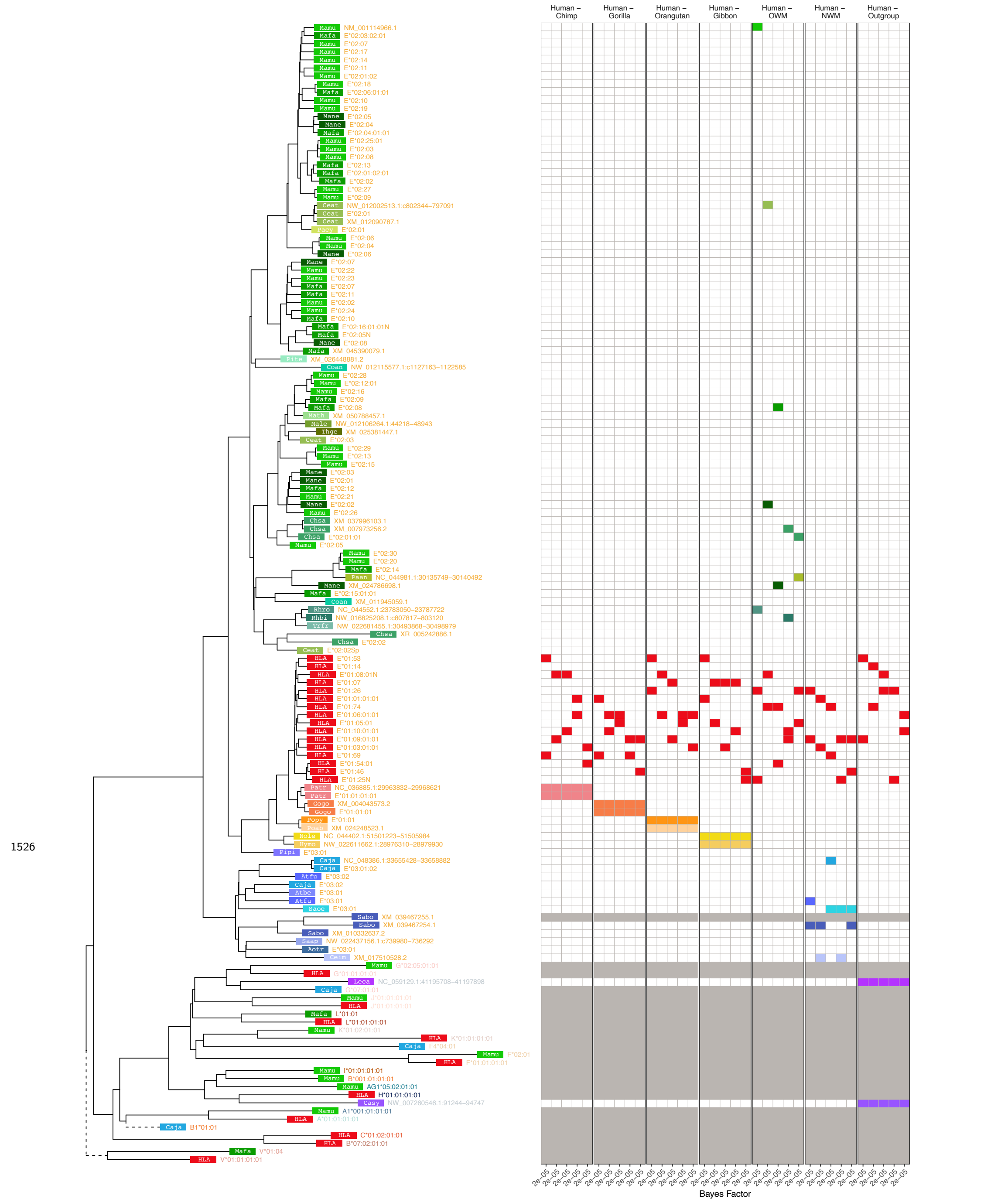


**Figure 2—figure supplement 4. MHC-E-Related Group *BEAST2* tree for exon 2 (PBR-encoding).** In the tree, each tip represents a sequence (see ***Appendix 1*** for more details on nomenclature), with the colored rectangle and four-letter abbreviation indicating the species (see ***Figure 1—figure Supplement 2*** for full species key). Following the rectangle, tips are labeled with the sequence name; sequences which have been assigned to loci are colored according to the gene group, while unassigned sequences are written in gray. Dashed branches are shortened to 10% of their length to expand detail in the rest of the tree. The left-hand side shows example Bayes factors we calculated from the set of posterior trees; the tree tips correspond to the rows of the grid. Each panel represents a type of comparison (labeled at top) and each column represents one of the top 5 highest-Bayes-factor comparisons (exact value at bottom; $> 100$ indicates strong evidence for TSP). The 4 colored blocks in each column include two red blocks (human sequences) that were tested against two sequences from other species (see Methods). Grayed-out rows correspond to sequences that were not considered for Bayes factors, either because they belong to a non-orthologous gene or backbone sequence (and thus not relevant to compare with the human gene in question), or because they may have been involved in a gene conversion event in this exon (according to *GENECONV* or from the literature).

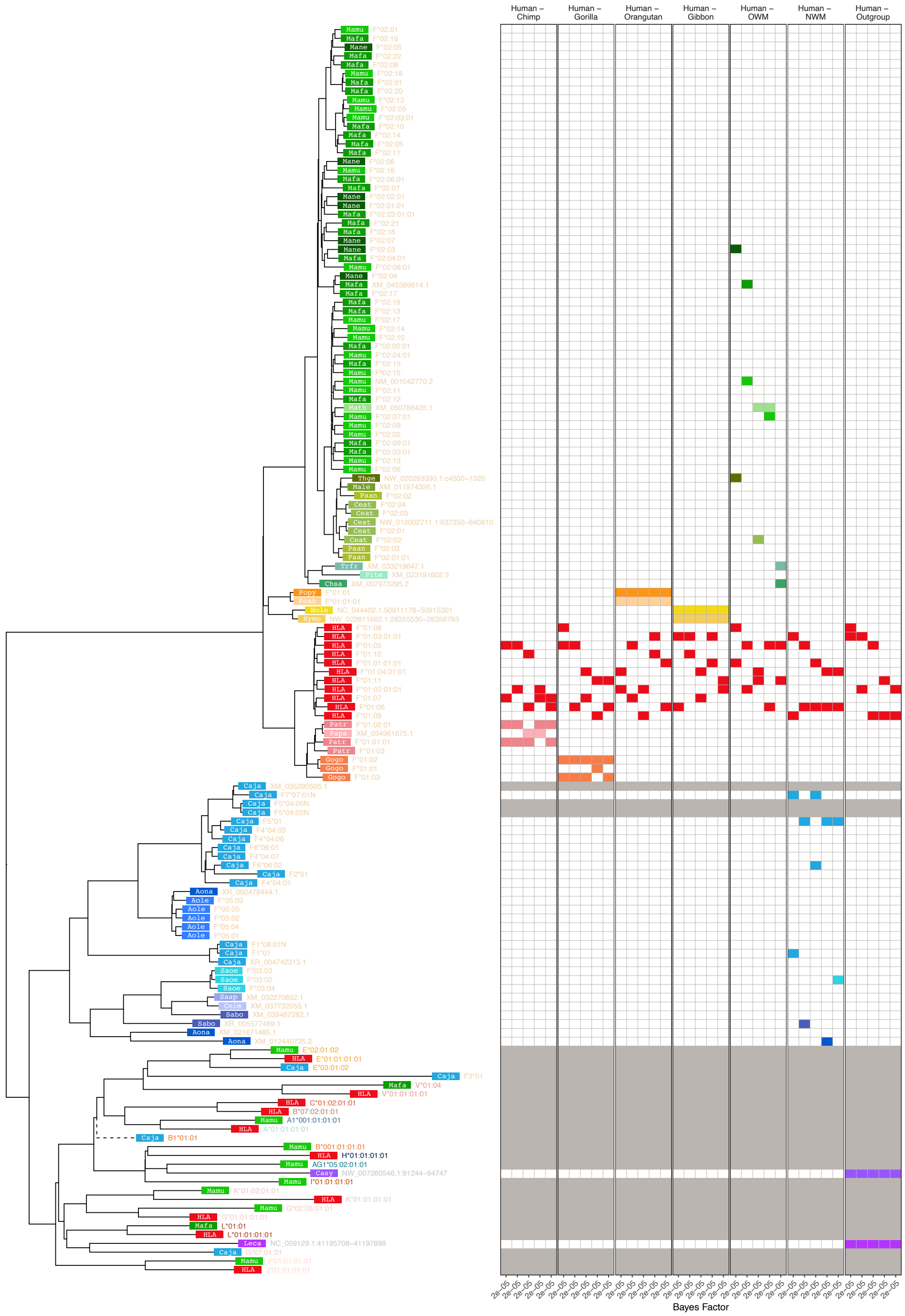


**Figure 2—figure supplement 5. MHC-F-Related Group *BEAST2* tree for exon 2 (PBR-encoding).** In the tree, each tip represents a sequence (see ***Appendix 1*** for more details on nomenclature), with the colored rectangle and four-letter abbreviation indicating the species (see ***Figure 1—figure Supplement 2*** for full species key). Following the rectangle, tips are labeled with the sequence name; sequences which have been assigned to loci are colored according to the gene group, while unassigned sequences are written in gray. Dashed branches are shortened to 10% of their length to expand detail in the rest of the tree. The left-hand side shows example Bayes factors we calculated from the set of posterior trees; the tree tips correspond to the rows of the grid. Each panel represents a type of comparison (labeled at top) and each column represents one of the top 5 highest-Bayes-factor comparisons (exact value at bottom; $> 100$ indicates strong evidence for TSP). The 4 colored blocks in each column include two red blocks (human sequences) that were tested against two sequences from other species (see Methods). Grayed-out rows correspond to sequences that were not considered for Bayes factors, either because they belong to a non-orthologous gene or backbone sequence (and thus not relevant to compare with the human gene in question), or because they may have been involved in a gene conversion event in this exon (according to *GENECONV* or from the literature).

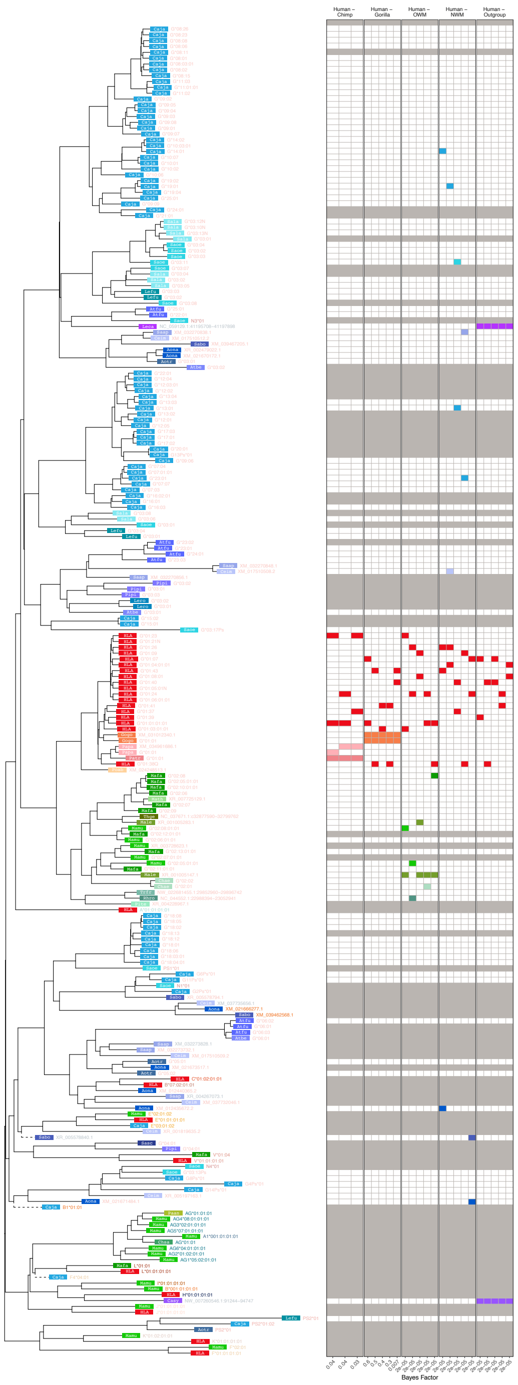


**Figure 2—figure supplement 6. MHC-G-Related Group *BEAST2* tree for exon 2 (PBR-encoding).** In the tree, each tip represents a sequence (see ***Appendix 1*** for more details on nomenclature), with the colored rectangle and four-letter abbreviation indicating the species (see ***Figure 1—figure Supplement 2*** for full species key). Following the rectangle, tips are labeled with the sequence name; sequences which have been assigned to loci are colored according to the gene group, while unassigned sequences are written in gray. Dashed branches are shortened to 10% of their length to expand detail in the rest of the tree. The left-hand side shows example Bayes factors we calculated from the set of posterior trees; the tree tips correspond to the rows of the grid. Each panel represents a type of comparison (labeled at top) and each column represents one of the top 5 highest-Bayes-factor comparisons (exact value at bottom; $> 100$ indicates strong evidence for TSP). The 4 colored blocks in each column include two red blocks (human sequences) that were tested against two sequences from other species (see Methods). Grayed-out rows correspond to sequences that were not considered for Bayes factors, either because they belong to a non-orthologous gene or backbone sequence (and thus not relevant to compare with the human gene in question), or because they may have been involved in a gene conversion event in this exon (according to *GENECONV* or from the literature).

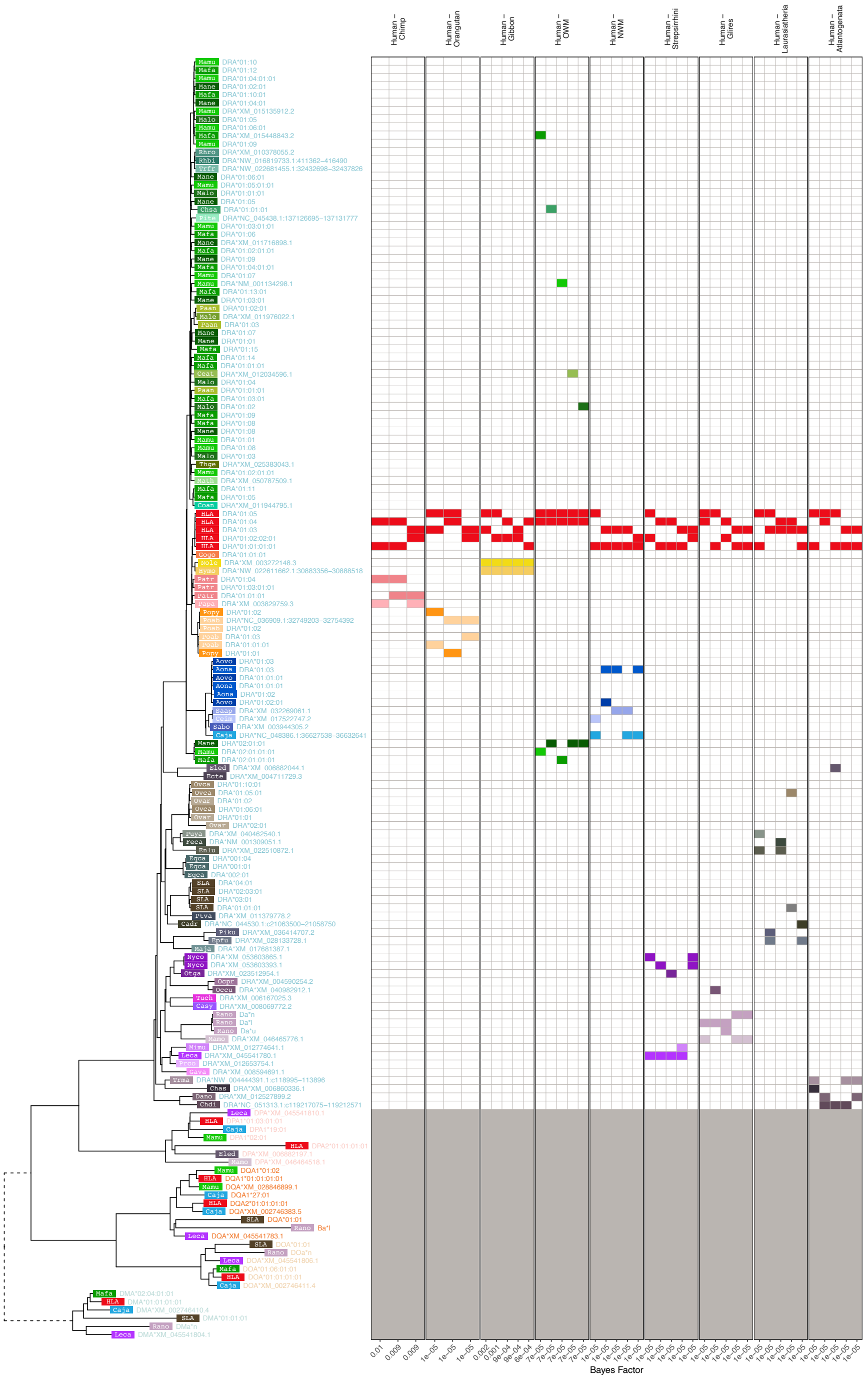


**Figure 2—figure supplement 7. MHC-DRA-Related Group *BEAST2* tree for exon 2 (PBR-encoding).** In the tree, each tip represents a sequence (see ***Appendix 1*** for more details on nomenclature), with the colored rectangle and four-letter abbreviation indicating the species (see ***Figure 1—figure Supplement 2*** for full species key). Following the rectangle, tips are labeled with the sequence name; sequences which have been assigned to loci are colored according to the gene group, while unassigned sequences are written in gray. Dashed branches are shortened to 10% of their length to expand detail in the rest of the tree. The left-hand side shows example Bayes factors we calculated from the set of posterior trees; the tree tips correspond to the rows of the grid. Each panel represents a type of comparison (labeled at top) and each column represents one of the top 5 highest-Bayes-factor comparisons (exact value at bottom; $> 100$ indicates strong evidence for TSP). The 4 colored blocks in each column include two red blocks (human sequences) that were tested against two sequences from other species (see Methods). Grayed-out rows correspond to sequences that were not considered for Bayes factors, either because they belong to a non-orthologous gene or backbone sequence (and thus not relevant to compare with the human gene in question), or because they may have been involved in a gene conversion event in this exon (according to *GENECONV* or from the literature).

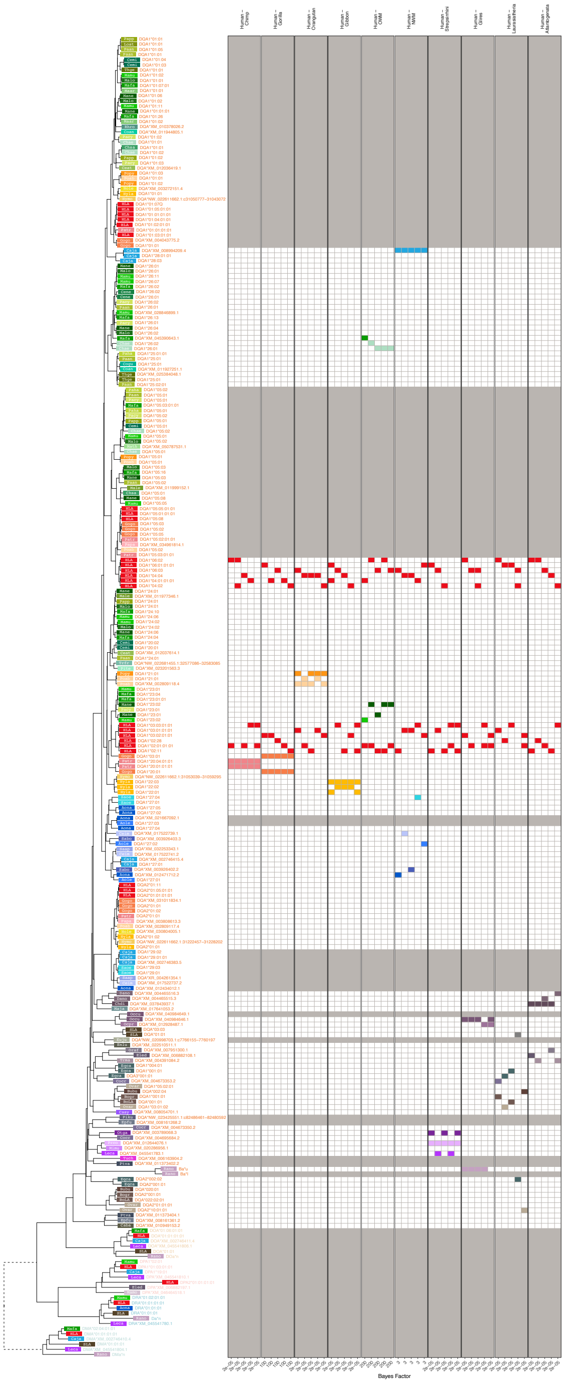


**Figure 2—figure supplement 8. MHC-DQA-Related Group *BEAST2* tree for exon 2 (PBR-encoding).** In the tree, each tip represents a sequence (see ***Appendix 1*** for more details on nomenclature), with the colored rectangle and four-letter abbreviation indicating the species (see ***Figure 1—figure Supplement 2*** for full species key). Following the rectangle, tips are labeled with the sequence name; sequences which have been assigned to loci are colored according to the gene group, while unassigned sequences are written in gray. Dashed branches are shortened to 10% of their length to expand detail in the rest of the tree. The left-hand side shows example Bayes factors we calculated from the set of posterior trees; the tree tips correspond to the rows of the grid. Each panel represents a type of comparison (labeled at top) and each column represents one of the top 5 highest-Bayes-factor comparisons (exact value at bottom; $> 100$ indicates strong evidence for TSP). The 4 colored blocks in each column include two red blocks (human sequences) that were tested against two sequences from other species (see Methods). Grayed-out rows correspond to sequences that were not considered for Bayes factors, either because they belong to a non-orthologous gene or backbone sequence (and thus not relevant to compare with the human gene in question), or because they may have been involved in a gene conversion event in this exon (according to *GENECONV* or from the literature).

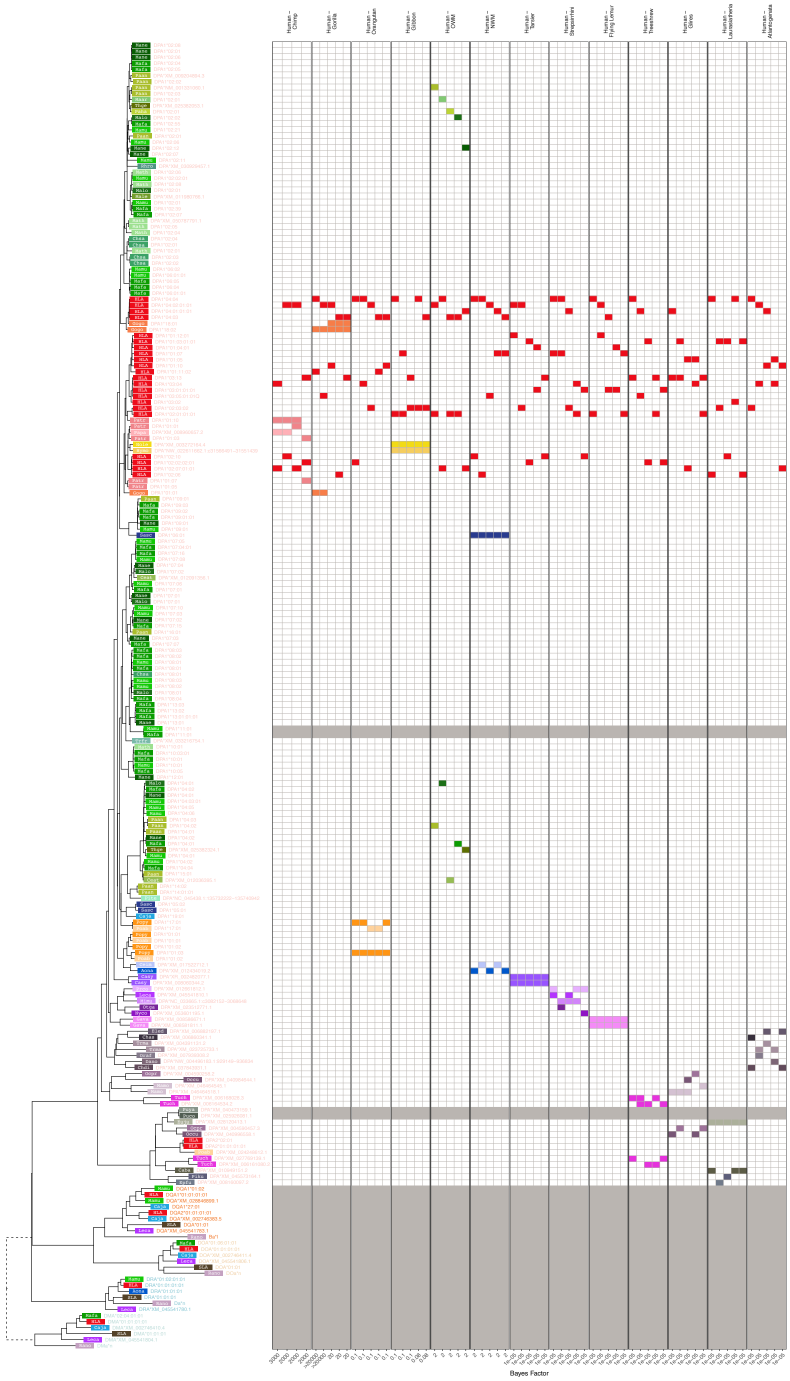


**Figure 2—figure supplement 9. MHC-DPA-Related Group *BEAST2* tree for exon 2 (PBR-encoding).** In the tree, each tip represents a sequence (see ***Appendix 1*** for more details on nomenclature), with the colored rectangle and four-letter abbreviation indicating the species (see ***Figure 1—figure Supplement 2*** for full species key). Following the rectangle, tips are labeled with the sequence name; sequences which have been assigned to loci are colored according to the gene group, while unassigned sequences are written in gray. Dashed branches are shortened to 10% of their length to expand detail in the rest of the tree. The left-hand side shows example Bayes factors we calculated from the set of posterior trees; the tree tips correspond to the rows of the grid. Each panel represents a type of comparison (labeled at top) and each column represents one of the top 5 highest-Bayes-factor comparisons (exact value at bottom; $> 100$ indicates strong evidence for TSP). The 4 colored blocks in each column include two red blocks (human sequences) that were tested against two sequences from other species (see Methods). Grayed-out rows correspond to sequences that were not considered for Bayes factors, either because they belong to a non-orthologous gene or backbone sequence (and thus not relevant to compare with the human gene in question), or because they may have been involved in a gene conversion event in this exon (according to *GENECONV* or from the literature).

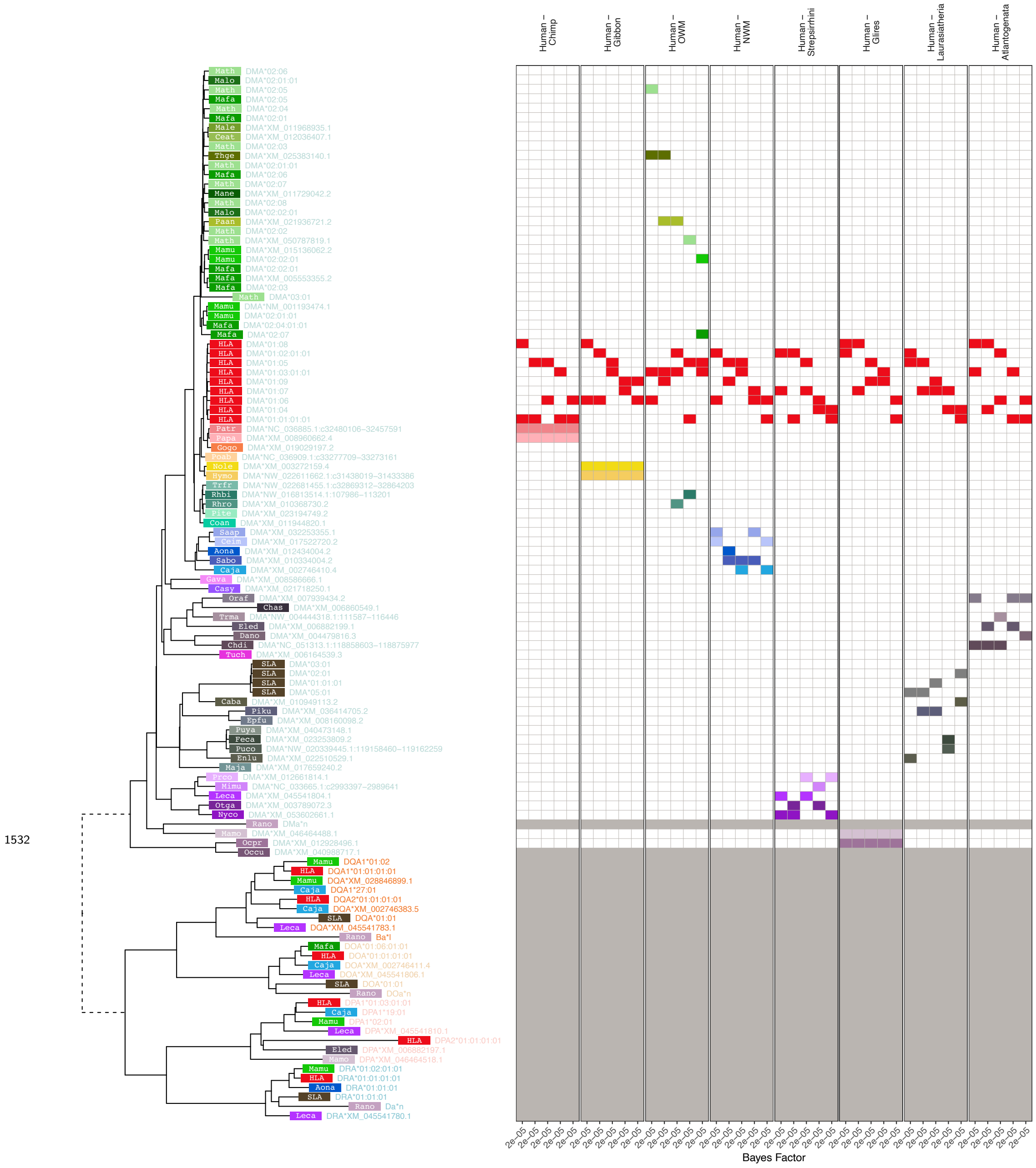


**Figure 2—figure supplement 10. MHC-DMA-Related Group *BEAST2* tree for exon 2 (PBR-encoding).** In the tree, each tip represents a sequence (see ***Appendix 1*** for more details on nomenclature), with the colored rectangle and four-letter abbreviation indicating the species (see ***Figure 1—figure Supplement 2*** for full species key). Following the rectangle, tips are labeled with the sequence name; sequences which have been assigned to loci are colored according to the gene group, while unassigned sequences are written in gray. Dashed branches are shortened to 10% of their length to expand detail in the rest of the tree. The left-hand side shows example Bayes factors we calculated from the set of posterior trees; the tree tips correspond to the rows of the grid. Each panel represents a type of comparison (labeled at top) and each column represents one of the top 5 highest-Bayes-factor comparisons (exact value at bottom; $> 100$ indicates strong evidence for TSP). The 4 colored blocks in each column include two red blocks (human sequences) that were tested against two sequences from other species (see Methods). Grayed-out rows correspond to sequences that were not considered for Bayes factors, either because they belong to a non-orthologous gene or backbone sequence (and thus not relevant to compare with the human gene in question), or because they may have been involved in a gene conversion event in this exon (according to *GENECONV* or from the literature).

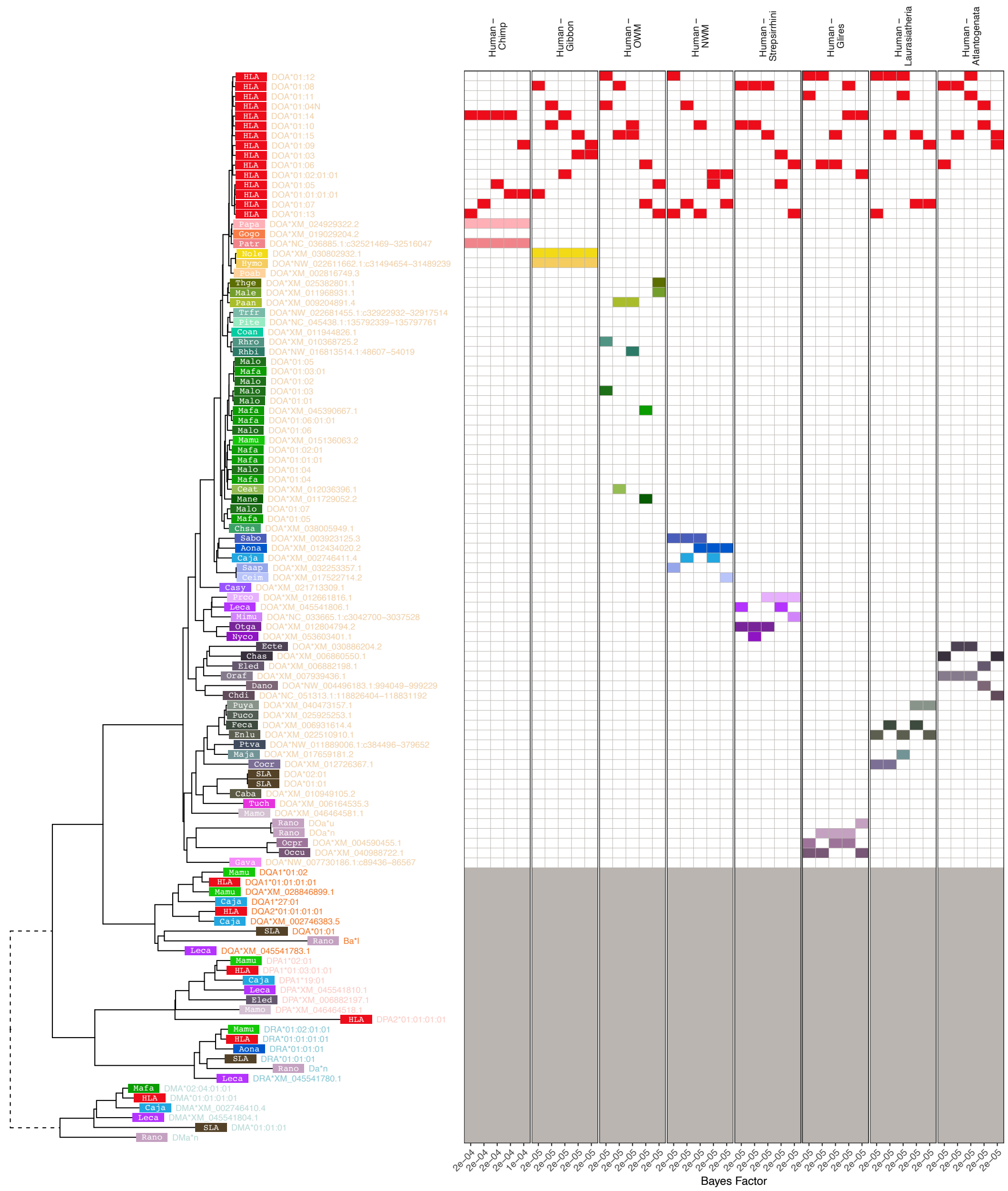


**Figure 2—figure supplement 11. MHC-DOA-Related Group *BEAST2* tree for exon 2 (PBR-encoding).** In the tree, each tip represents a sequence (see ***Appendix 1*** for more details on nomenclature), with the colored rectangle and four-letter abbreviation indicating the species (see ***Figure 1—figure Supplement 2*** for full species key). Following the rectangle, tips are labeled with the sequence name; sequences which have been assigned to loci are colored according to the gene group, while unassigned sequences are written in gray. Dashed branches are shortened to 10% of their length to expand detail in the rest of the tree. The left-hand side shows example Bayes factors we calculated from the set of posterior trees; the tree tips correspond to the rows of the grid. Each panel represents a type of comparison (labeled at top) and each column represents one of the top 5 highest-Bayes-factor comparisons (exact value at bottom; $> 100$ indicates strong evidence for TSP). The 4 colored blocks in each column include two red blocks (human sequences) that were tested against two sequences from other species (see Methods). Grayed-out rows correspond to sequences that were not considered for Bayes factors, either because they belong to a non-orthologous gene or backbone sequence (and thus not relevant to compare with the human gene in question), or because they may have been involved in a gene conversion event in this exon (according to *GENECONV* or from the literature).

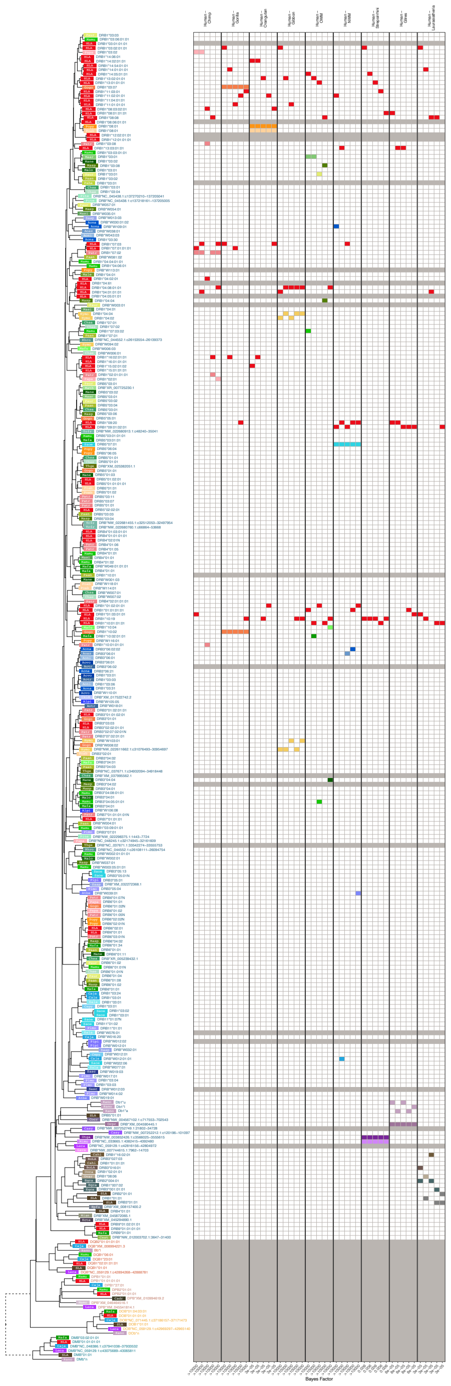

**Figure 2—figure supplement 12. MHC-DRB-Related Group *BEAST2* tree for exon 2 (PBR-encoding).** In the tree, each tip represents a sequence (see ***Appendix 1*** for more details on nomenclature), with the colored rectangle and four-letter abbreviation indicating the species (see ***Figure 1—figure Supplement 2*** for full species key). Following the rectangle, tips are labeled with the sequence name; sequences which have been assigned to loci are colored according to the gene group, while unassigned sequences are written in gray. Dashed branches are shortened to 10% of their length to expand detail in the rest of the tree. The left-hand side shows example Bayes factors we calculated from the set of posterior trees; the tree tips correspond to the rows of the grid. Each panel represents a type of comparison (labeled at top) and each column represents one of the top 5 highest-Bayes-factor comparisons (exact value at bottom; $> 100$ indicates strong evidence for TSP). The 4 colored blocks in each column include two red blocks (human sequences) that were tested against two sequences from other species (see Methods). Grayed-out rows correspond to sequences that were not considered for Bayes factors, either because they belong to a non-orthologous gene or backbone sequence (and thus not relevant to compare with the human gene in question), or because they may have been involved in a gene conversion event in this exon (according to *GENECONV* or from the literature).

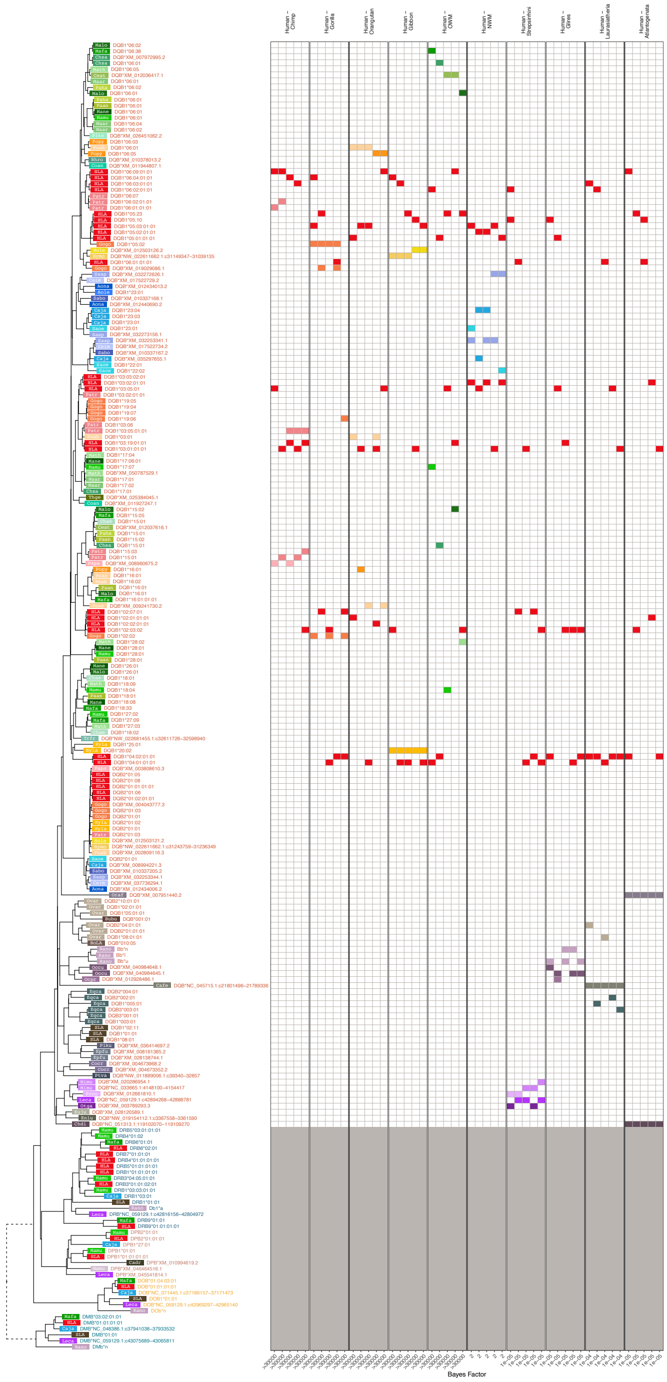


**Figure 2—figure supplement 13. MHC-DQB-Related Group *BEAST2* tree for exon 2 (PBR-encoding).** In the tree, each tip represents a sequence (see ***Appendix 1*** for more details on nomenclature), with the colored rectangle and four-letter abbreviation indicating the species (see ***Figure 1—figure Supplement 2*** for full species key). Following the rectangle, tips are labeled with the sequence name; sequences which have been assigned to loci are colored according to the gene group, while unassigned sequences are written in gray. Dashed branches are shortened to 10% of their length to expand detail in the rest of the tree. The left-hand side shows example Bayes factors we calculated from the set of posterior trees; the tree tips correspond to the rows of the grid. Each panel represents a type of comparison (labeled at top) and each column represents one of the top 5 highest-Bayes-factor comparisons (exact value at bottom; $> 100$ indicates strong evidence for TSP). The 4 colored blocks in each column include two red blocks (human sequences) that were tested against two sequences from other species (see Methods). Grayed-out rows correspond to sequences that were not considered for Bayes factors, either because they belong to a non-orthologous gene or backbone sequence (and thus not relevant to compare with the human gene in question), or because they may have been involved in a gene conversion event in this exon (according to *GENECONV* or from the literature).

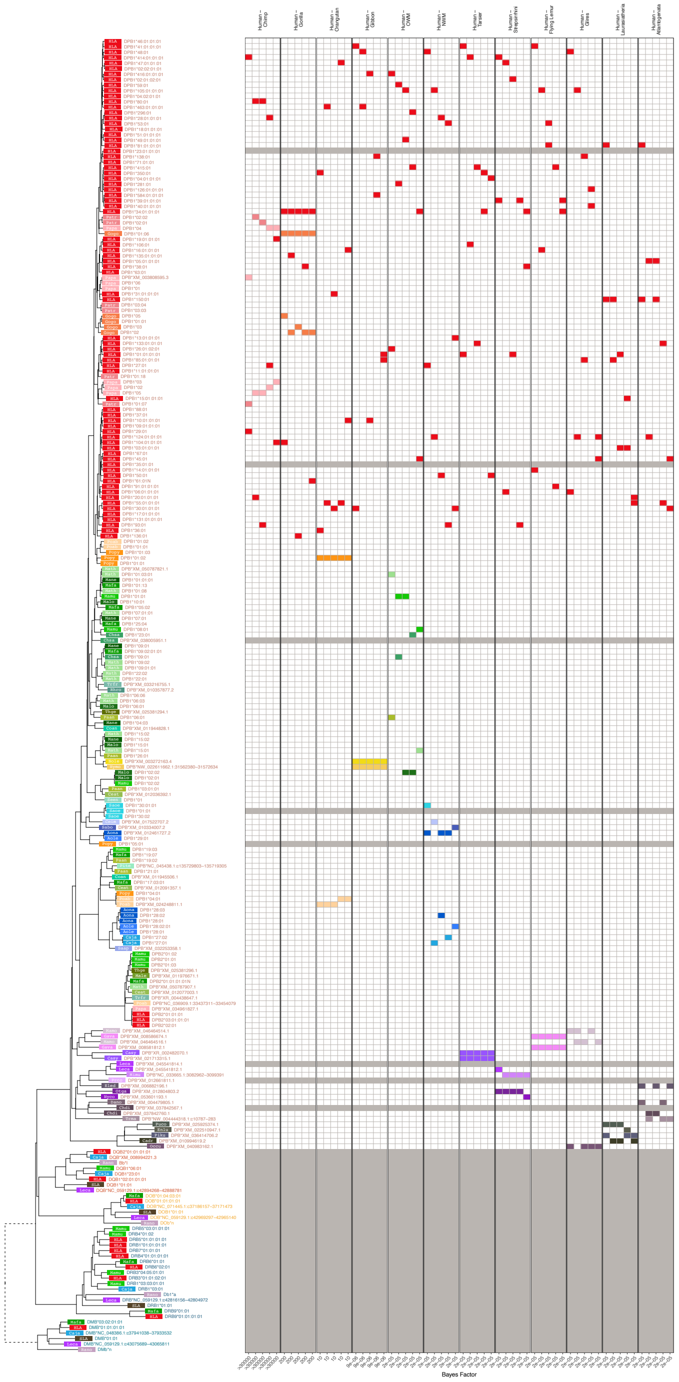


**Figure 2—figure supplement 14. MHC-DPB-Related Group *BEAST2* tree for exon 2 (PBR-encoding).** In the tree, each tip represents a sequence (see ***Appendix 1*** for more details on nomenclature), with the colored rectangle and four-letter abbreviation indicating the species (see ***Figure 1—figure Supplement 2*** for full species key). Following the rectangle, tips are labeled with the sequence name; sequences which have been assigned to loci are colored according to the gene group, while unassigned sequences are written in gray. Dashed branches are shortened to 10% of their length to expand detail in the rest of the tree. The left-hand side shows example Bayes factors we calculated from the set of posterior trees; the tree tips correspond to the rows of the grid. Each panel represents a type of comparison (labeled at top) and each column represents one of the top 5 highest-Bayes-factor comparisons (exact value at bottom; $> 100$ indicates strong evidence for TSP). The 4 colored blocks in each column include two red blocks (human sequences) that were tested against two sequences from other species (see Methods). Grayed-out rows correspond to sequences that were not considered for Bayes factors, either because they belong to a non-orthologous gene or backbone sequence (and thus not relevant to compare with the human gene in question), or because they may have been involved in a gene conversion event in this exon (according to *GENECONV* or from the literature).

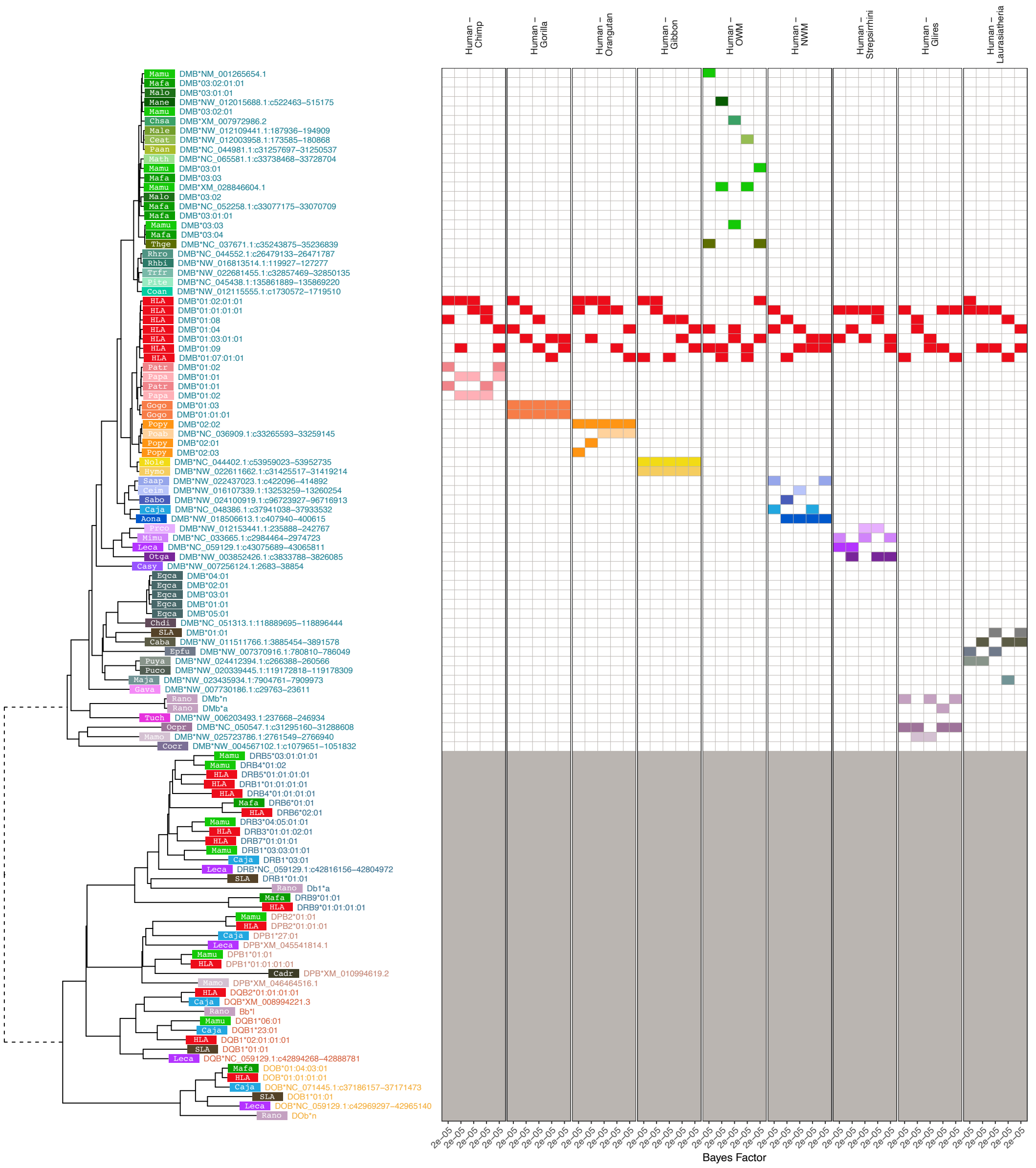


**Figure 2—figure supplement 15. MHC-DMB-Related Group *BEAST2* tree for exon 2 (PBR-encoding).** In the tree, each tip represents a sequence (see ***Appendix 1*** for more details on nomenclature), with the colored rectangle and four-letter abbreviation indicating the species (see ***Figure 1—figure Supplement 2*** for full species key). Following the rectangle, tips are labeled with the sequence name; sequences which have been assigned to loci are colored according to the gene group, while unassigned sequences are written in gray. Dashed branches are shortened to 10% of their length to expand detail in the rest of the tree. The left-hand side shows example Bayes factors we calculated from the set of posterior trees; the tree tips correspond to the rows of the grid. Each panel represents a type of comparison (labeled at top) and each column represents one of the top 5 highest-Bayes-factor comparisons (exact value at bottom; $> 100$ indicates strong evidence for TSP). The 4 colored blocks in each column include two red blocks (human sequences) that were tested against two sequences from other species (see Methods). Grayed-out rows correspond to sequences that were not considered for Bayes factors, either because they belong to a non-orthologous gene or backbone sequence (and thus not relevant to compare with the human gene in question), or because they may have been involved in a gene conversion event in this exon (according to *GENECONV* or from the literature).

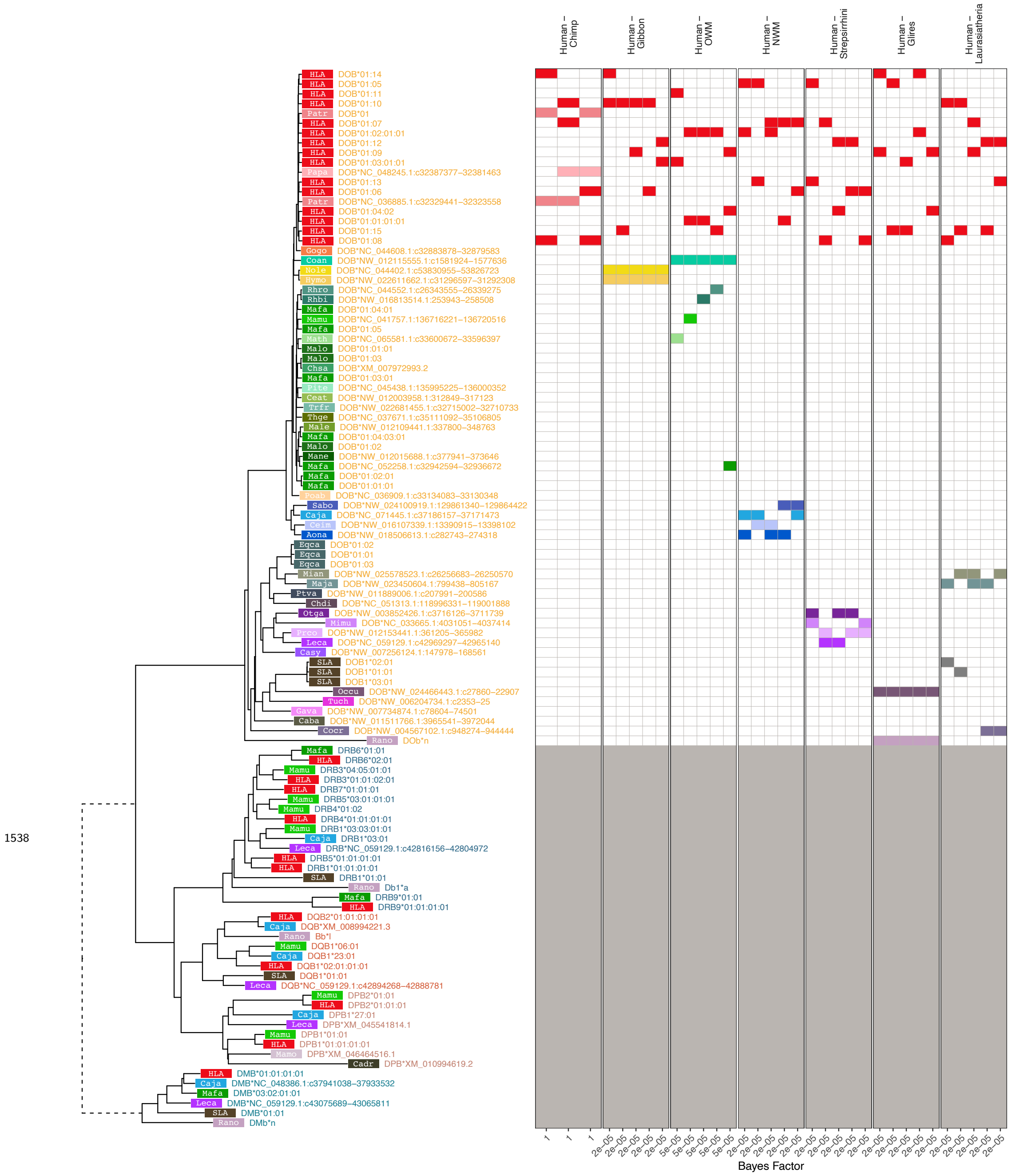


**Figure 2—figure supplement 16. MHC-DOB-Related Group *BEAST2* tree for exon 2 (PBR-encoding).** In the tree, each tip represents a sequence (see ***Appendix 1*** for more details on nomenclature), with the colored rectangle and four-letter abbreviation indicating the species (see ***Figure 1—figure Supplement 2*** for full species key). Following the rectangle, tips are labeled with the sequence name; sequences which have been assigned to loci are colored according to the gene group, while unassigned sequences are written in gray. Dashed branches are shortened to 10% of their length to expand detail in the rest of the tree. The left-hand side shows example Bayes factors we calculated from the set of posterior trees; the tree tips correspond to the rows of the grid. Each panel represents a type of comparison (labeled at top) and each column represents one of the top 5 highest-Bayes-factor comparisons (exact value at bottom; $> 100$ indicates strong evidence for TSP). The 4 colored blocks in each column include two red blocks (human sequences) that were tested against two sequences from other species (see Methods). Grayed-out rows correspond to sequences that were not considered for Bayes factors, either because they belong to a non-orthologous gene or backbone sequence (and thus not relevant to compare with the human gene in question), or because they may have been involved in a gene conversion event in this exon (according to *GENECONV* or from the literature).

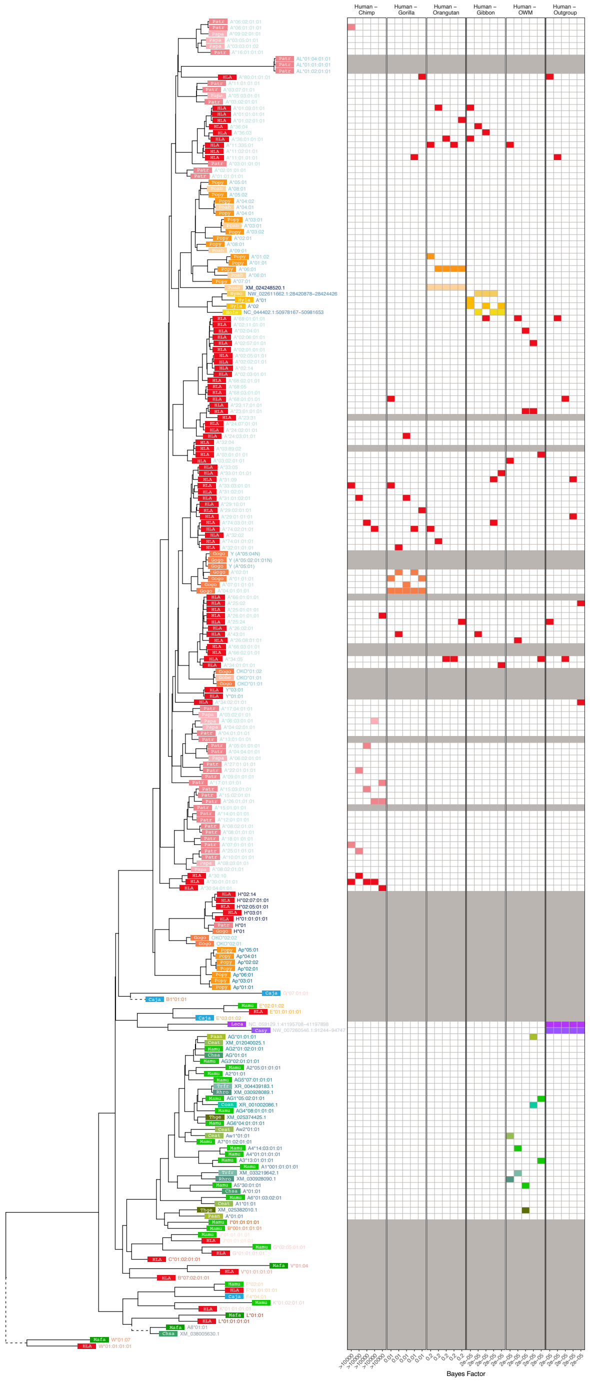


**Figure 3—figure supplement 1. MHC-A-Related Group *BEAST2* tree for exon 3 (PBR-encoding).** In the tree, each tip represents a sequence (see ***Appendix 1*** for more details on nomenclature), with the colored rectangle and four-letter abbreviation indicating the species (see ***Figure 1—figure Supplement 2*** for full species key). Following the rectangle, tips are labeled with the sequence name; sequences which have been assigned to loci are colored according to the gene group, while unassigned sequences are written in gray. Dashed branches are shortened to 10% of their length to expand detail in the rest of the tree. The left-hand side shows example Bayes factors we calculated from the set of posterior trees; the tree tips correspond to the rows of the grid. Each panel represents a type of comparison (labeled at top) and each column represents one of the top 5 highest-Bayes-factor comparisons (exact value at bottom; $> 100$ indicates strong evidence for TSP). The 4 colored blocks in each column include two red blocks (human sequences) that were tested against two sequences from other species (see Methods). Grayed-out rows correspond to sequences that were not considered for Bayes factors, either because they belong to a non-orthologous gene or backbone sequence (and thus not relevant to compare with the human gene in question), or because they may have been involved in a gene conversion event in this exon (according to *GENECONV* or from the literature).

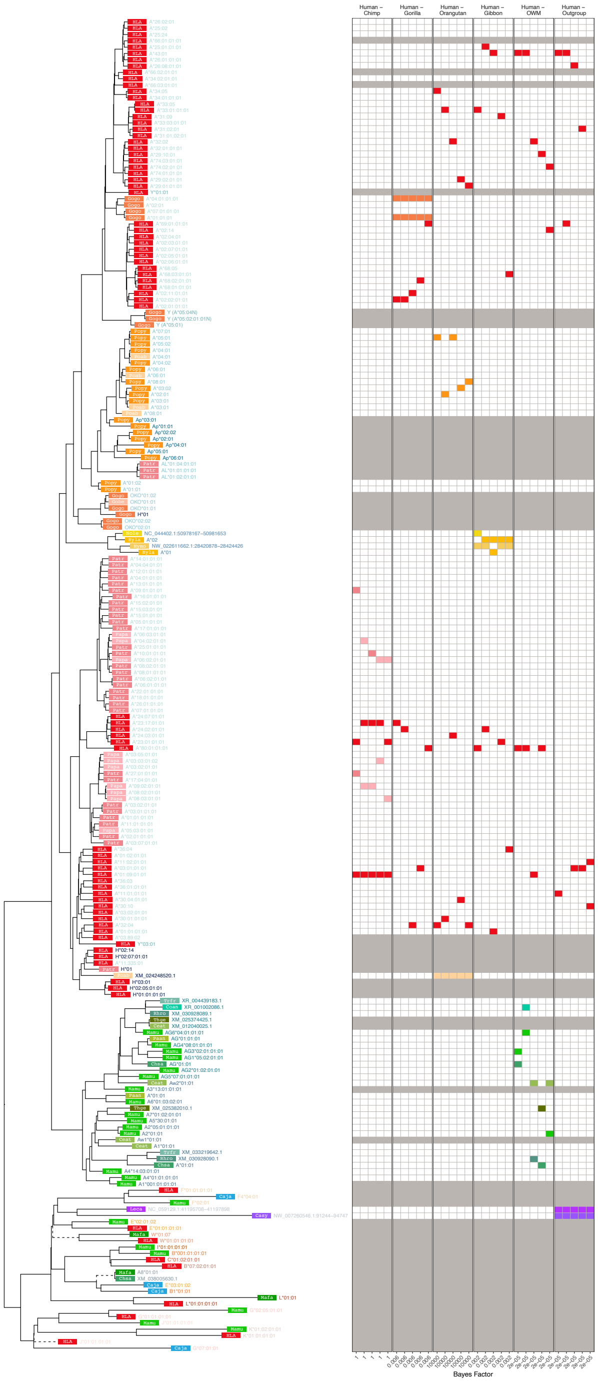


**Figure 3—figure supplement 2. MHC-A-Related Group *BEAST2* tree for exon 4 (PBR-encoding).** In the tree, each tip represents a sequence (see ***Appendix 1*** for more details on nomenclature), with the colored rectangle and four-letter abbreviation indicating the species (see ***Figure 1—figure Supplement 2*** for full species key). Following the rectangle, tips are labeled with the sequence name; sequences which have been assigned to loci are colored according to the gene group, while unassigned sequences are written in gray. Dashed branches are shortened to 10% of their length to expand detail in the rest of the tree. The left-hand side shows example Bayes factors we calculated from the set of posterior trees; the tree tips correspond to the rows of the grid. Each panel represents a type of comparison (labeled at top) and each column represents one of the top 5 highest-Bayes-factor comparisons (exact value at bottom; $> 100$ indicates strong evidence for TSP). The 4 colored blocks in each column include two red blocks (human sequences) that were tested against two sequences from other species (see Methods). Grayed-out rows correspond to sequences that were not considered for Bayes factors, either because they belong to a non-orthologous gene or backbone sequence (and thus not relevant to compare with the human gene in question), or because they may have been involved in a gene conversion event in this exon (according to *GENECONV* or from the literature).

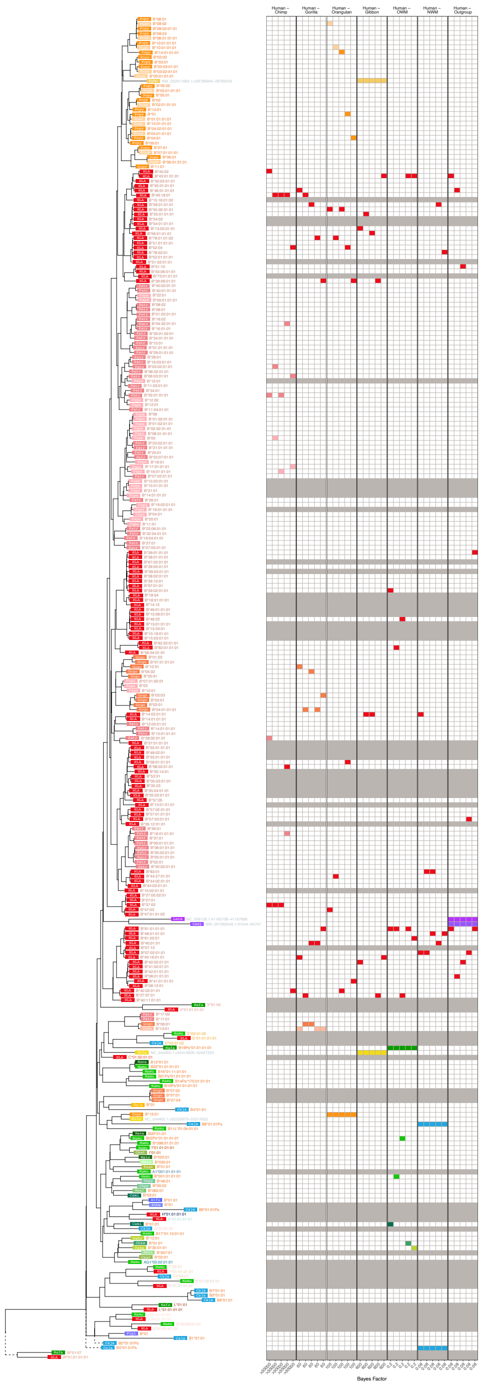


**Figure 3—figure supplement 3. MHC-B-Related Group *BEAST2* tree for exon 3 (PBR-encoding).** In the tree, each tip represents a sequence (see ***Appendix 1*** for more details on nomenclature), with the colored rectangle and four-letter abbreviation indicating the species (see ***Figure 1—figure Supplement 2*** for full species key). Following the rectangle, tips are labeled with the sequence name; sequences which have been assigned to loci are colored according to the gene group, while unassigned sequences are written in gray. Dashed branches are shortened to 10% of their length to expand detail in the rest of the tree. The left-hand side shows example Bayes factors we calculated from the set of posterior trees; the tree tips correspond to the rows of the grid. Each panel represents a type of comparison (labeled at top) and each column represents one of the top 5 highest-Bayes-factor comparisons (exact value at bottom; $> 100$ indicates strong evidence for TSP). The 4 colored blocks in each column include two red blocks (human sequences) that were tested against two sequences from other species (see Methods). Grayed-out rows correspond to sequences that were not considered for Bayes factors, either because they belong to a non-orthologous gene or backbone sequence (and thus not relevant to compare with the human gene in question), or because they may have been involved in a gene conversion event in this exon (according to *GENECONV* or from the literature).

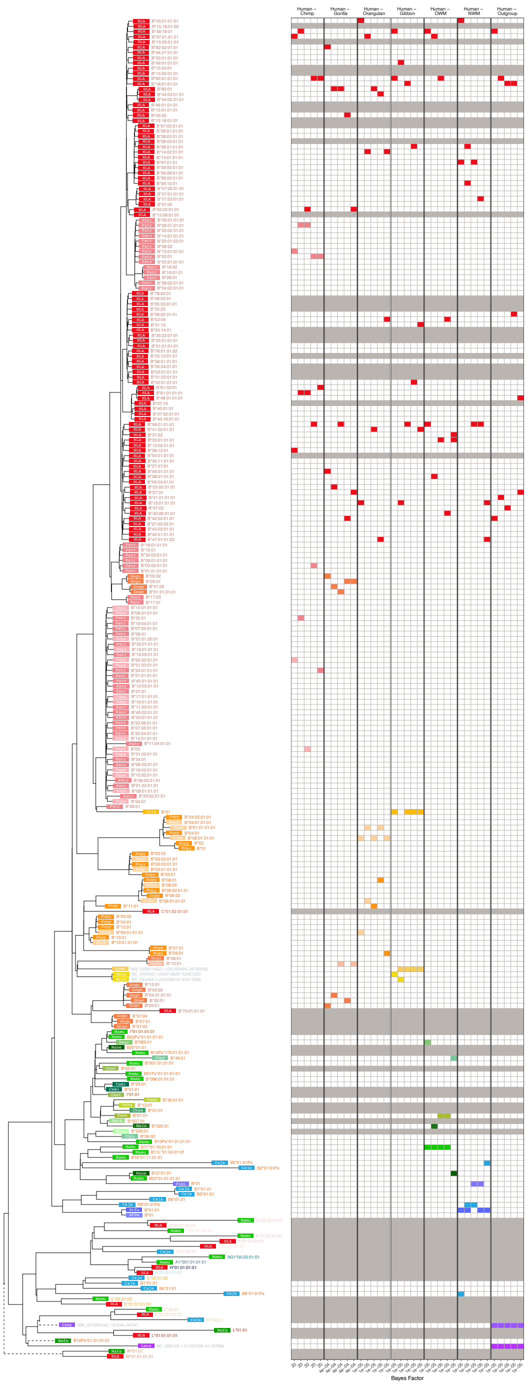


**Figure 3—figure supplement 4. MHC-B-Related Group *BEAST2* tree for exon 4 (PBR-encoding).** In the tree, each tip represents a sequence (see ***Appendix 1*** for more details on nomenclature), with the colored rectangle and four-letter abbreviation indicating the species (see ***Figure 1—figure Supplement 2*** for full species key). Following the rectangle, tips are labeled with the sequence name; sequences which have been assigned to loci are colored according to the gene group, while unassigned sequences are written in gray. Dashed branches are shortened to 10% of their length to expand detail in the rest of the tree. The left-hand side shows example Bayes factors we calculated from the set of posterior trees; the tree tips correspond to the rows of the grid. Each panel represents a type of comparison (labeled at top) and each column represents one of the top 5 highest-Bayes-factor comparisons (exact value at bottom; $> 100$ indicates strong evidence for TSP). The 4 colored blocks in each column include two red blocks (human sequences) that were tested against two sequences from other species (see Methods). Grayed-out rows correspond to sequences that were not considered for Bayes factors, either because they belong to a non-orthologous gene or backbone sequence (and thus not relevant to compare with the human gene in question), or because they may have been involved in a gene conversion event in this exon (according to *GENECONV* or from the literature).

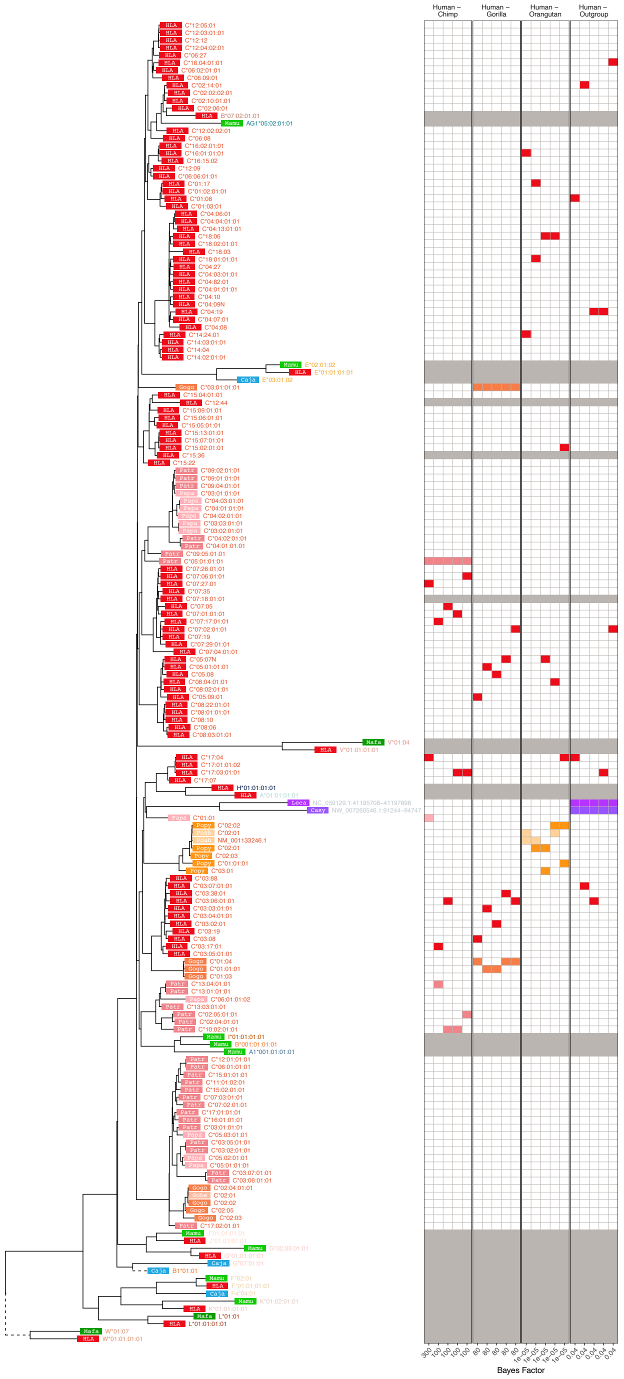


**Figure 3—figure supplement 5. MHC-C-Related Group *BEAST2* tree for exon 3 (PBR-encoding).** In the tree, each tip represents a sequence (see ***Appendix 1*** for more details on nomenclature), with the colored rectangle and four-letter abbreviation indicating the species (see ***Figure 1—figure Supplement 2*** for full species key). Following the rectangle, tips are labeled with the sequence name; sequences which have been assigned to loci are colored according to the gene group, while unassigned sequences are written in gray. Dashed branches are shortened to 10% of their length to expand detail in the rest of the tree. The left-hand side shows example Bayes factors we calculated from the set of posterior trees; the tree tips correspond to the rows of the grid. Each panel represents a type of comparison (labeled at top) and each column represents one of the top 5 highest-Bayes-factor comparisons (exact value at bottom; $> 100$ indicates strong evidence for TSP). The 4 colored blocks in each column include two red blocks (human sequences) that were tested against two sequences from other species (see Methods). Grayed-out rows correspond to sequences that were not considered for Bayes factors, either because they belong to a non-orthologous gene or backbone sequence (and thus not relevant to compare with the human gene in question), or because they may have been involved in a gene conversion event in this exon (according to *GENECONV* or from the literature).

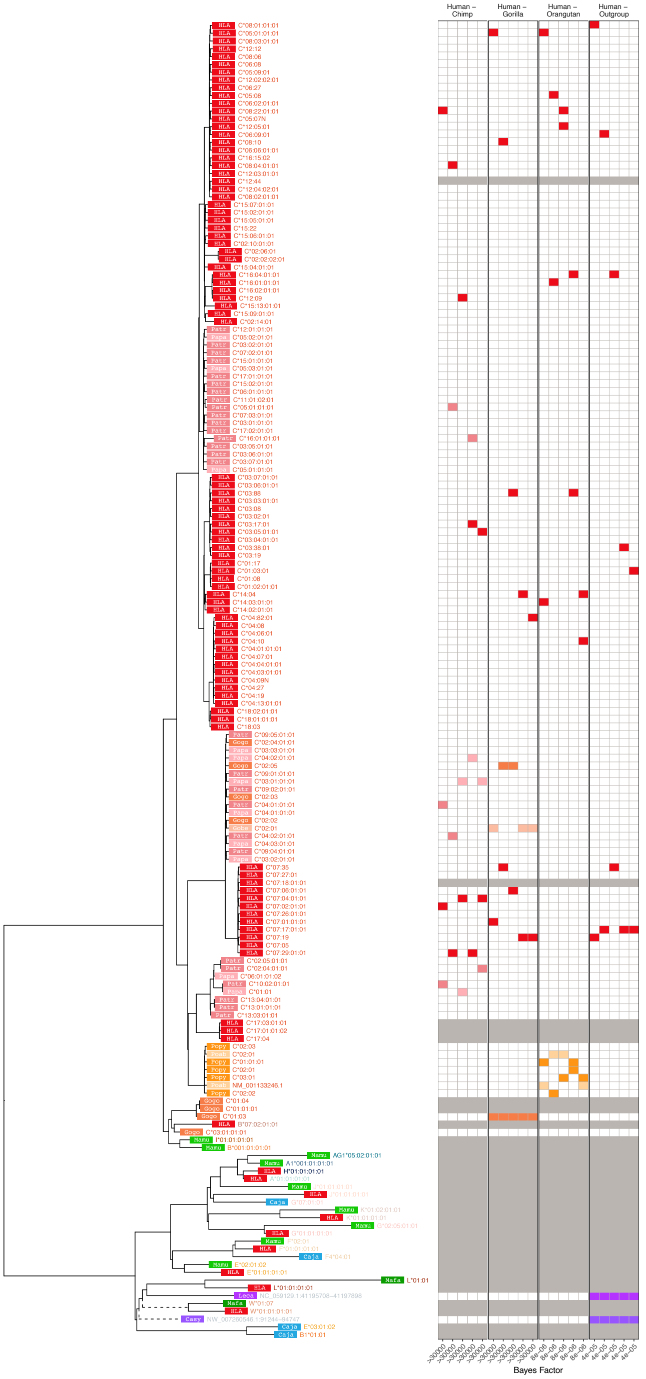


**Figure 3—figure supplement 6. MHC-C-Related Group *BEAST2* tree for exon 4 (PBR-encoding).** In the tree, each tip represents a sequence (see ***Appendix 1*** for more details on nomenclature), with the colored rectangle and four-letter abbreviation indicating the species (see ***Figure 1—figure Supplement 2*** for full species key). Following the rectangle, tips are labeled with the sequence name; sequences which have been assigned to loci are colored according to the gene group, while unassigned sequences are written in gray. Dashed branches are shortened to 10% of their length to expand detail in the rest of the tree. The left-hand side shows example Bayes factors we calculated from the set of posterior trees; the tree tips correspond to the rows of the grid. Each panel represents a type of comparison (labeled at top) and each column represents one of the top 5 highest-Bayes-factor comparisons (exact value at bottom; $> 100$ indicates strong evidence for TSP). The 4 colored blocks in each column include two red blocks (human sequences) that were tested against two sequences from other species (see Methods). Grayed-out rows correspond to sequences that were not considered for Bayes factors, either because they belong to a non-orthologous gene or backbone sequence (and thus not relevant to compare with the human gene in question), or because they may have been involved in a gene conversion event in this exon (according to *GENECONV* or from the literature).

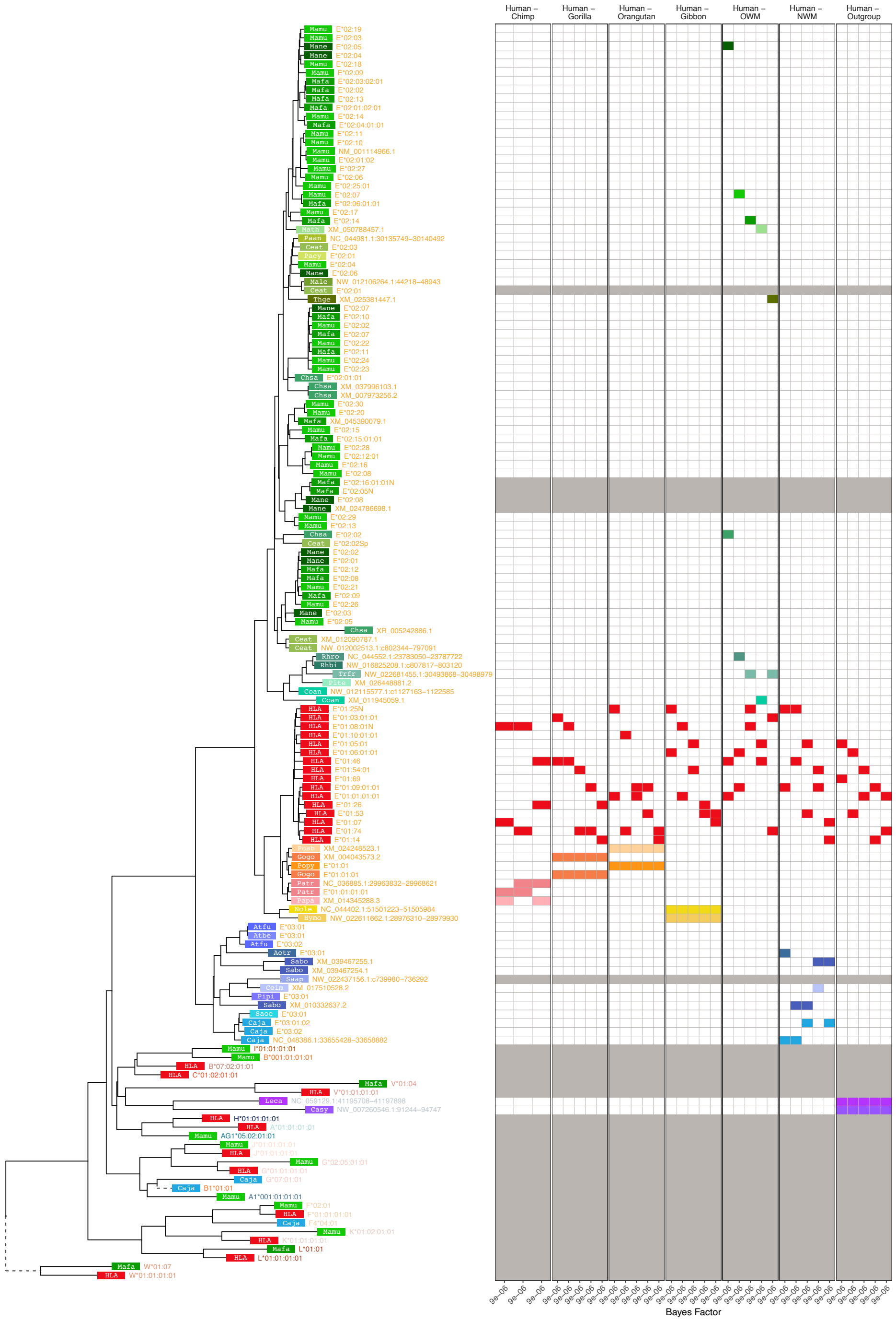


**Figure 3—figure supplement 7. MHC-E-Related Group *BEAST2* tree for exon 3 (PBR-encoding).** In the tree, each tip represents a sequence (see ***Appendix 1*** for more details on nomenclature), with the colored rectangle and four-letter abbreviation indicating the species (see ***Figure 1—figure Supplement 2*** for full species key). Following the rectangle, tips are labeled with the sequence name; sequences which have been assigned to loci are colored according to the gene group, while unassigned sequences are written in gray. Dashed branches are shortened to 10% of their length to expand detail in the rest of the tree. The left-hand side shows example Bayes factors we calculated from the set of posterior trees; the tree tips correspond to the rows of the grid. Each panel represents a type of comparison (labeled at top) and each column represents one of the top 5 highest-Bayes-factor comparisons (exact value at bottom; $> 100$ indicates strong evidence for TSP). The 4 colored blocks in each column include two red blocks (human sequences) that were tested against two sequences from other species (see Methods). Grayed-out rows correspond to sequences that were not considered for Bayes factors, either because they belong to a non-orthologous gene or backbone sequence (and thus not relevant to compare with the human gene in question), or because they may have been involved in a gene conversion event in this exon (according to *GENECONV* or from the literature).

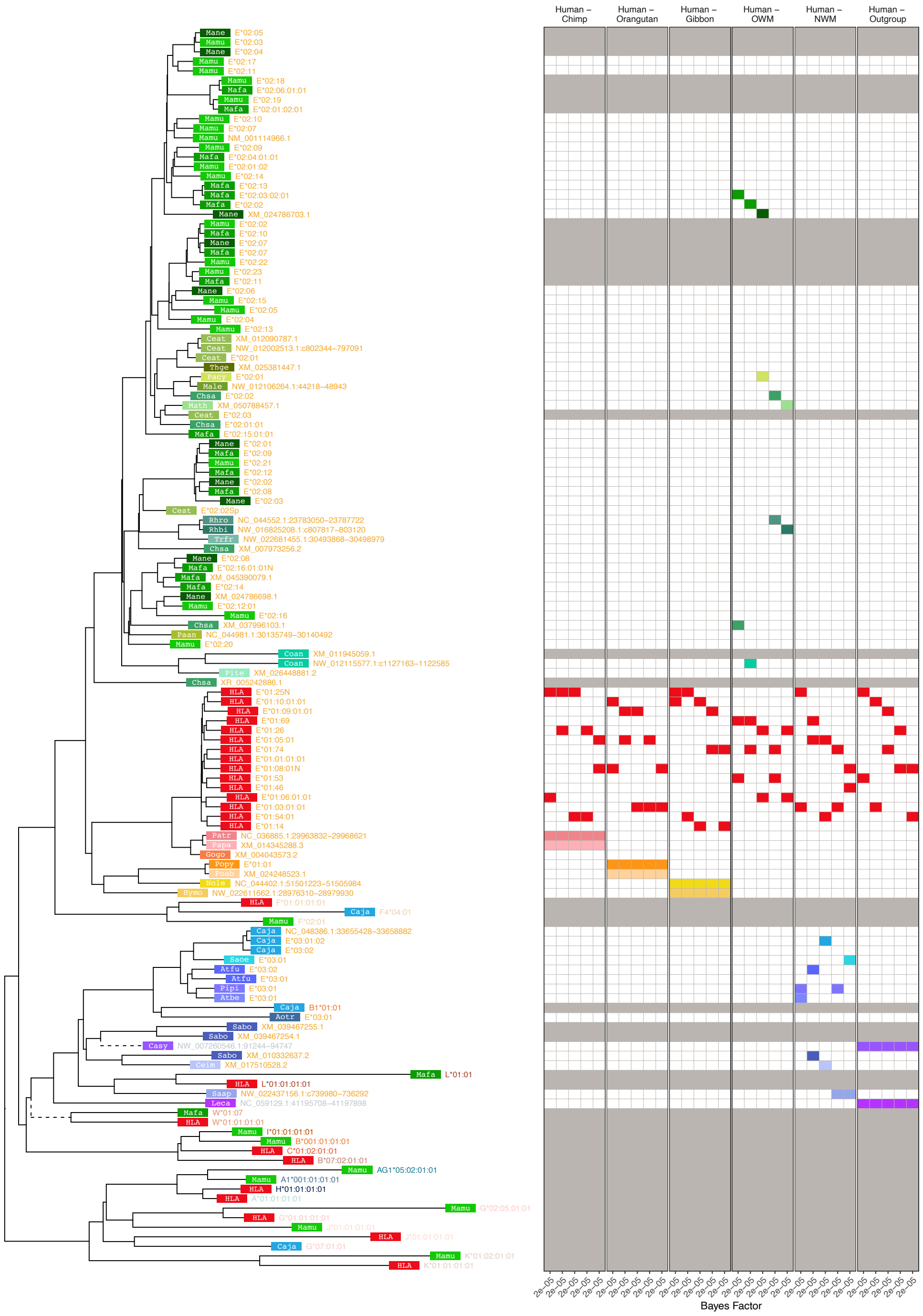


**Figure 3—figure supplement 8. MHC-E-Related Group *BEAST2* tree for exon 4 (PBR-encoding).** In the tree, each tip represents a sequence (see ***Appendix 1*** for more details on nomenclature), with the colored rectangle and four-letter abbreviation indicating the species (see ***Figure 1—figure Supplement 2*** for full species key). Following the rectangle, tips are labeled with the sequence name; sequences which have been assigned to loci are colored according to the gene group, while unassigned sequences are written in gray. Dashed branches are shortened to 10% of their length to expand detail in the rest of the tree. The left-hand side shows example Bayes factors we calculated from the set of posterior trees; the tree tips correspond to the rows of the grid. Each panel represents a type of comparison (labeled at top) and each column represents one of the top 5 highest-Bayes-factor comparisons (exact value at bottom; $> 100$ indicates strong evidence for TSP). The 4 colored blocks in each column include two red blocks (human sequences) that were tested against two sequences from other species (see Methods). Grayed-out rows correspond to sequences that were not considered for Bayes factors, either because they belong to a non-orthologous gene or backbone sequence (and thus not relevant to compare with the human gene in question), or because they may have been involved in a gene conversion event in this exon (according to *GENECONV* or from the literature).

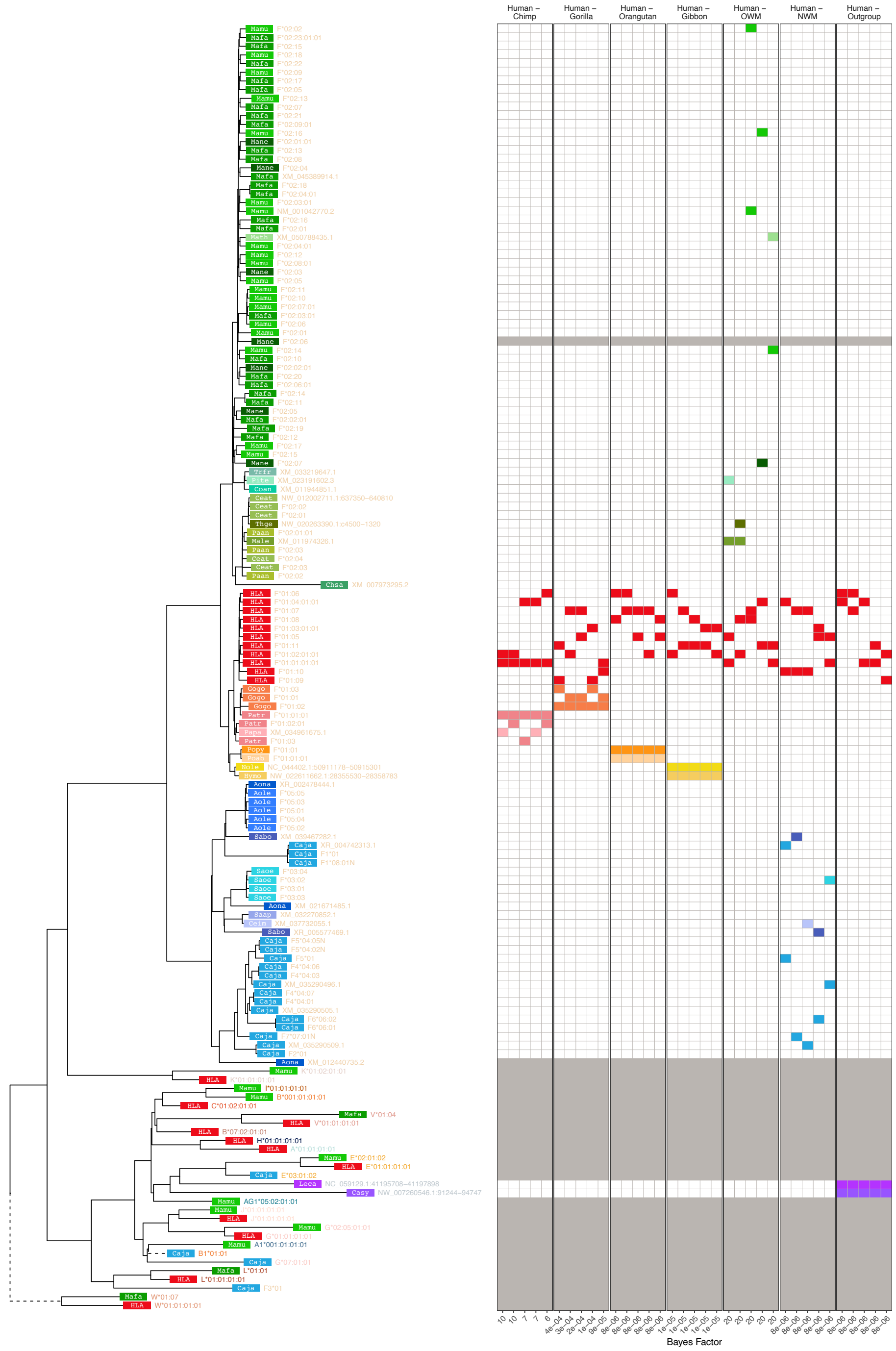


**Figure 3—figure supplement 9. MHC-F-Related Group *BEAST2* tree for exon 3 (PBR-encoding).** In the tree, each tip represents a sequence (see ***Appendix 1*** for more details on nomenclature), with the colored rectangle and four-letter abbreviation indicating the species (see ***Figure 1—figure Supplement 2*** for full species key). Following the rectangle, tips are labeled with the sequence name; sequences which have been assigned to loci are colored according to the gene group, while unassigned sequences are written in gray. Dashed branches are shortened to 10% of their length to expand detail in the rest of the tree. The left-hand side shows example Bayes factors we calculated from the set of posterior trees; the tree tips correspond to the rows of the grid. Each panel represents a type of comparison (labeled at top) and each column represents one of the top 5 highest-Bayes-factor comparisons (exact value at bottom; $> 100$ indicates strong evidence for TSP). The 4 colored blocks in each column include two red blocks (human sequences) that were tested against two sequences from other species (see Methods). Grayed-out rows correspond to sequences that were not considered for Bayes factors, either because they belong to a non-orthologous gene or backbone sequence (and thus not relevant to compare with the human gene in question), or because they may have been involved in a gene conversion event in this exon (according to *GENECONV* or from the literature).

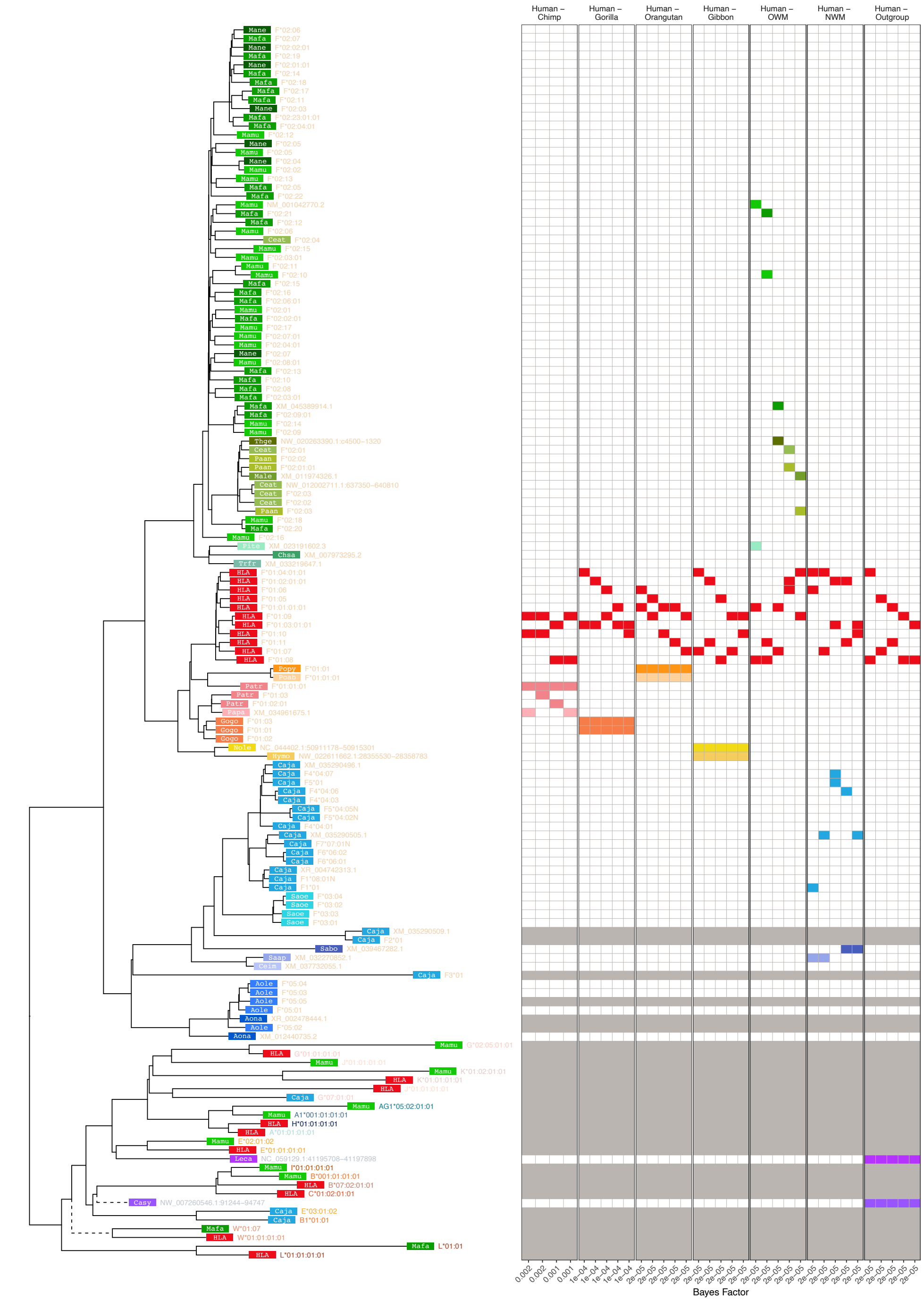


**Figure 3—figure supplement 10. MHC-F-Related Group *BEAST2* tree for exon 4 (PBR-encoding).** In the tree, each tip represents a sequence (see ***Appendix 1*** for more details on nomenclature), with the colored rectangle and four-letter abbreviation indicating the species (see ***Figure 1—figure Supplement 2*** for full species key). Following the rectangle, tips are labeled with the sequence name; sequences which have been assigned to loci are colored according to the gene group, while unassigned sequences are written in gray. Dashed branches are shortened to 10% of their length to expand detail in the rest of the tree. The left-hand side shows example Bayes factors we calculated from the set of posterior trees; the tree tips correspond to the rows of the grid. Each panel represents a type of comparison (labeled at top) and each column represents one of the top 5 highest-Bayes-factor comparisons (exact value at bottom; $> 100$ indicates strong evidence for TSP). The 4 colored blocks in each column include two red blocks (human sequences) that were tested against two sequences from other species (see Methods). Grayed-out rows correspond to sequences that were not considered for Bayes factors, either because they belong to a non-orthologous gene or backbone sequence (and thus not relevant to compare with the human gene in question), or because they may have been involved in a gene conversion event in this exon (according to *GENECONV* or from the literature).

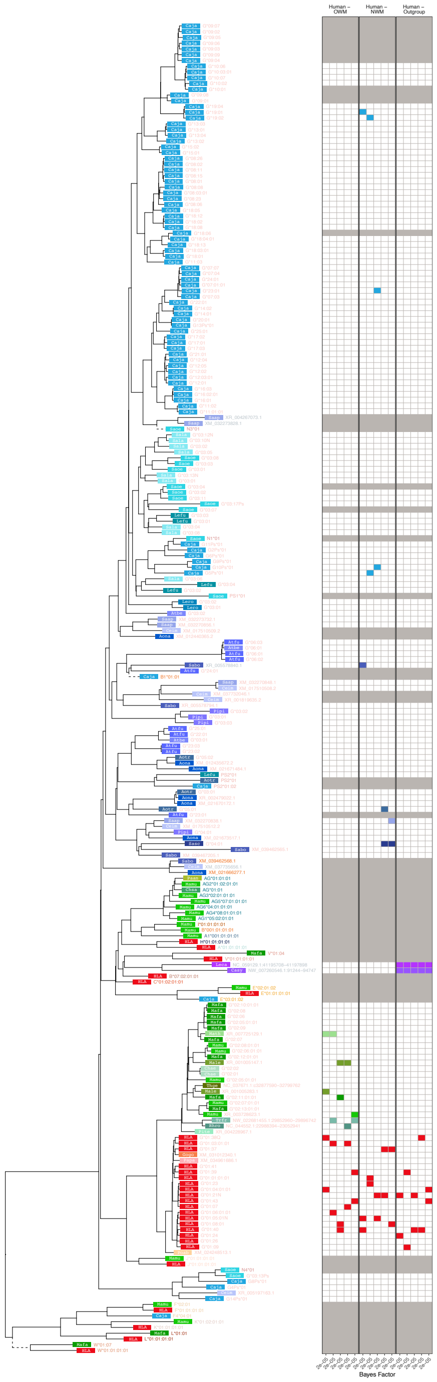


**Figure 3—figure supplement 11. MHC-G-Related Group *BEAST2* tree for exon 3 (PBR-encoding).** In the tree, each tip represents a sequence (see ***Appendix 1*** for more details on nomenclature), with the colored rectangle and four-letter abbreviation indicating the species (see ***Figure 1—figure Supplement 2*** for full species key). Following the rectangle, tips are labeled with the sequence name; sequences which have been assigned to loci are colored according to the gene group, while unassigned sequences are written in gray. Dashed branches are shortened to 10% of their length to expand detail in the rest of the tree. The left-hand side shows example Bayes factors we calculated from the set of posterior trees; the tree tips correspond to the rows of the grid. Each panel represents a type of comparison (labeled at top) and each column represents one of the top 5 highest-Bayes-factor comparisons (exact value at bottom; $> 100$ indicates strong evidence for TSP). The 4 colored blocks in each column include two red blocks (human sequences) that were tested against two sequences from other species (see Methods). Grayed-out rows correspond to sequences that were not considered for Bayes factors, either because they belong to a non-orthologous gene or backbone sequence (and thus not relevant to compare with the human gene in question), or because they may have been involved in a gene conversion event in this exon (according to *GENECONV* or from the literature).

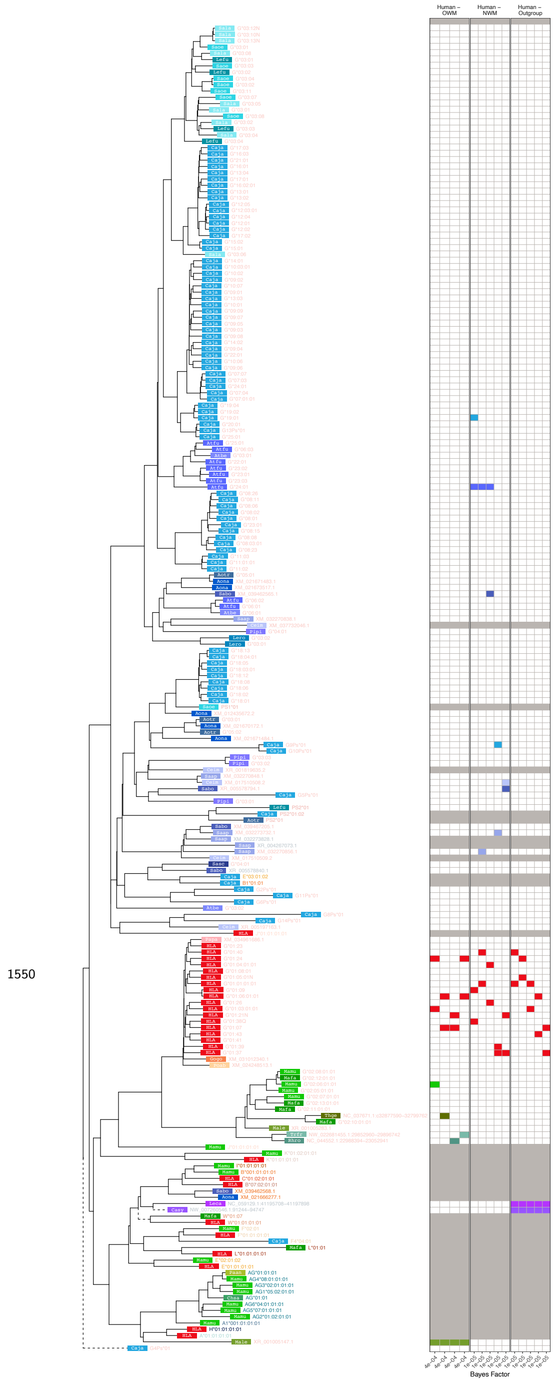


**Figure 3—figure supplement 12. MHC-G-Related Group *BEAST2* tree for exon 4 (PBR-encoding).** In the tree, each tip represents a sequence (see ***Appendix 1*** for more details on nomenclature), with the colored rectangle and four-letter abbreviation indicating the species (see ***Figure 1—figure Supplement 2*** for full species key). Following the rectangle, tips are labeled with the sequence name; sequences which have been assigned to loci are colored according to the gene group, while unassigned sequences are written in gray. Dashed branches are shortened to 10% of their length to expand detail in the rest of the tree. The left-hand side shows example Bayes factors we calculated from the set of posterior trees; the tree tips correspond to the rows of the grid. Each panel represents a type of comparison (labeled at top) and each column represents one of the top 5 highest-Bayes-factor comparisons (exact value at bottom; $> 100$ indicates strong evidence for TSP). The 4 colored blocks in each column include two red blocks (human sequences) that were tested against two sequences from other species (see Methods). Grayed-out rows correspond to sequences that were not considered for Bayes factors, either because they belong to a non-orthologous gene or backbone sequence (and thus not relevant to compare with the human gene in question), or because they may have been involved in a gene conversion event in this exon (according to *GENECONV* or from the literature).

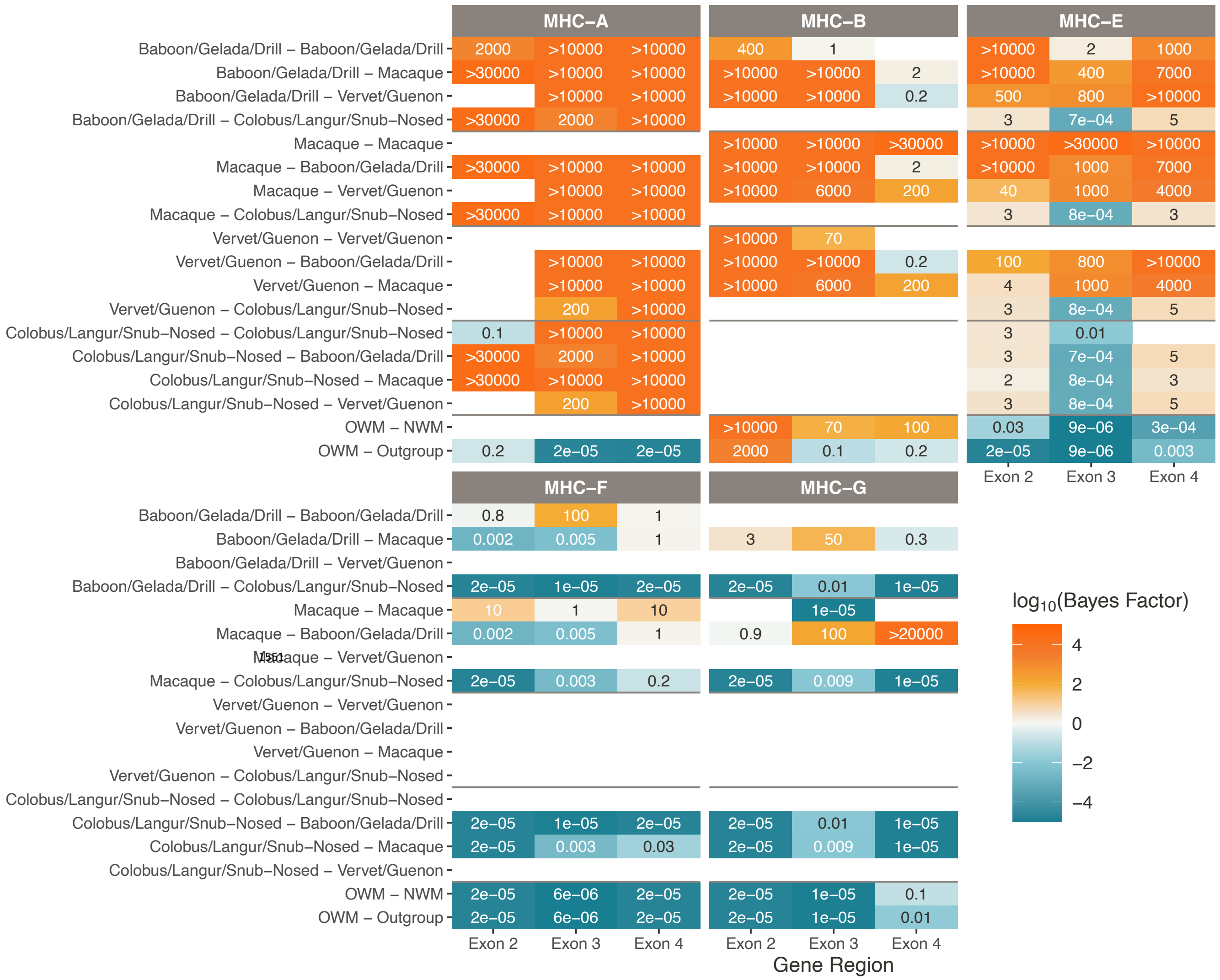


**Figure 3—figure supplement 13. TSP among old-world monkey groups for the Class I genes.** Different species comparisons are listed on the y-axis, and different gene regions are listed on the x-axis. Each table entry is colored and labeled with the maximum Bayes factor among all tested quartets of alleles belonging to that category. High Bayes factors (orange) indicate support for TSP among the given species for that gene region, while low Bayes factors (teal) indicate that alleles assort according to the species tree, as expected. Bayes factors above 100 are considered decisive. Tan values show poor support for either hypothesis, while white boxes indicate that there are not enough alleles in that category with which to calculate Bayes factors. MHC-C was not present before the human-orangutan ancestor, so it is not possible to calculate Bayes factors for MHC-C for these species comparisons.

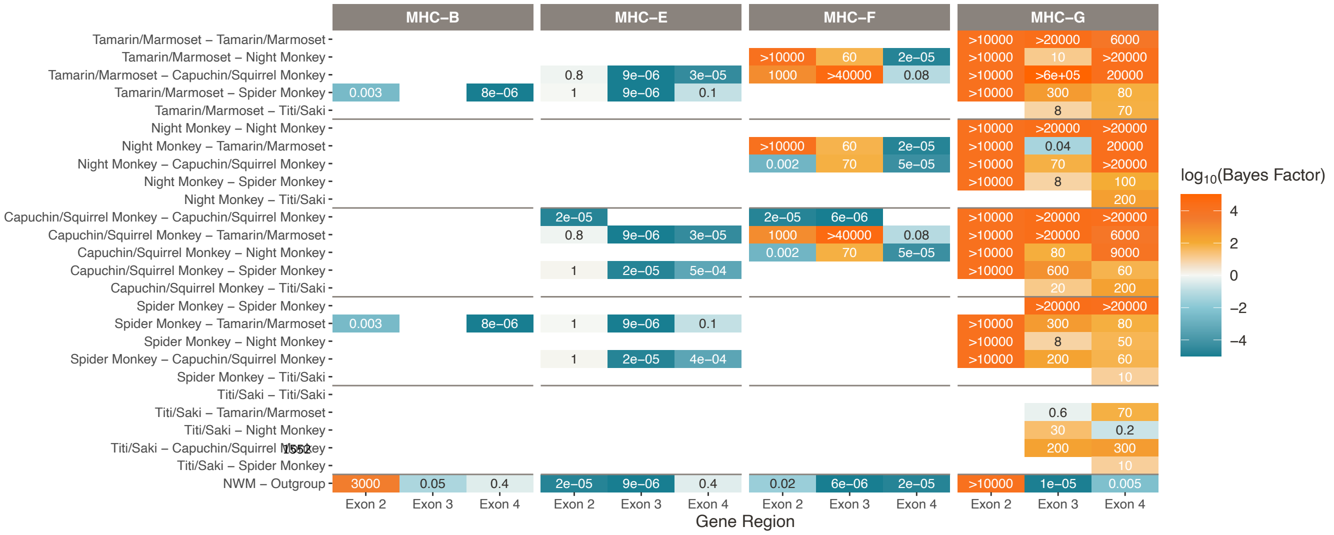


**Figure 3—figure supplement 14. TSP among new-world monkey groups for the Class I genes.** Different species comparisons are listed on the y-axis, and different gene regions are listed on the x-axis. Each table entry is colored and labeled with the maximum Bayes factor among all tested quartets of alleles belonging to that category. High Bayes factors (orange) indicate support for TSP among the given species for that gene region, while low Bayes factors (teal) indicate that alleles assort according to the species tree, as expected. Bayes factors above 100 are considered decisive. Tan values show poor support for either hypothesis, while white boxes indicate that there are not enough alleles in that category with which to calculate Bayes factors. MHC-C was not present before the human-orangutan ancestor, and MHC-A is not present in the NWM, so it is not possible to calculate Bayes factors for these genes.

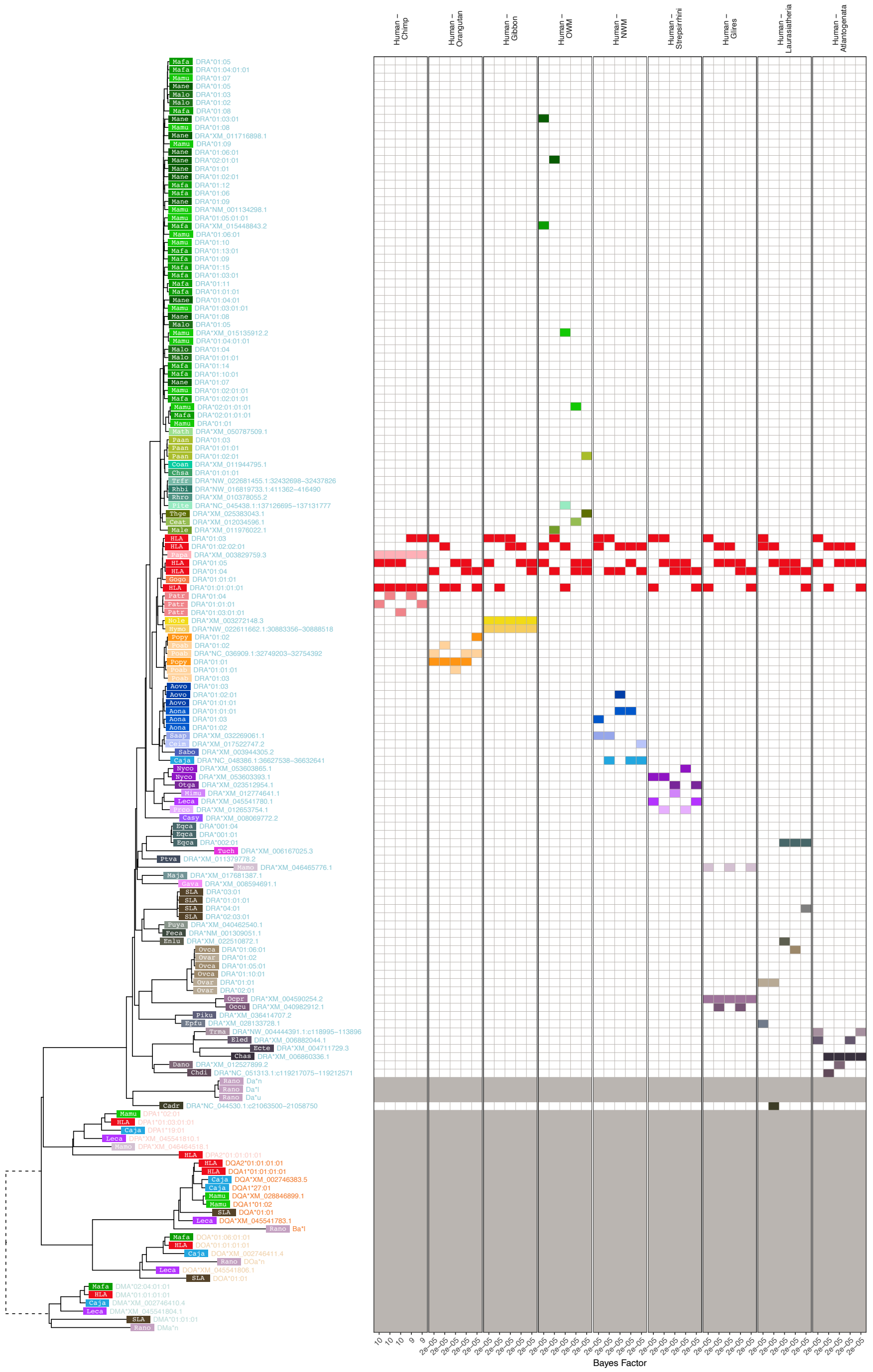


**Figure 4—figure supplement 1. MHC-DRA-Related Group *BEAST2* tree for exon 3 (PBR-encoding).** In the tree, each tip represents a sequence (see ***Appendix 1*** for more details on nomenclature), with the colored rectangle and four-letter abbreviation indicating the species (see ***Figure 1—figure Supplement 2*** for full species key). Following the rectangle, tips are labeled with the sequence name; sequences which have been assigned to loci are colored according to the gene group, while unassigned sequences are written in gray. Dashed branches are shortened to 10% of their length to expand detail in the rest of the tree. The left-hand side shows example Bayes factors we calculated from the set of posterior trees; the tree tips correspond to the rows of the grid. Each panel represents a type of comparison (labeled at top) and each column represents one of the top 5 highest-Bayes-factor comparisons (exact value at bottom; $> 100$ indicates strong evidence for TSP). The 4 colored blocks in each column include two red blocks (human sequences) that were tested against two sequences from other species (see Methods). Grayed-out rows correspond to sequences that were not considered for Bayes factors, either because they belong to a non-orthologous gene or backbone sequence (and thus not relevant to compare with the human gene in question), or because they may have been involved in a gene conversion event in this exon (according to *GENECONV* or from the literature).

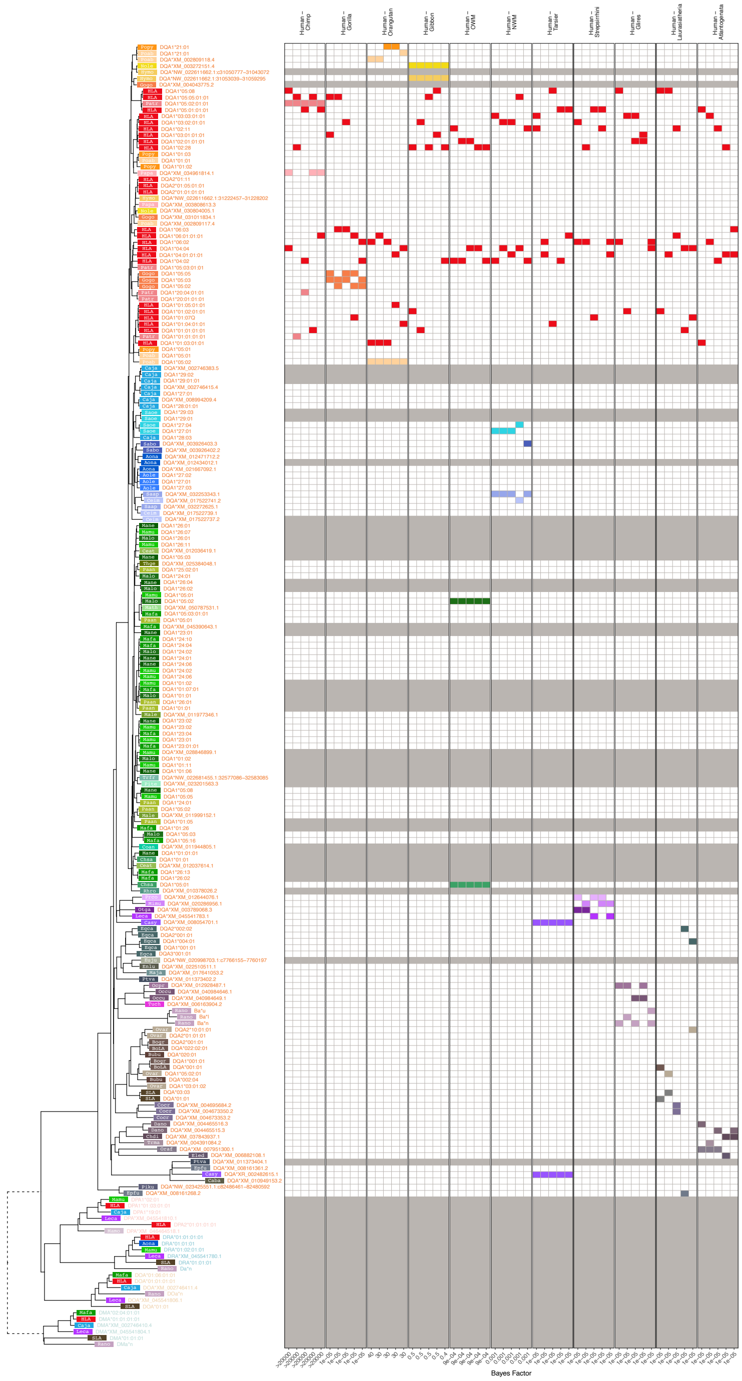


**Figure 4—figure supplement 2. MHC-DQA-Related Group *BEAST2* tree for exon 3 (PBR-encoding).** In the tree, each tip represents a sequence (see ***Appendix 1*** for more details on nomenclature), with the colored rectangle and four-letter abbreviation indicating the species (see ***Figure 1—figure Supplement 2*** for full species key). Following the rectangle, tips are labeled with the sequence name; sequences which have been assigned to loci are colored according to the gene group, while unassigned sequences are written in gray. Dashed branches are shortened to 10% of their length to expand detail in the rest of the tree. The left-hand side shows example Bayes factors we calculated from the set of posterior trees; the tree tips correspond to the rows of the grid. Each panel represents a type of comparison (labeled at top) and each column represents one of the top 5 highest-Bayes-factor comparisons (exact value at bottom; $> 100$ indicates strong evidence for TSP). The 4 colored blocks in each column include two red blocks (human sequences) that were tested against two sequences from other species (see Methods). Grayed-out rows correspond to sequences that were not considered for Bayes factors, either because they belong to a non-orthologous gene or backbone sequence (and thus not relevant to compare with the human gene in question), or because they may have been involved in a gene conversion event in this exon (according to *GENECONV* or from the literature).

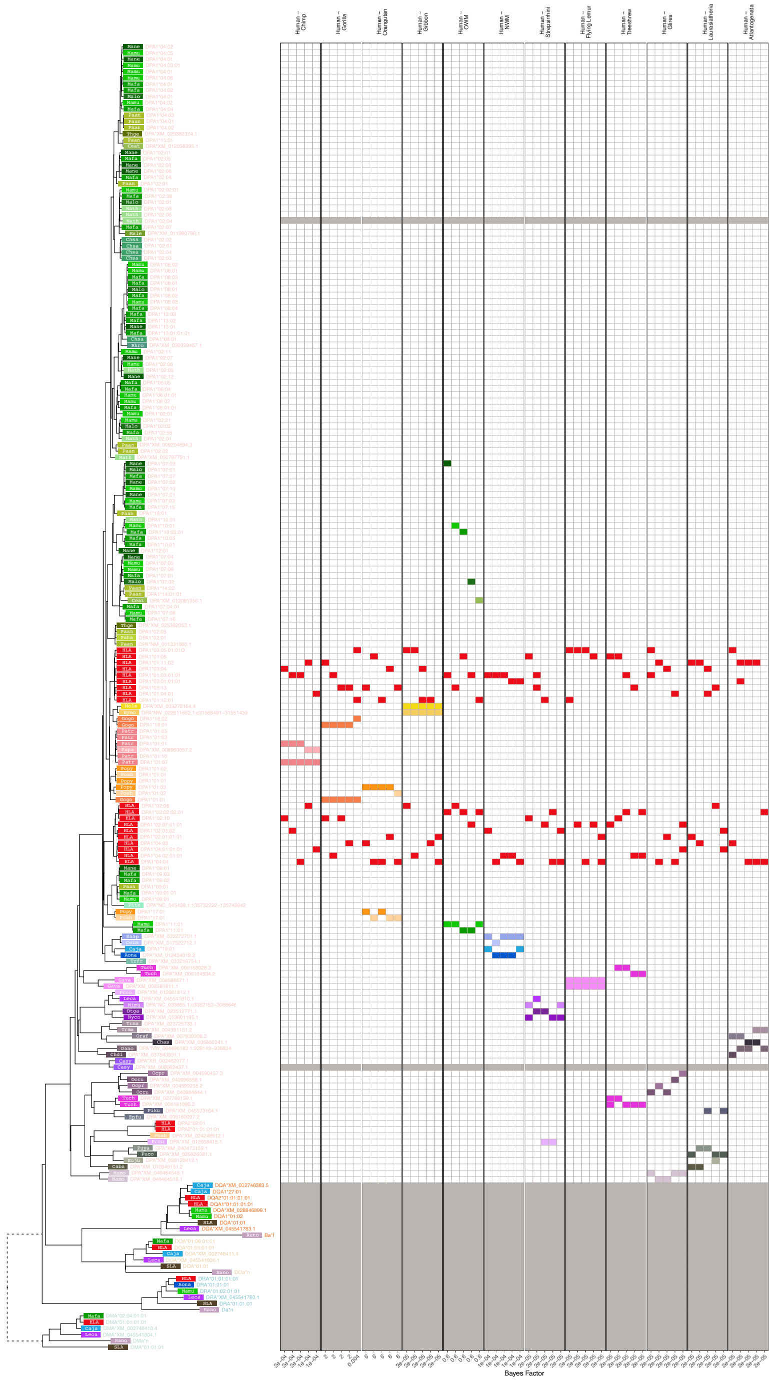


**Figure 4—figure supplement 3. MHC-DPA-Related Group *BEAST2* tree for exon 3 (PBR-encoding).** In the tree, each tip represents a sequence (see ***Appendix 1*** for more details on nomenclature), with the colored rectangle and four-letter abbreviation indicating the species (see ***Figure 1—figure Supplement 2*** for full species key). Following the rectangle, tips are labeled with the sequence name; sequences which have been assigned to loci are colored according to the gene group, while unassigned sequences are written in gray. Dashed branches are shortened to 10% of their length to expand detail in the rest of the tree. The left-hand side shows example Bayes factors we calculated from the set of posterior trees; the tree tips correspond to the rows of the grid. Each panel represents a type of comparison (labeled at top) and each column represents one of the top 5 highest-Bayes-factor comparisons (exact value at bottom; $> 100$ indicates strong evidence for TSP). The 4 colored blocks in each column include two red blocks (human sequences) that were tested against two sequences from other species (see Methods). Grayed-out rows correspond to sequences that were not considered for Bayes factors, either because they belong to a non-orthologous gene or backbone sequence (and thus not relevant to compare with the human gene in question), or because they may have been involved in a gene conversion event in this exon (according to *GENECONV* or from the literature).

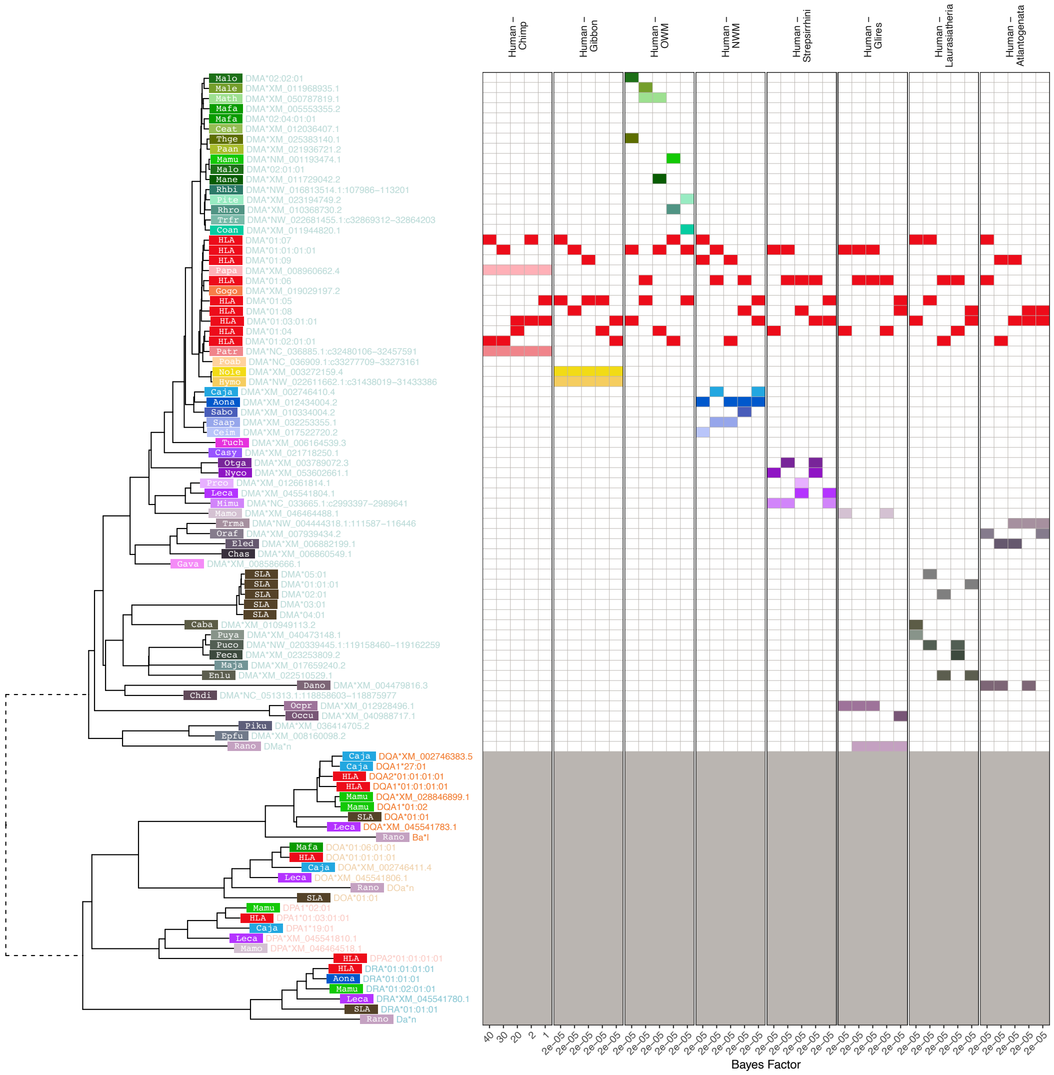


**Figure 4—figure supplement 4. MHC-DMA-Related Group *BEAST2* tree for exon 3 (PBR-encoding).** In the tree, each tip represents a sequence (see ***Appendix 1*** for more details on nomenclature), with the colored rectangle and four-letter abbreviation indicating the species (see ***Figure 1—figure Supplement 2*** for full species key). Following the rectangle, tips are labeled with the sequence name; sequences which have been assigned to loci are colored according to the gene group, while unassigned sequences are written in gray. Dashed branches are shortened to 10% of their length to expand detail in the rest of the tree. The left-hand side shows example Bayes factors we calculated from the set of posterior trees; the tree tips correspond to the rows of the grid. Each panel represents a type of comparison (labeled at top) and each column represents one of the top 5 highest-Bayes-factor comparisons (exact value at bottom; $> 100$ indicates strong evidence for TSP). The 4 colored blocks in each column include two red blocks (human sequences) that were tested against two sequences from other species (see Methods). Grayed-out rows correspond to sequences that were not considered for Bayes factors, either because they belong to a non-orthologous gene or backbone sequence (and thus not relevant to compare with the human gene in question), or because they may have been involved in a gene conversion event in this exon (according to *GENECONV* or from the literature).

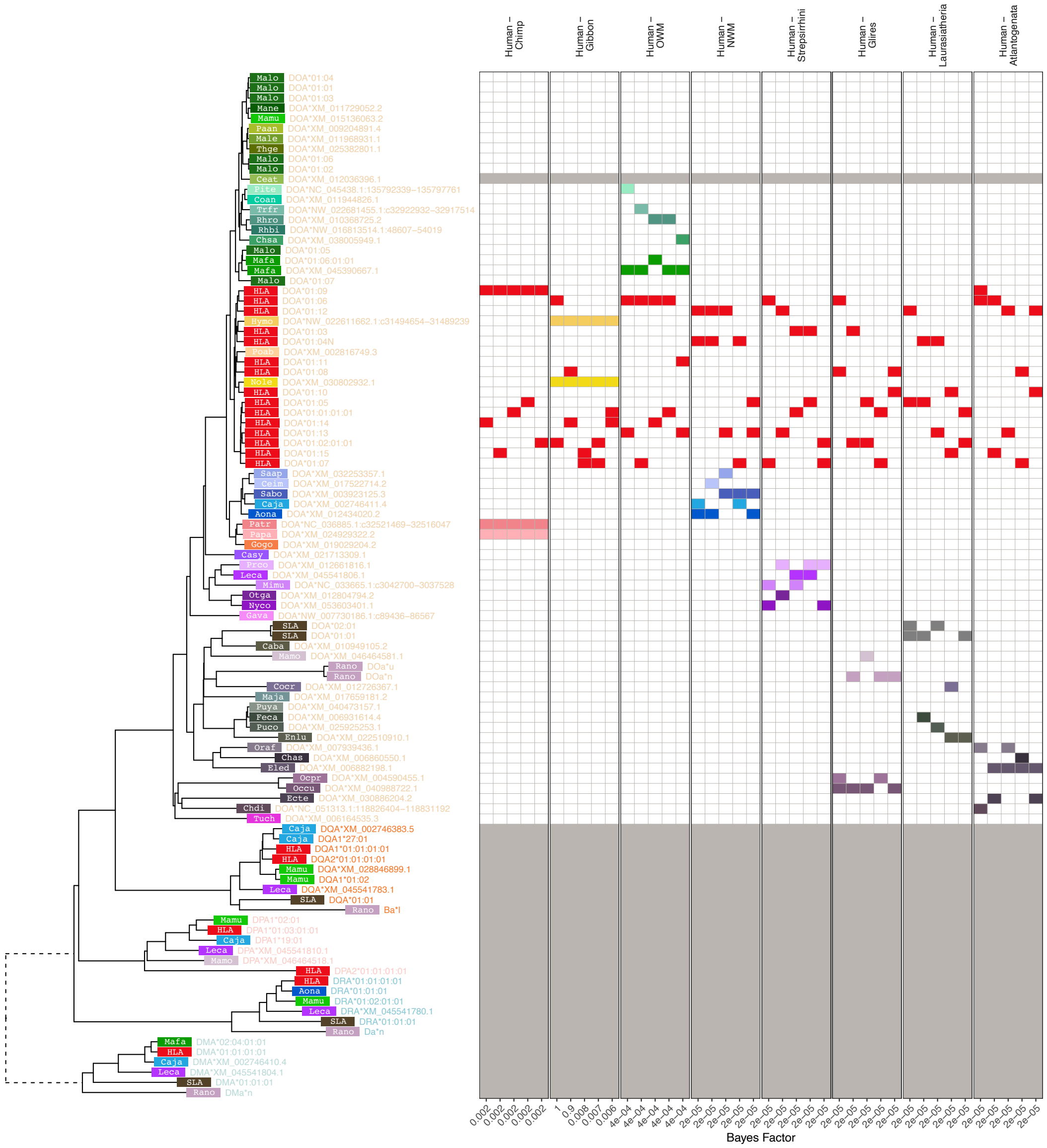

**Figure 4—figure supplement 5. MHC-DOA-Related Group *BEAST2* tree for exon 3 (PBR-encoding).** In the tree, each tip represents a sequence (see ***Appendix 1*** for more details on nomenclature), with the colored rectangle and four-letter abbreviation indicating the species (see ***Figure 1—figure Supplement 2*** for full species key). Following the rectangle, tips are labeled with the sequence name; sequences which have been assigned to loci are colored according to the gene group, while unassigned sequences are written in gray. Dashed branches are shortened to 10% of their length to expand detail in the rest of the tree. The left-hand side shows example Bayes factors we calculated from the set of posterior trees; the tree tips correspond to the rows of the grid. Each panel represents a type of comparison (labeled at top) and each column represents one of the top 5 highest-Bayes-factor comparisons (exact value at bottom; $> 100$ indicates strong evidence for TSP). The 4 colored blocks in each column include two red blocks (human sequences) that were tested against two sequences from other species (see Methods). Grayed-out rows correspond to sequences that were not considered for Bayes factors, either because they belong to a non-orthologous gene or backbone sequence (and thus not relevant to compare with the human gene in question), or because they may have been involved in a gene conversion event in this exon (according to *GENECONV* or from the literature).

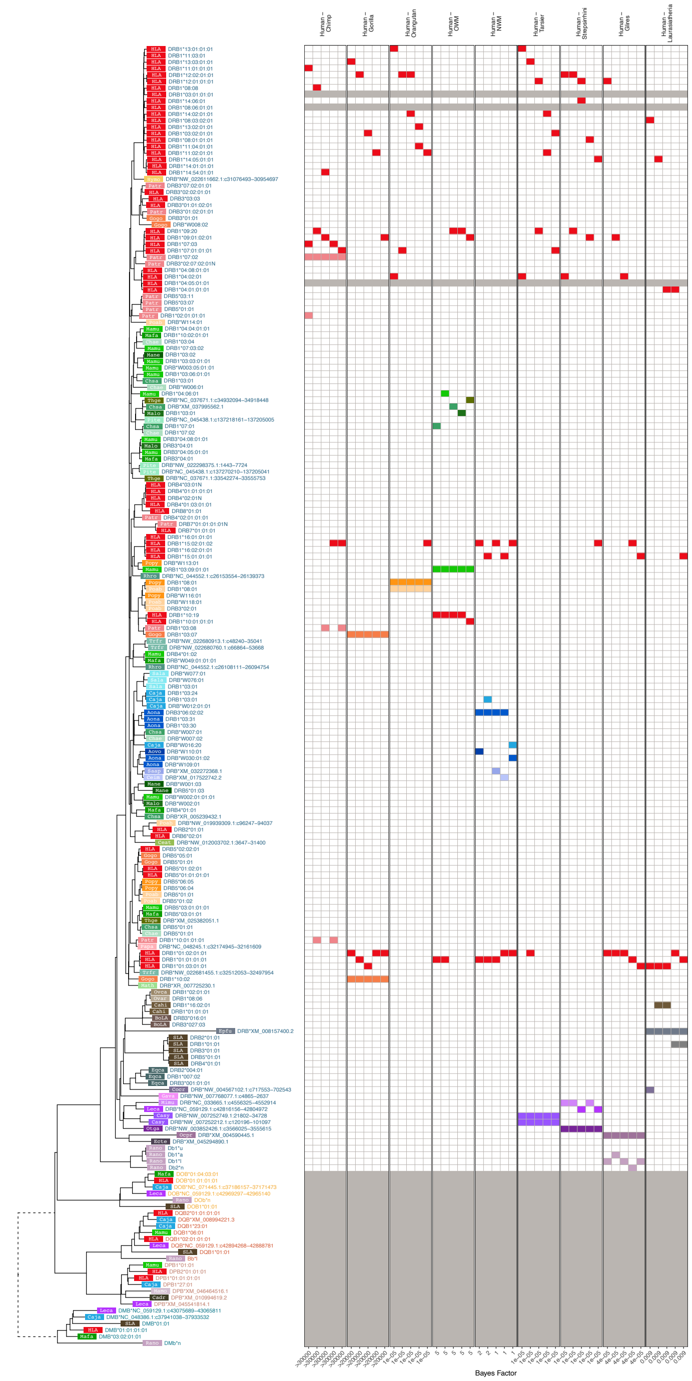


**Figure 4—figure supplement 6. MHC-DRB-Related Group *BEAST2* tree for exon 3 (PBR-encoding).** In the tree, each tip represents a sequence (see ***Appendix 1*** for more details on nomenclature), with the colored rectangle and four-letter abbreviation indicating the species (see ***Figure 1—figure Supplement 2*** for full species key). Following the rectangle, tips are labeled with the sequence name; sequences which have been assigned to loci are colored according to the gene group, while unassigned sequences are written in gray. Dashed branches are shortened to 10% of their length to expand detail in the rest of the tree. The left-hand side shows example Bayes factors we calculated from the set of posterior trees; the tree tips correspond to the rows of the grid. Each panel represents a type of comparison (labeled at top) and each column represents one of the top 5 highest-Bayes-factor comparisons (exact value at bottom; $> 100$ indicates strong evidence for TSP). The 4 colored blocks in each column include two red blocks (human sequences) that were tested against two sequences from other species (see Methods). Grayed-out rows correspond to sequences that were not considered for Bayes factors, either because they belong to a non-orthologous gene or backbone sequence (and thus not relevant to compare with the human gene in question), or because they may have been involved in a gene conversion event in this exon (according to *GENECONV* or from the literature).

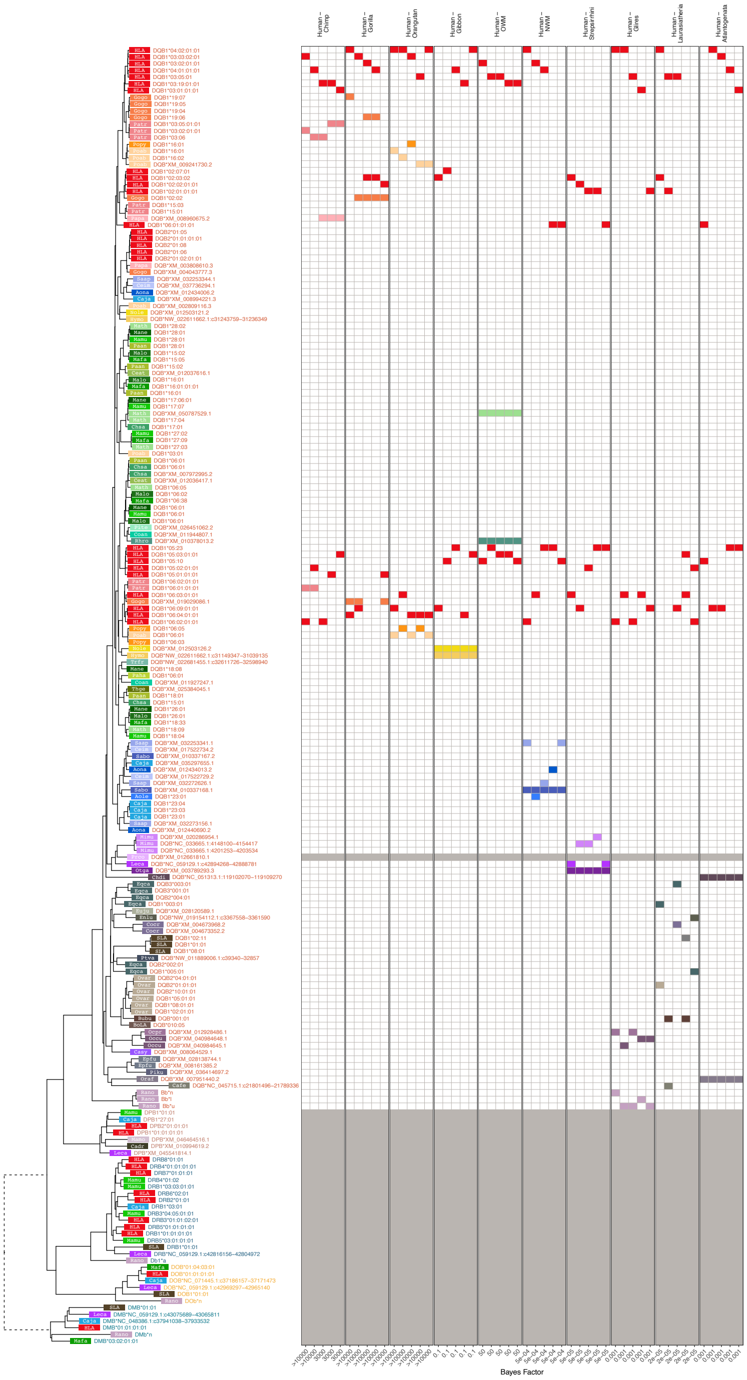


**Figure 4—figure supplement 7. MHC-DQB-Related Group *BEAST2* tree for exon 3 (PBR-encoding).** In the tree, each tip represents a sequence (see ***Appendix 1*** for more details on nomenclature), with the colored rectangle and four-letter abbreviation indicating the species (see ***Figure 1—figure Supplement 2*** for full species key). Following the rectangle, tips are labeled with the sequence name; sequences which have been assigned to loci are colored according to the gene group, while unassigned sequences are written in gray. Dashed branches are shortened to 10% of their length to expand detail in the rest of the tree. The left-hand side shows example Bayes factors we calculated from the set of posterior trees; the tree tips correspond to the rows of the grid. Each panel represents a type of comparison (labeled at top) and each column represents one of the top 5 highest-Bayes-factor comparisons (exact value at bottom; $> 100$ indicates strong evidence for TSP). The 4 colored blocks in each column include two red blocks (human sequences) that were tested against two sequences from other species (see Methods). Grayed-out rows correspond to sequences that were not considered for Bayes factors, either because they belong to a non-orthologous gene or backbone sequence (and thus not relevant to compare with the human gene in question), or because they may have been involved in a gene conversion event in this exon (according to *GENECONV* or from the literature).

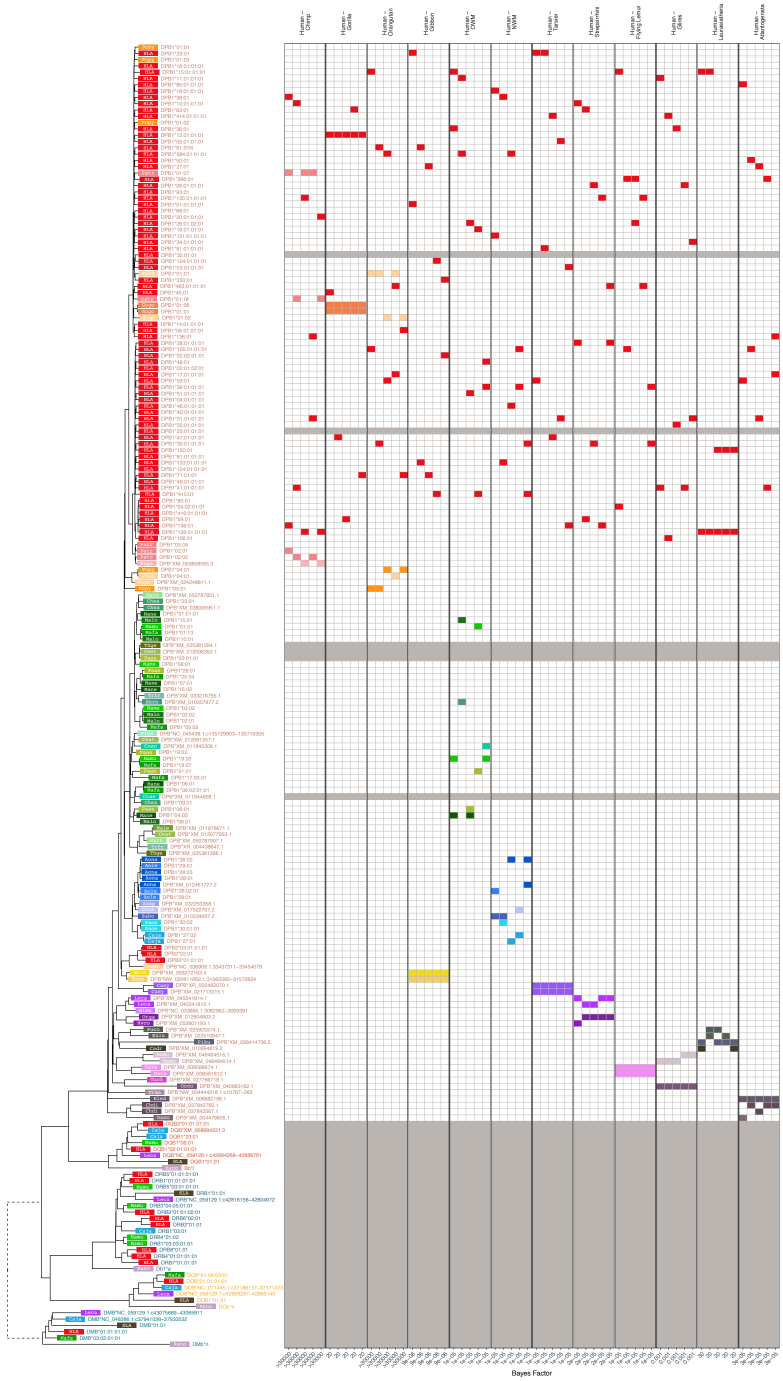


**Figure 4—figure supplement 8. MHC-DPB-Related Group *BEAST2* tree for exon 3 (PBR-encoding).** In the tree, each tip represents a sequence (see ***Appendix 1*** for more details on nomenclature), with the colored rectangle and four-letter abbreviation indicating the species (see ***Figure 1—figure Supplement 2*** for full species key). Following the rectangle, tips are labeled with the sequence name; sequences which have been assigned to loci are colored according to the gene group, while unassigned sequences are written in gray. Dashed branches are shortened to 10% of their length to expand detail in the rest of the tree. The left-hand side shows example Bayes factors we calculated from the set of posterior trees; the tree tips correspond to the rows of the grid. Each panel represents a type of comparison (labeled at top) and each column represents one of the top 5 highest-Bayes-factor comparisons (exact value at bottom; $> 100$ indicates strong evidence for TSP). The 4 colored blocks in each column include two red blocks (human sequences) that were tested against two sequences from other species (see Methods). Grayed-out rows correspond to sequences that were not considered for Bayes factors, either because they belong to a non-orthologous gene or backbone sequence (and thus not relevant to compare with the human gene in question), or because they may have been involved in a gene conversion event in this exon (according to *GENECONV* or from the literature).

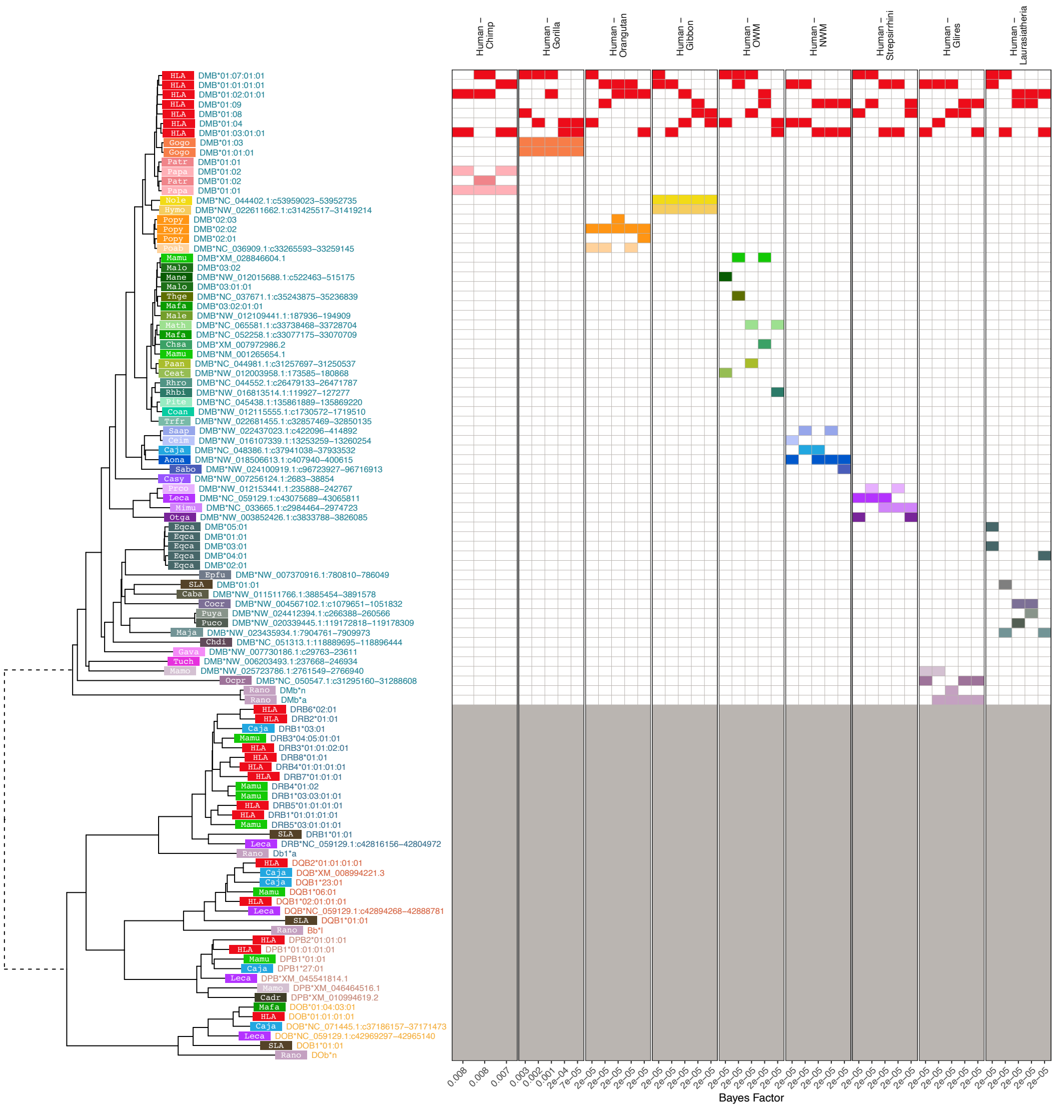


**Figure 4—figure supplement 9. MHC-DMB-Related Group *BEAST2* tree for exon 3 (PBR-encoding).** In the tree, each tip represents a sequence (see ***Appendix 1*** for more details on nomenclature), with the colored rectangle and four-letter abbreviation indicating the species (see ***Figure 1—figure Supplement 2*** for full species key). Following the rectangle, tips are labeled with the sequence name; sequences which have been assigned to loci are colored according to the gene group, while unassigned sequences are written in gray. Dashed branches are shortened to 10% of their length to expand detail in the rest of the tree. The left-hand side shows example Bayes factors we calculated from the set of posterior trees; the tree tips correspond to the rows of the grid. Each panel represents a type of comparison (labeled at top) and each column represents one of the top 5 highest-Bayes-factor comparisons (exact value at bottom; $> 100$ indicates strong evidence for TSP). The 4 colored blocks in each column include two red blocks (human sequences) that were tested against two sequences from other species (see Methods). Grayed-out rows correspond to sequences that were not considered for Bayes factors, either because they belong to a non-orthologous gene or backbone sequence (and thus not relevant to compare with the human gene in question), or because they may have been involved in a gene conversion event in this exon (according to *GENECONV* or from the literature).

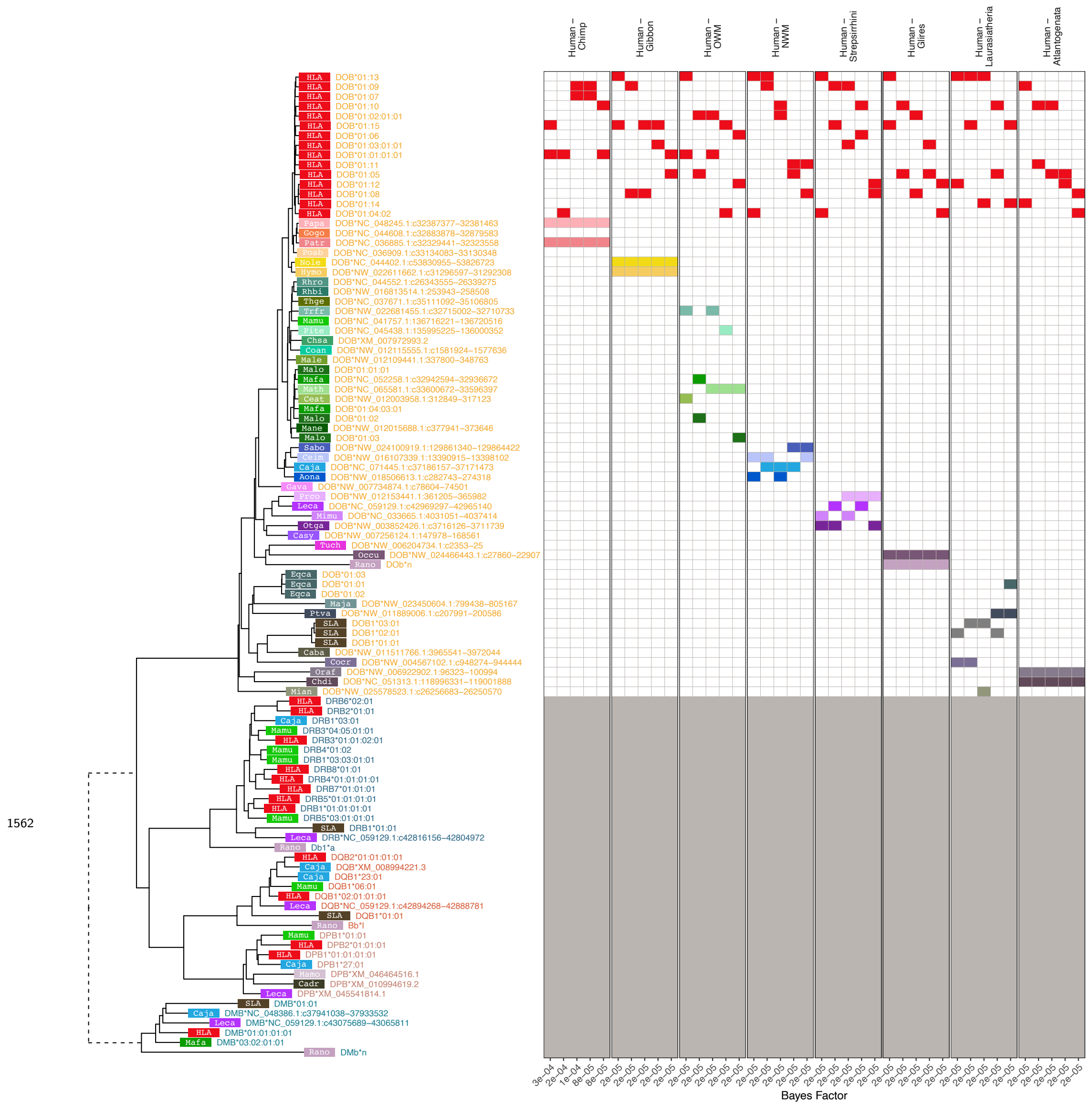


**Figure 4—figure supplement 10. MHC-DOB-Related Group *BEAST2* tree for exon 3 (PBR-encoding).** In the tree, each tip represents a sequence (see ***Appendix 1*** for more details on nomenclature), with the colored rectangle and four-letter abbreviation indicating the species (see ***Figure 1—figure Supplement 2*** for full species key). Following the rectangle, tips are labeled with the sequence name; sequences which have been assigned to loci are colored according to the gene group, while unassigned sequences are written in gray. Dashed branches are shortened to 10% of their length to expand detail in the rest of the tree. The left-hand side shows example Bayes factors we calculated from the set of posterior trees; the tree tips correspond to the rows of the grid. Each panel represents a type of comparison (labeled at top) and each column represents one of the top 5 highest-Bayes-factor comparisons (exact value at bottom; $> 100$ indicates strong evidence for TSP). The 4 colored blocks in each column include two red blocks (human sequences) that were tested against two sequences from other species (see Methods). Grayed-out rows correspond to sequences that were not considered for Bayes factors, either because they belong to a non-orthologous gene or backbone sequence (and thus not relevant to compare with the human gene in question), or because they may have been involved in a gene conversion event in this exon (according to *GENECONV* or from the literature).

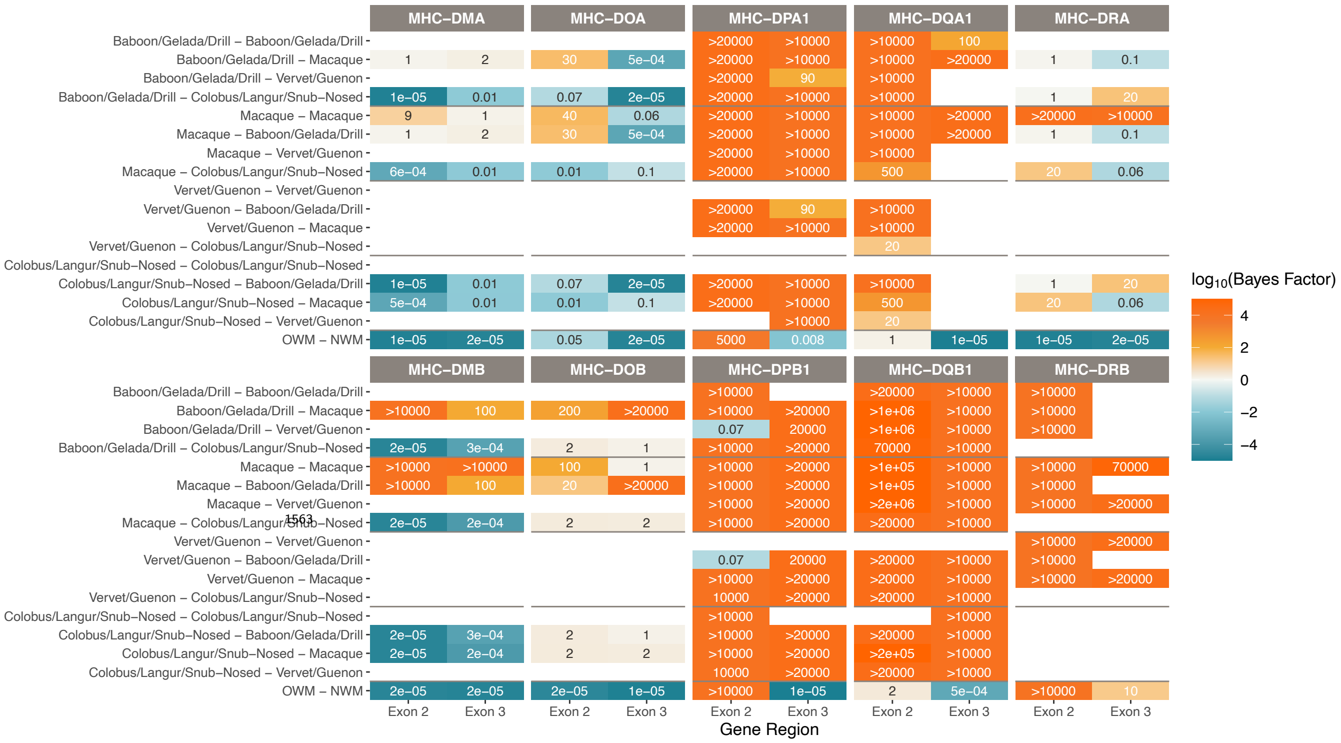


**Figure 4—figure supplement 11. TSP among old-world monkey groups for the Class II genes.** Different species comparisons are listed on the y-axis, and different gene regions are listed on the x-axis. Each table entry is colored and labeled with the maximum Bayes factor among all tested quartets of alleles belonging to that category. High Bayes factors (orange) indicate support for TSP among the given species for that gene region, while low Bayes factors (teal) indicate that alleles assort according to the species tree, as expected. Bayes factors above 100 are considered decisive. Tan values show poor support for either hypothesis, while white boxes indicate that there are not enough alleles in that category with which to calculate Bayes factors.

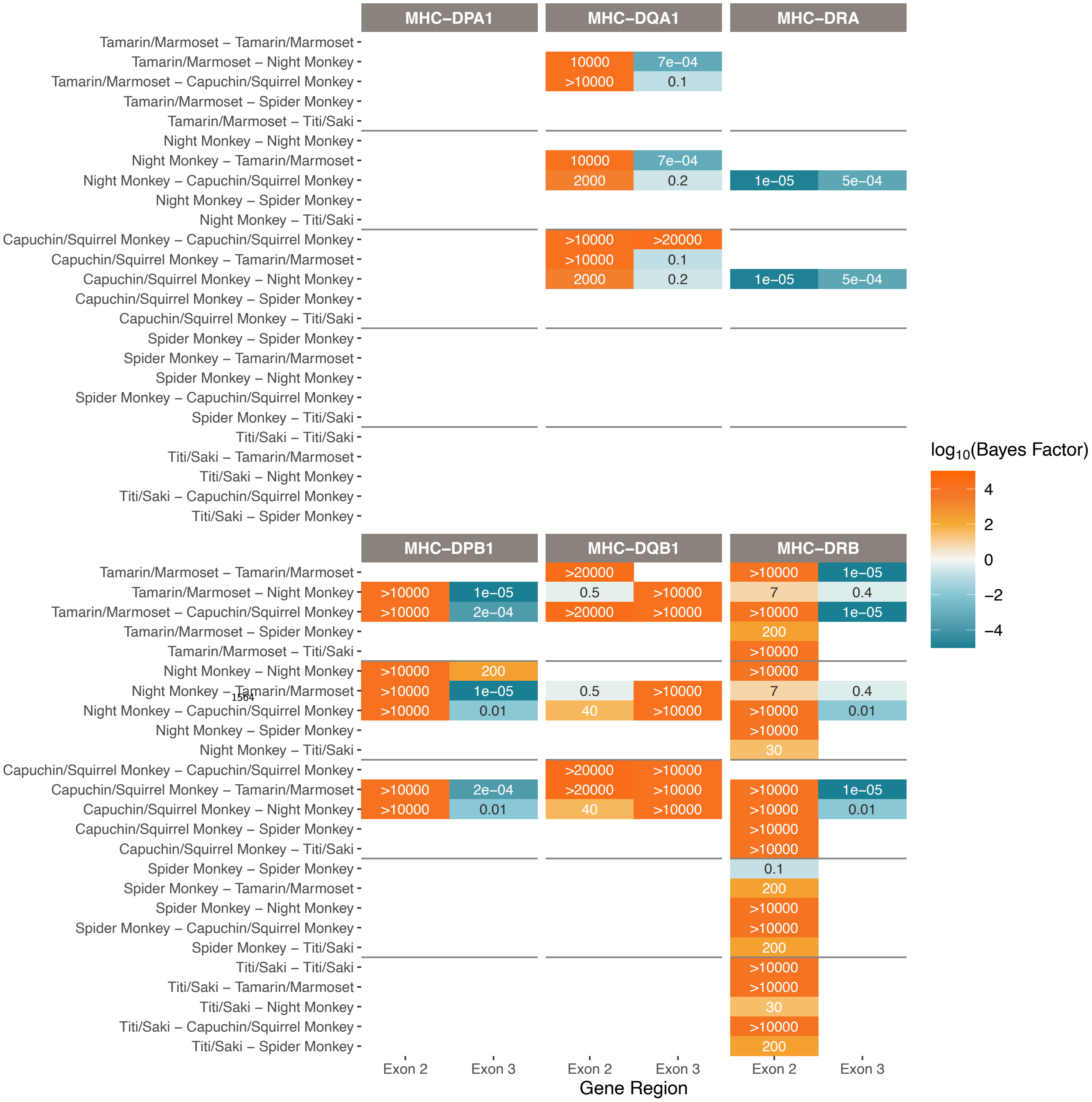


**Figure 4—figure supplement 12. TSP among new-world monkey groups for the Class II genes.** Different species comparisons are listed on the y-axis, and different gene regions are listed on the x-axis. Each table entry is colored and labeled with the maximum Bayes factor among all tested quartets of alleles belonging to that category. High Bayes factors (orange) indicate support for TSP among the given species for that gene region, while low Bayes factors (teal) indicate that alleles assort according to the species tree, as expected. Bayes factors above 100 are considered decisive. Tan values show poor support for either hypothesis, while white boxes indicate that there are not enough alleles in that category with which to calculate Bayes factors. There is not enough data in each category to compute Bayes factors for these groups for MHC-DPA1, -DMA, -DMB, -DOA, and -DOB.

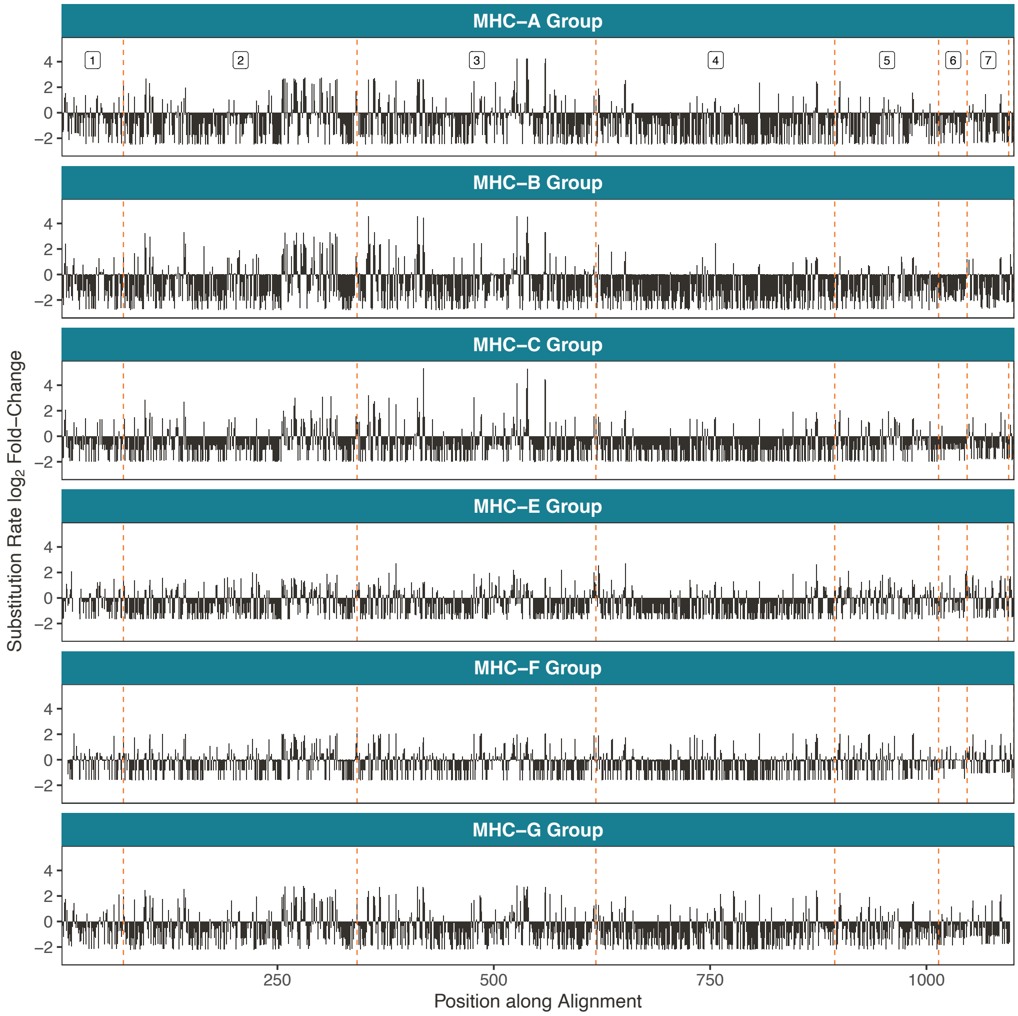


**Figure 5—figure supplement 1. Rapidly-evolving sites in the Class I genes.** Rapidly-evolving sites are primarily located in exons 2 and 3. Here, the exons are concatenated such that the cumulative position along the coding region is on the x-axis. The dashed orange lines denote exon boundaries. The genes are aligned such that the same vertical position indicates an evolutionarily equivalent site. The y-axis shows the substitution rate at each site, expressed as a fold-change (the base-2 logarithm of each site's evolutionary rate divided by the mean rate among mostly-gap sites in each alignment; see Methods).

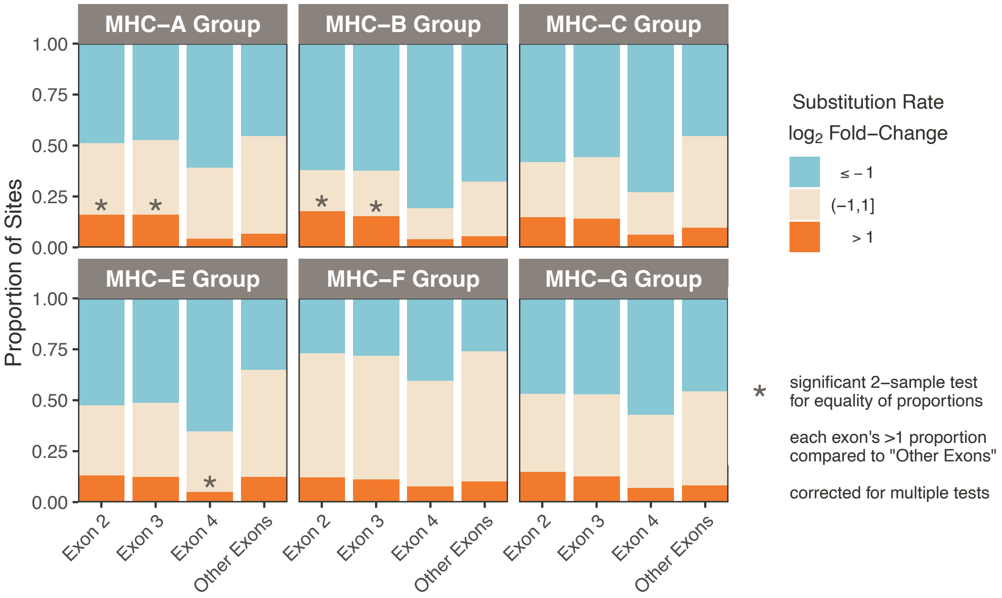


**Figure 5—figure supplement 2. Proportions of rapidly-evolving sites for Class I.** Sites were binned into slowly-evolving (≤-1), rapidly-evolving (>1), or baseline (>-1 but ≤1) categories. We then calculated proportions of these categories for exon 2, exon 3, exon 4, and the "other" exons (not exons 2, 3, or 4). These bins are all approximately the same size, $\sim 270$bp. For each gene and exon, we tested the difference in the proportion of rapidly-evolving sites between that exon and the "other" exons group (2-sample z-test for equality of proportions with continuity correction). Significant tests (Bonferroni corrected; p<0.05×3) are marked with an asterisk.

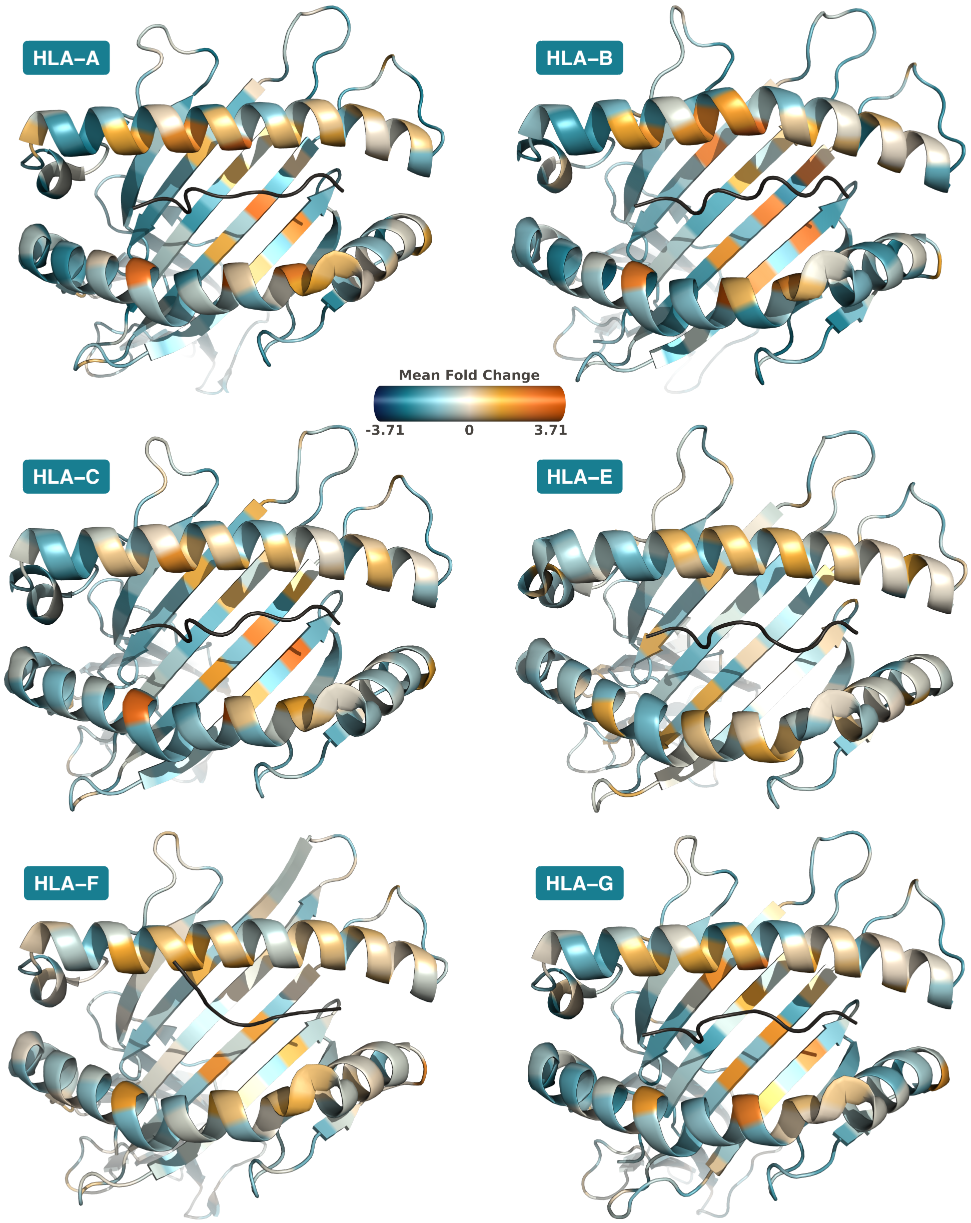


**Figure 5—figure supplement 3. Rapidly-Evolving Sites on Class I Protein Structures.** Structures are Protein Data Bank (***Berman et al., 2000***) 6J1W (***Zhu et al., 2019***) for HLA-A, 3BVN (***Kumar et al., 2009***) for HLA-B, 4NT6 (***Choo et al., 2014***) for HLA-C, 7P4B (***Walters et al., 2022***) for HLA-E, 5IUE (***Dulberger et al., 2017***) for HLA-F, and 3KYN (***Walpole et al., 2010***) for HLA-G, with images created in *PyMOL* (***Sch, 2021***). Substitution rates for each amino acid are computed as the mean substitution rate of the three sites composing the codon. Orange indicates rapidly-evolving amino acids, while teal indicates conserved amino acids.

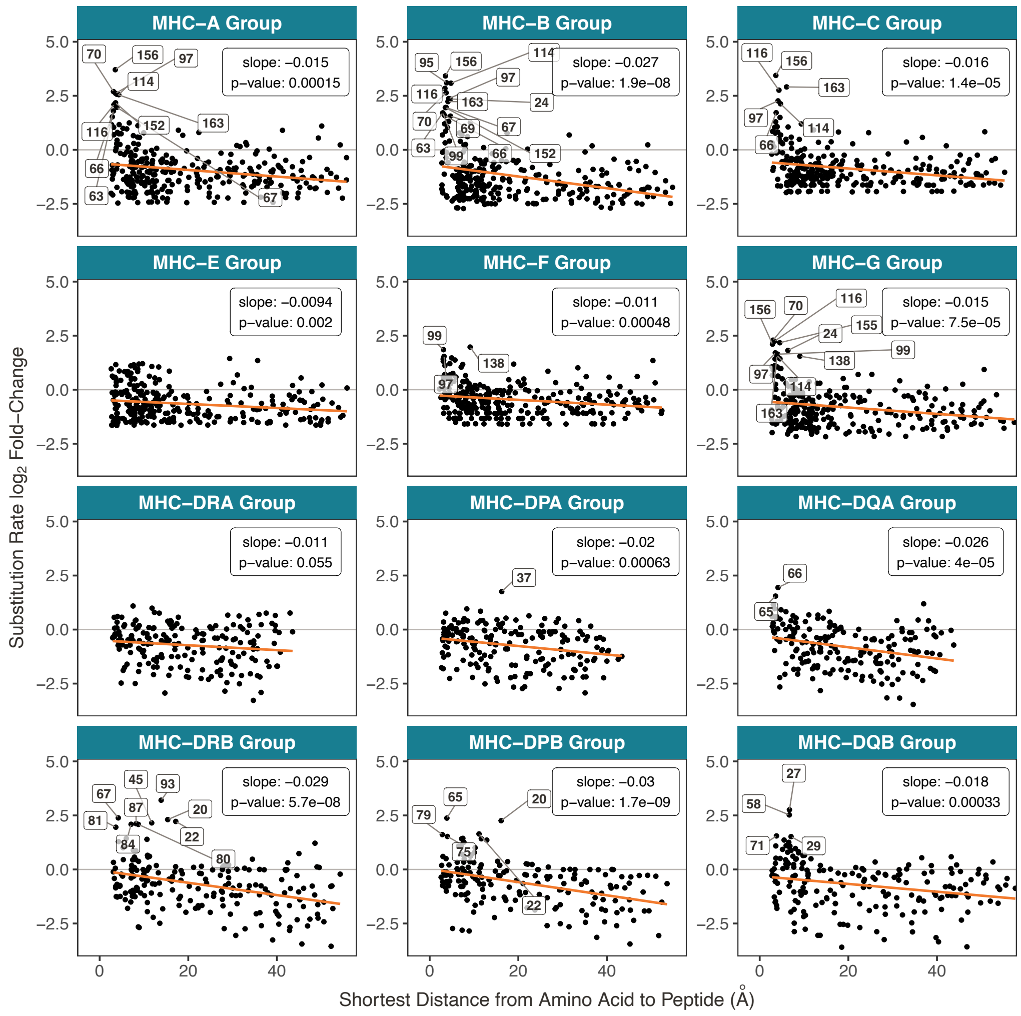


**Figure 5—figure supplement 4. Evolutionary rate is related to the distance to peptide.** The y-axis shows the *BEAST2* substitution rate, expressed as a fold-change (the base-2 logarithm of each site's evolutionary rate divided by the mean rate among mostly-gap sites in each alignment; see Methods). The x-axis shows the minimum distance to the bound peptide, measured in *PyMOL* (***Sch, 2021***). Each point is an amino acid, and distances are averaged over several structures (see ***Table 5***). The orange line is a linear regression of substitution rate on minimum distance, with slope and p-value annotated on each panel. Amino acids with a fold-change greater than 1.5 are labeled.

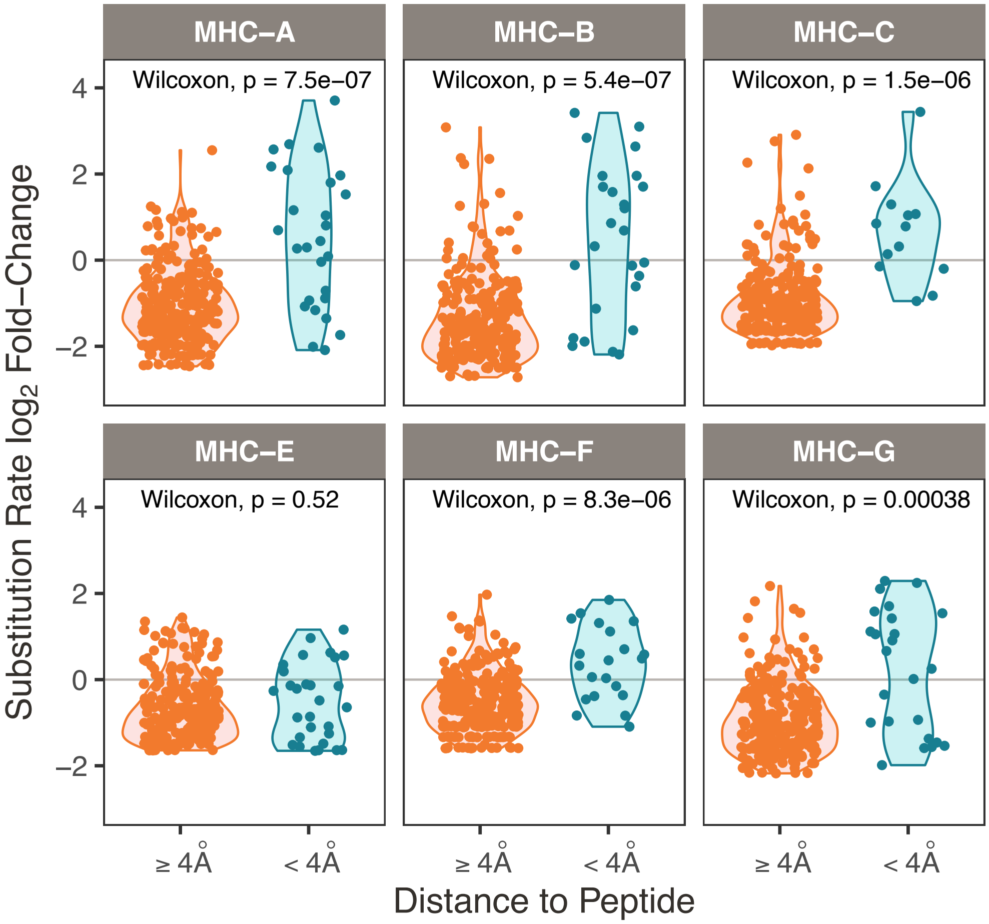


**Figure 5—figure supplement 5. Class I rapidly-evolving sites by binned distance to peptide.** Nucleotide sites were divided into those whose corresponding amino acids contact the peptide (< 4Å) versus do not contact the peptide (≥ 4Å), shown on the x-axis. This distance has been used previously to define plausible peptide-contacting residues (***Nielsen et al., 2007***). The y-axis shows the substitution rate at each site, expressed as a fold-change (the base-2 logarithm of each site's evolutionary rate divided by the mean rate among mostly-gap sites in each alignment; see Methods). For each gene, the groups are compared using a Wilcoxon test, with p-value displayed at the top of each panel.

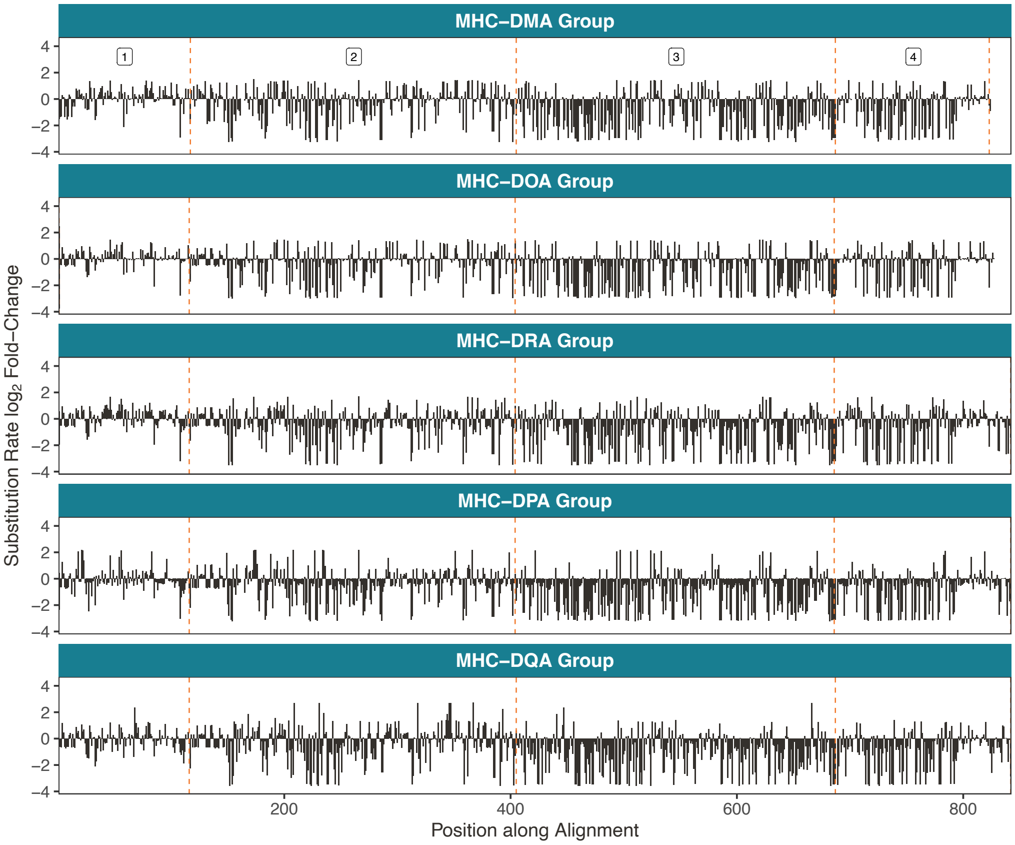


**Figure 6—figure supplement 1. Rapidly-evolving sites in the Class IIA genes. A)** Rapidly-evolving sites are primarily located in exon 2. Here, the exons are concatenated such that the cumulative position along the coding region is on the x-axis. The dashed orange lines denote exon boundaries. The genes are aligned such that the same vertical position indicates an evolutionarily equivalent site. The y-axis shows the substitution rate at each site, expressed as a fold-change (the base-2 logarithm of each site's evolutionary rate divided by the mean rate among mostly-gap sites in each alignment; see Methods).

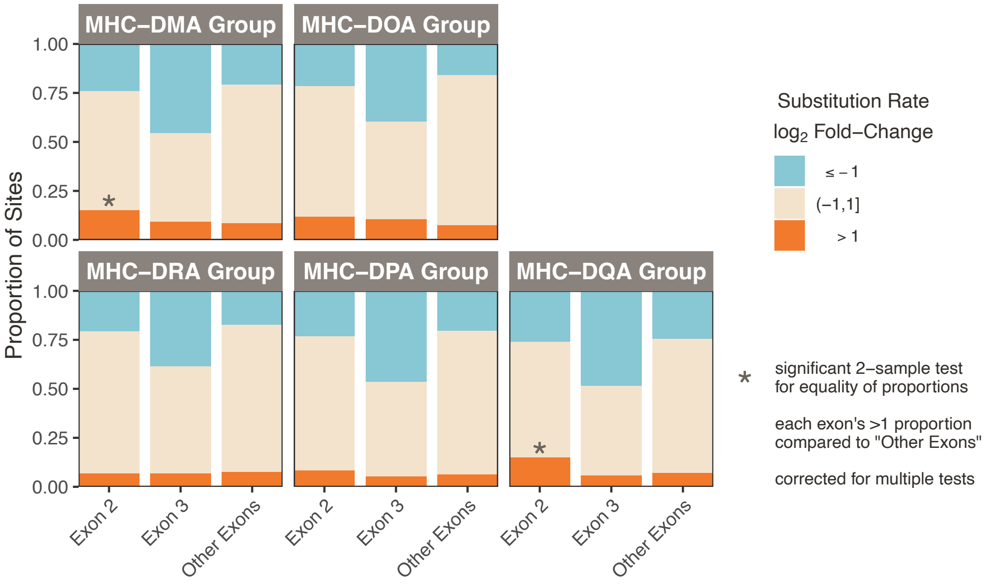


**Figure 6—figure supplement 2. Proportions of rapidly-evolving sites for Class IIA.** Sites were binned into slowly-evolving (≤-1), rapidly-evolving (>1), or baseline (>-1 but ≤1) categories. We then calculated proportions of these categories for exon 2, exon 3, and the "other" exons (not exons 2 or 3). These bins are all approximately the same size, $\sim 270$bp. For each gene and exon, we tested the difference in the proportion of rapidly-evolving sites between that exon and the "other" exons group (2-sample z-test for equality of proportions with continuity correction). Significant tests (Bonferroni corrected; p<0.05×2) are marked with an asterisk.

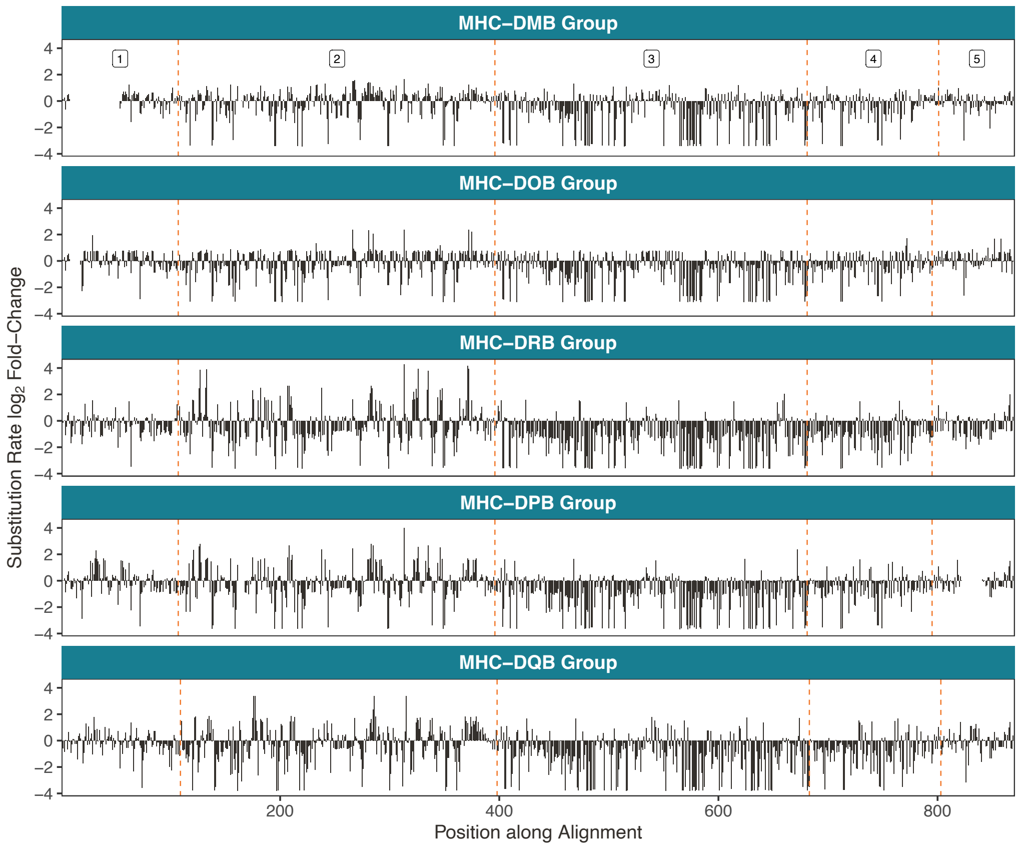


**Figure 6—figure supplement 3. Rapidly-evolving sites in the Class IIB genes. A)** Rapidly-evolving sites are primarily located in exon 2. Here, the exons are concatenated such that the cumulative position along the coding region is on the x-axis. The dashed orange lines denote exon boundaries. The genes are aligned such that the same vertical position indicates an evolutionarily equivalent site. The y-axis shows the substitution rate at each site, expressed as a fold-change (the base-2 logarithm of each site's evolutionary rate divided by the mean rate among mostly-gap sites in each alignment; see Methods).

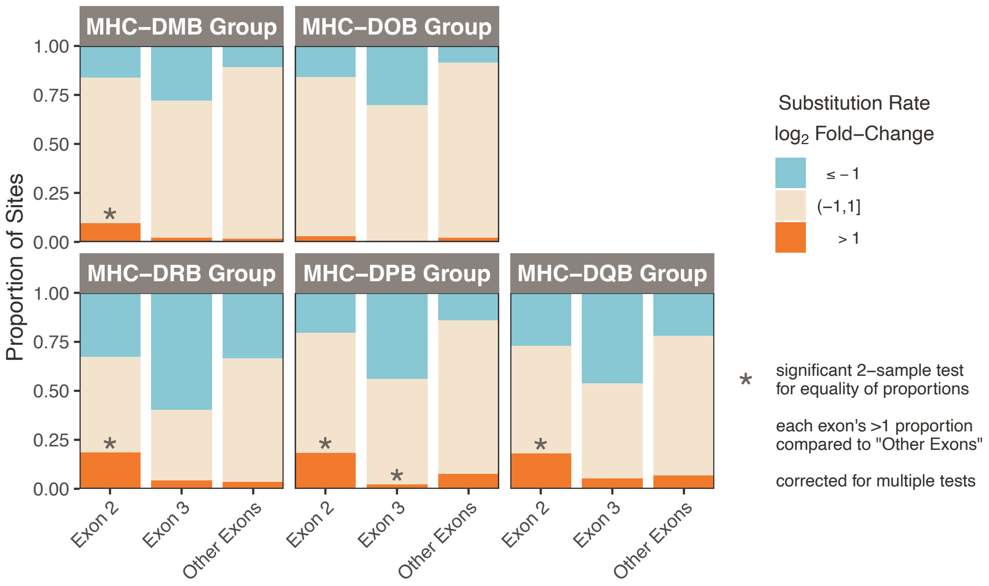


**Figure 6—figure supplement 4. Proportions of rapidly-evolving sites for Class IIB.** Sites were binned into slowly-evolving (≤-1), rapidly-evolving (>1), or baseline (>-1 but ≤1) categories. We then calculated proportions of these categories for exon 2, exon 3, and the "other" exons (not exons 2 or 3). These bins are all approximately the same size, $\sim 270$bp. For each gene and exon, we tested the difference in the proportion of rapidly-evolving sites between that exon and the "other" exons group (2-sample z-test for equality of proportions with continuity correction). Significant tests (Bonferroni corrected; $p<0.05\times2$) are marked with an asterisk.

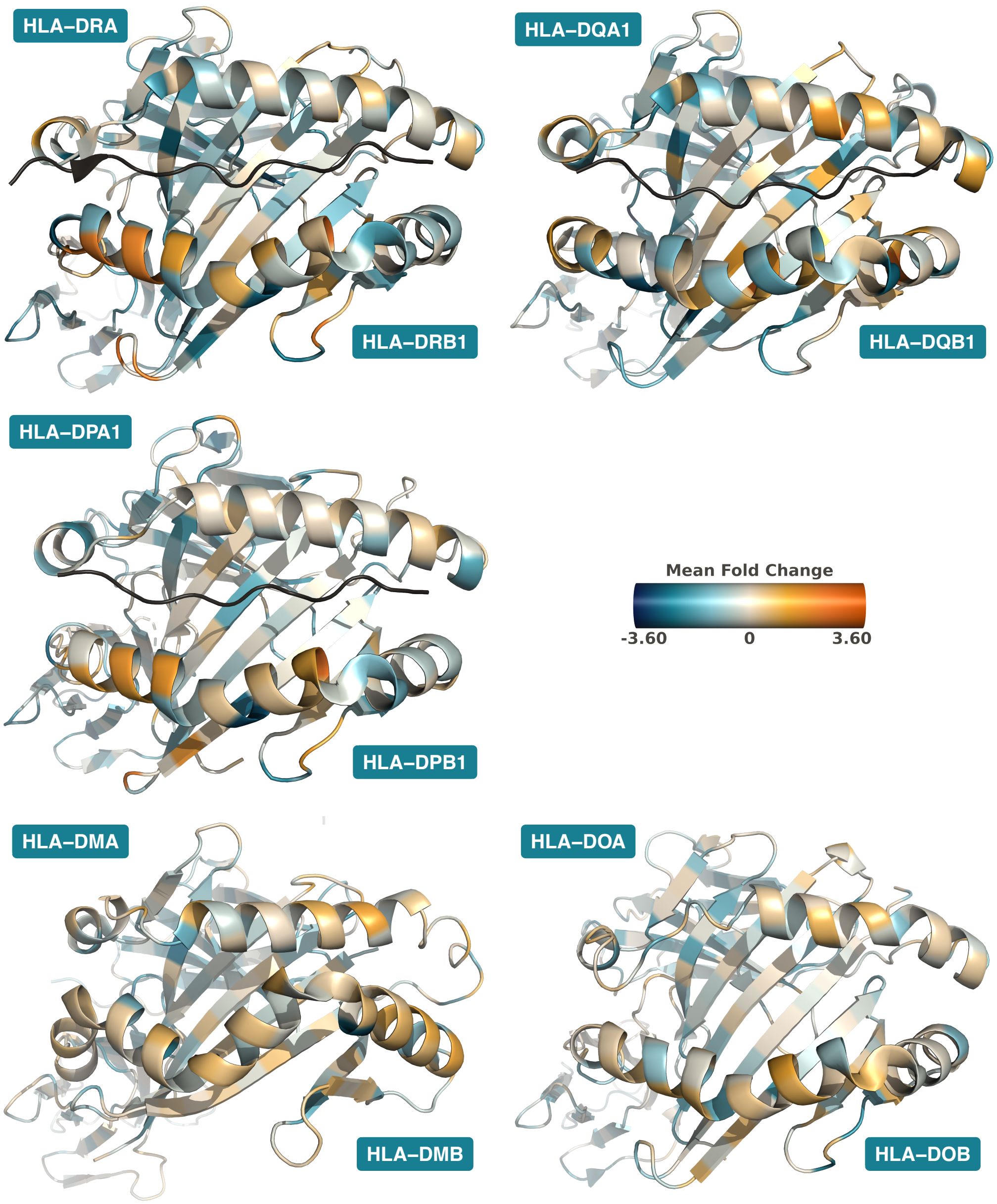


**Figure 6—figure supplement 5. Rapidly-Evolving Sites on Class II Protein Structures.** Structures are Protein Data Bank (***Berman et al., 2000***) 5JLZ (***Gerstner et al., 2016***) for HLA-DR, 2NNA (***Henderson et al., 2007***) for HLA-DQ, 7T2A (***Ciacchi et al., 2023***) for HLA-DP, 2BC4 (***Nicholson et al., 2006***) for HLA-DM, and 4I0P (***Guce et al., 2013***) for HLA-DO, with images created in *PyMOL* (***Sch, 2021***). Substitution rates for each amino acid are computed as the mean substitution rate of the three sites composing the codon. Orange indicates rapidly-evolving amino acids, while teal indicates conserved amino acids.

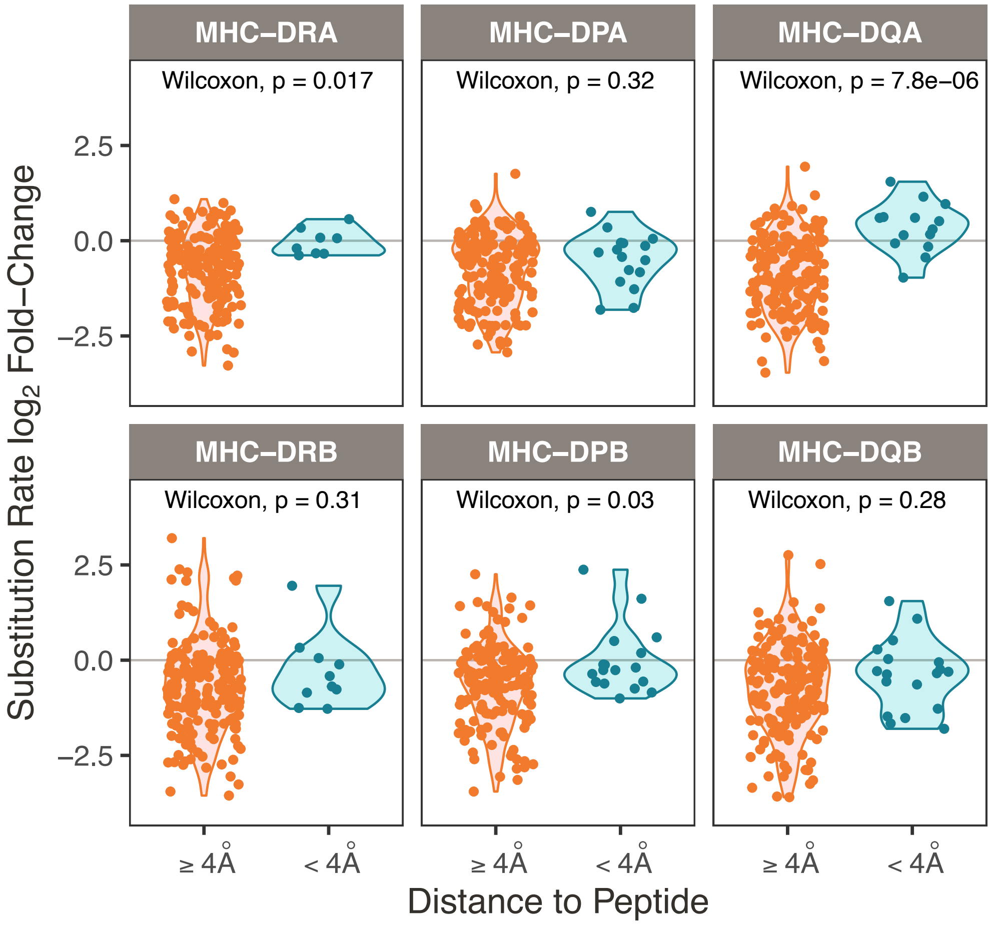


**Figure 6—figure supplement 6. Class II rapidly-evolving sites by binned distance to peptide.** Nucleotide sites were divided into those whose corresponding amino acids contact the peptide (< 4Å) versus do not contact the peptide (≥ 4Å), shown on the x-axis. This distance has been used previously to define plausible peptide-contacting residues (***Nielsen et al., 2007***). The y-axis shows the substitution rate at each site, expressed as a fold-change (the base-2 logarithm of each site's evolutionary rate divided by the mean rate among mostly-gap sites in each alignment; see Methods). For each gene, the groups are compared using a Wilcoxon test, with p-value displayed at the top of each panel. MHC-DM and -DO are not shown because they do not bind peptides.

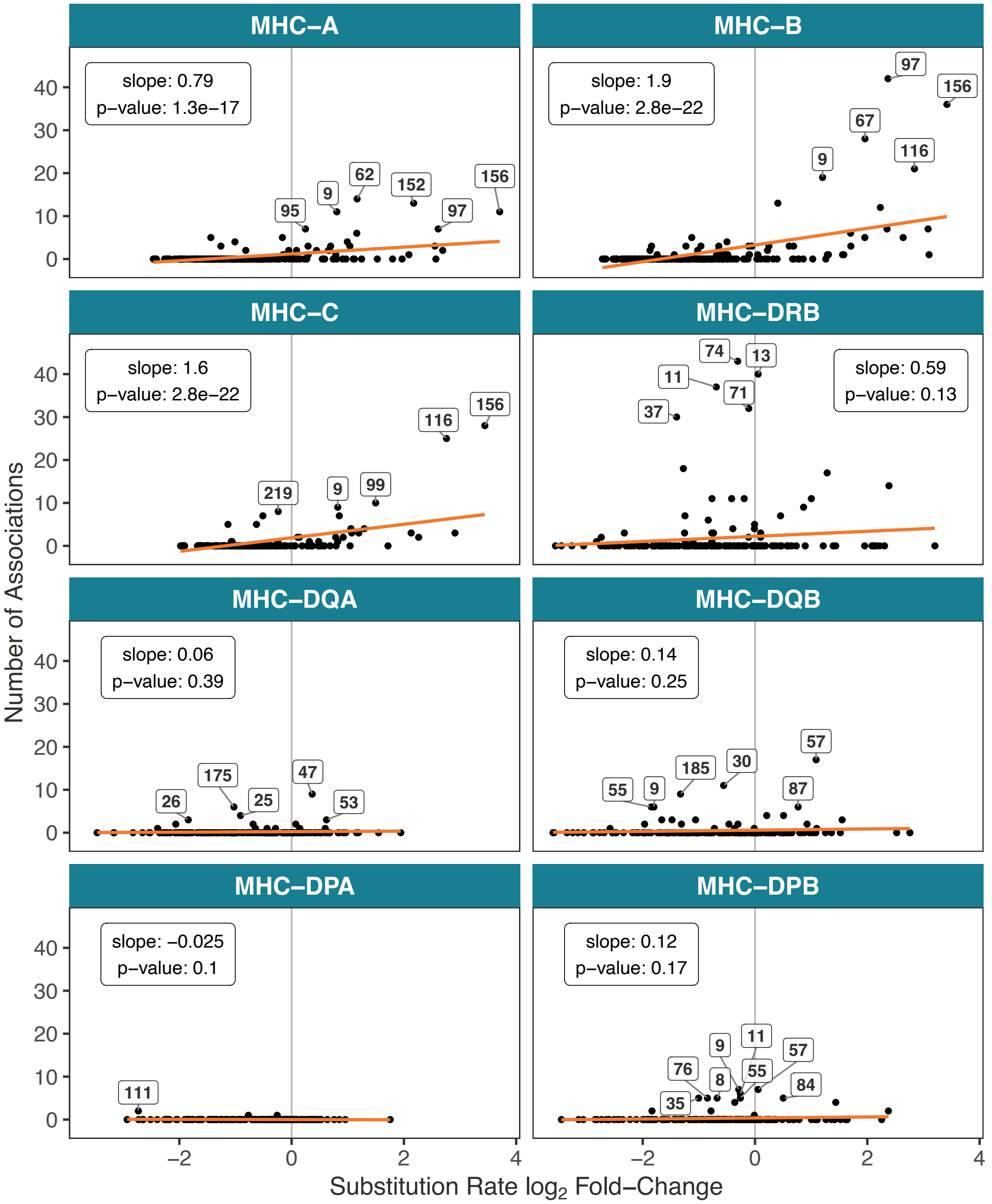


**Figure 6—figure supplement 7. Number of associations per amino acid as a function of evolutionary rate.** The x-axis shows the substitution rate at each site, expressed as a fold-change (the base-2 logarithm of each site's evolutionary rate divided by the mean rate among mostly-gap sites in each alignment; see Methods). The y-axis shows the number of unique associations for each amino acid, including diseases, TCR phenotypes, and protein expression levels. Only genes with associations are shown; at the time of publishing, there were no amino acid associations meeting our criteria for MHC-E, -F, -G, -DRA, -DMA, -DMB, -DOA, or -DOB. For each gene, a regression line is shown in orange, with slope and p-value displayed at the top of each panel. The amino acids with the greatest number of associations within each gene are also labeled.
